# Improved ribosome-footprint and mRNA measurements provide insights into dynamics and regulation of yeast translation

**DOI:** 10.1101/021501

**Authors:** David E. Weinberg, Premal Shah, Stephen W. Eichhorn, Jeffrey A. Hussmann, Joshua B. Plotkin, David P. Bartel

## Abstract

Ribosome-footprint profiling provides genome-wide snapshots of translation, but technical challenges can confound its analysis. Here, we use improved methods to obtain ribosome-footprint profiles and mRNA abundances that more faithfully reflect gene expression in *Saccharomyces cerevisiae*. Our results support proposals that both the beginning of coding regions and codons matching rare tRNAs are more slowly translated. They also indicate that emergent polypeptides with as few as three basic residues within a 10-residue window tend to slow translation. With the improved mRNA measurements, the variation attributable to translational control in exponentially growing yeast was less than previously reported, and most of this variation could be predicted with a simple model that considered mRNA abundance, upstream open reading frames, cap-proximal structure and nucleotide composition, and lengths of the coding and 5’-untranslated regions. Collectively, our results reveal key features of translational control in yeast and provide a framework for executing and interpreting ribosome-profiling studies.

## Introduction

The central dogma of molecular biology culminates in translation. During translation initiation, the small ribosomal subunit is recruited to the mRNA substrate, a suitable start codon is identified, and then the large ribosomal subunit joins to form a functional, elongation-competent ribosome. The ribosome then translocates along the mRNA, synthesizing the cognate protein, until reaching a stop codon, where release factors mediate release of the polypeptide, and recycling factors then dissociate the translation machinery.

Although most cellular mRNAs use the same translation machinery, the dynamics of translation can vary between mRNAs and within mRNAs, sometimes with regulatory or functional consequences. For example, strong secondary structure within the 5′ untranslated region (UTR) of a eukaryotic mRNA can impede the scanning ribosome, thereby reducing the rate of protein synthesis (Kozak, 1986a). The accessibility of the 5′ cap, the presence of small ORFs within 5′ UTRs referred to as upstream ORFs (uORFs), and the sequence context of the start codon of the ORF can also modulate the rate of translation initiation (Kozak, 1984; Godefroy-Colburn et al., 1985; Kozak, 1986b). Likewise, codon choice, mRNA structure, and the identity of the nascent polypeptide can influence elongation rates (Varenne et al., 1984; Hosoda et al., 2003; Brandman et al., 2012). In addition, differences in elongation rates can influence co-translational protein folding, localization of the mRNA or protein, and in extreme cases the rate of protein production (Crombie et al., 1992; Letzring et al., 2010; Zhang and Shan, 2012). Finally, although translation termination is generally quite efficient, stop-codon read-through can introduce alternative C-terminal regions that affect protein stability, localization, or activity (Dunn et al., 2013).

Variation in protein abundances observed in eukaryotic cells largely reflects variation in mRNA abundances, indicating that much of gene regulation occurs at the level of mRNA synthesis and decay (Csárdi et al. 2015). However, differences in translation rate also contribute to the regulation of eukaryotic protein abundances. Despite known examples of regulation at each stage of translation, translational regulation is largely controlled at the step of initiation, as expected when considering that this step is rate limiting for most mRNAs (Bulmer, 1991; Shah et al., 2013).

The first demonstration that different mRNAs can be translated at different rates within the same cells was for the alpha and beta subunits of hemoglobin in rabbit reticulocytes. Despite the subunits being encoded on mRNAs of similar size, the beta-subunit mRNA is found on larger polysomes than the alpha-subunit mRNA (Hunt et al., 1968). This principle was later expanded to the genome-wide level with the use of microarrays to analyze the polysome profiles of thousands of mRNAs at once (Arava et al., 2003). Such studies in *Saccharomyces cerevisiae* suggested that ribosome densities vary among mRNAs over a 100-fold range (from 0.03 to 3.3 ribosomes per 100 nucleotides), indicating extensive translation control. More recently, the use of ribosome-footprint profiling has enabled transcriptome-wide analyses of translation using high-throughput sequencing, which again suggested a nearly 100-fold range of translational efficiencies (TEs) in log-phase yeast (Ingolia et al., 2009). Here, we generate and analyze substantially improved transcriptome-wide datasets that yield new insights into the dynamics and regulation of translation in yeast. These improved data also constrict the differences in TEs observed in log-phase yeast, which can be largely predicted using a simple model that considers only six features of the mRNAs.

## Results

### Native ribosome footprints reveal the dynamics of elongation

Protocols for analyzing polysome profiles or capturing ribosome footprints (referred to as ribosome-protected fragments, or RPFs) typically involve treating yeast cultures with the elongation inhibitor cycloheximide (CHX) to arrest the ribosomes prior to harvesting cells (Ingolia et al., 2009; Gerashchenko et al., 2012; Zinshteyn and Gilbert, 2013; Artieri and Fraser, 2014b; McManus et al., 2014). An advantage of CHX pre-treatment is that it prevents the run-off of ribosomes that can otherwise occur during the harvesting procedure. However, this treatment can also have some undesirable effects. Because CHX does not inhibit translation initiation or termination, pre-treatment of cultures leads to an accumulation of ribosomes at start codons and depletion of ribosomes at stop codons (Ingolia et al., 2011; Ingolia et al., 2012; Guydosh and Green, 2014). In addition, because CHX binding to the 80*S* ribosome is both non-instantaneous and reversible, the kinetics of CHX binding and dissociation might allow newly initiated ribosomes to translocate beyond the start codon. Another possible effect of CHX treatment is that ribosomes may preferentially arrest at specific codons that do not necessarily correspond to codons that are more abundantly occupied by ribosomes in untreated cells. Although these effects have minimal consequence for analyses at the mRNA level when comparing the mRNAs from the same gene in different conditions (e.g., Guo et al., 2010; Brar et al., 2012; Hsieh et al., 2012; Thoreen et al., 2012), or when comparing mRNAs from different genes after discarding reads corresponding to the 5′ regions of ORFs (e.g., Subtelny et al., 2014), these effects of CHX pre-treatment could have severe consequences for analyses that require single-codon resolution. With this in mind, some recent studies have used alternative methods to arrest ribosomes (Guydosh and Green, 2014; Lareau et al., 2014).

To avoid the confounding effects of CHX pre-treatment, we employed a protocol to rapidly harvest yeast cultures using filtration and flash freezing (Figure 1A). Importantly, our protocol minimizes the time that the cells experience starvation conditions, which lead to rapid ribosome run-off (Ashe et al., 2000; Guydosh and Green, 2014). CHX was included in the lysis buffer to inhibit translation elongation that might occur in lysates, perhaps during the room-temperature incubation used for RNase digestion, although we doubt this precaution was necessary.

**Figure 1:**
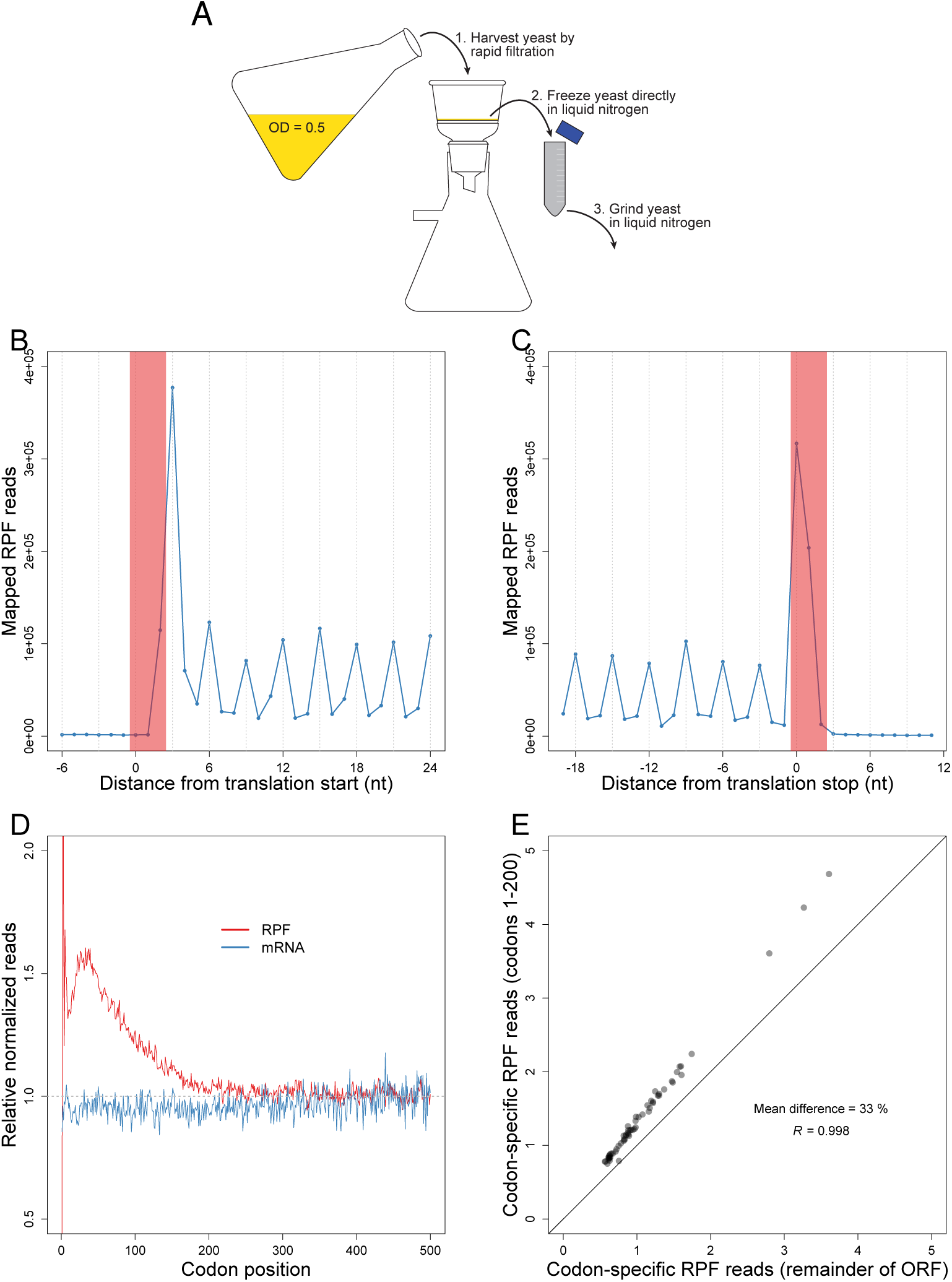
Unperturbed RPFs reveal a codon-independent 5’ ramp. (**A**) Outline of the flash-freeze protocol performed without CHX pre-treatment. (**B-C**) Metagene analyses of RPFs. Coding sequences were aligned by their start (**B**) or stop (**C**) codons (red shading). Plotted are the numbers of 28-30 nt RPF reads with the inferred ribosomal A site mapping to the indicated position along the ORF. (**D**) Metagene analyses of RPFs and RNA-seq reads (mRNA). ORFs with at least 128 total mapped reads between ribosome-footprint (red) and RNA-seq (blue) samples were individually normalized by the mean reads within the ORF, and then averaged with equal weight for each codon position across all ORFs (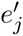 Eqn. S10 and 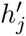 Eqn. S14). (**E**) Comparison of codon-specific RPFs as a function of the 5’ ramp. For each of the codons, densities of RPFs with ribosomal A sites mapping to that codon were calculated using either only the ramp region of each ORF (codons 1-200) or the remainder of each ORF (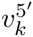 Eqn. S16 and 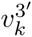 Eqn. S17, respectively). The diagonal line indicates the result expected for no difference between the two regions.

Another improvement was in the reaction used to attach the 5′ adapter sequence. The original protocol used cDNA circularization (Ingolia et al., 2009), which can introduce a strong sequence-specific bias at the 5′ ends of reads (Artieri and Fraser, 2014a). Some subsequent protocols ligate to an RNA adapter prior to cDNA synthesis (Guo et al., 2010), which has other biases. Although the biases from either circularization or ligating adaptors are not expected to influence results of analyses performed at the level of whole mRNAs, they might influence results of higher-resolution analyses, such as those at the level of codons. Borrowing from methods developed for small-RNA sequencing (Jayaprakash et al., 2011; Sorefan et al., 2012), we minimized these other biases by ligating to a library of adapter molecules that included all possible sequences at the eight 3′-terminal nucleotides.

### The 5′ ramp of ribosomes

Using the 5′ ends of ribosome footprints and the known geometry of the ribosome, we inferred the position of the A site on each footprint and thereby identified the codon that was being decoded (Ingolia et al., 2009). Analysis of all mapped reads revealed the expected three-nucleotide periodicity along the ORFs, as well as ribosome accumulation at the start (Figure 1B) and stop (Figure 1C) codons that presumably reflects slow steps following 60*S* subunit joining and preceding subunit dissociation, respectively.

To examine the global landscape of 80*S* ribosomes, we averaged the position-specific ribosome-footprint densities of individual genes into a composite metagene in which each gene is first normalized for its overall density of ribosome footprints (i.e., RPKM of RPFs) and then weighted equally in the average (Eqn S10). We observed a small 5′ “ramp” of ribosomes in this metagene, with excess ribosome footprints across the first ∼200 codons compared to the remainder of the ORF (Figure 1D). Compared to previous studies, the ramp in our dataset spanned a similar distance from the start codon but had a much smaller amplitude, with the excess relative ribosome-footprint density (*e*’_*j*_ Eqn S10) following the first ten codons peaking at ∼60% in our data compared to 110%–300% in other studies (Figure S1) (Ingolia et al., 2009; Gerashchenko et al., 2012; Zinshteyn and Gilbert, 2013; Artieri and Fraser, 2014b; Guydosh and Green, 2014; McManus et al., 2014). The trend towards decreasing ribosome-footprint density with codon position was also evident on a gene-by-gene basis in our data, as 82% of genes exhibited a decreasing number of raw RPF reads along their entire gene-length based on linear-regression of RPF reads with codon position (binomial test, *p* < 10^-15^).

For some previous ribosome-profiling studies, CHX pre-treatment presumably contributed to the size of the observed 5′ ramp. To examine this possibility, we used a whole-cell stochastic model of yeast translation (Shah et al., 2013) to simulate protein translation in a yeast cell in the presence of CHX, and we found that simulated CHX pre-treatment can indeed induce 5′ ramps of up to 300% (Figure S2). In these simulations, the ramp is due to both non-instantaneous CHX binding, which enables newly initiated ribosomes to begin translating the ORF before being initially arrested, as well as reversible binding, which enables elongating ribosomes to translocate along the ORF as they undergo cycles of CHX binding, dissociation, and rebinding. Indeed, simulated CHX pre-treatment can induce ramps of different shapes and sizes depending on the on- and off-rates of CHX binding (Figure S2). Thus, CHX pre-treatment might be responsible for the large ramps observed in other datasets, as has been recently demonstrated in experiments with variable amounts of CHX (Gerashchenko and Gladyshev, 2014). However, CHX pre-treatment cannot be responsible for the more modest 60% ramp observed in our dataset, since our protocol did not involve such treatment.

The 5’ ramp of ribosomes has previously been attributed to slower elongation due to preferential use of codons corresponding to low-abundance cognate tRNAs in the 5’ ends of genes (Tuller et al., 2010). To determine the contribution of codon usage, we re-analyzed our ribosome-profiling data to determine whether differences in ribosome-footprint densities between the 5’ and 3’ ends of a gene depend on codon choice. For each of the 61 sense codons, the average density of ribosome footprints was 33% greater when the codon fell within the first 200 codons of an ORF than when it fell within the remainder of the ORF (Figure 1E). We observed similar results, though of an increased magnitude ranging from 38–89%, when analyzing data from previous ribosome-profiling studies (Figure S1). Thus, even the same codon triplet had elevated ribosome densities in the 5’ ends of ORFs compared to 3’ ends. Consistent with these experimental results, our simulation of protein translation (now in the absence of CHX) indicated that codon ordering could account for only a 20% ramp, over a large range of simulated parameters (Figure S2). These simulation results suggested that codon ordering may explain some of the ∼60% ramp observed in our dataset, but that the majority of the ramp in our dataset was likely caused by mechanisms other than patterns of codon usage (see Discussion).

### Codon-specific elongation dwell times are inversely correlated with tRNA abundances

In our dataset, the 61 sense codons varied in their average RPF densities by more than 6 fold (Figure 1E), suggesting that different codons are decoded at different rates. Molecular biologists have long assumed that such differences in elongation rates among codons are caused by corresponding differences in the cellular abundances of cognate tRNAs (Ikemura, 1981, 1985; Andersson and Kurland, 1990; Bulmer, 1991). Several early experiments provided empirical support for this view (Varenne et al., 1984; Sorensen and Pedersen, 1991; Zhou et al., 1999). However, early ribosome-profiling studies do not report the strong anti-correlation between ribosome-footprint density and cognate tRNA abundance expected from this model (Ingolia et al., 2011; Li et al., 2012; Qian et al., 2012; Charneski and Hurst, 2013; Zinshteyn and Gilbert, 2013).

Suspecting that our improved methods might more precisely map the positions of the ribosomes during normal translation, we examined the relationship between our experimentally measured codon occupancies and measures of cognate tRNA abundance. The codon-specific excess ribosome densities (*v*_*k*_, Eqn S19) were strongly anti-correlated with cognate tRNA abundances, as estimated by copy numbers of tRNA genes and wobble parameters (Figures 2A–B). As expected, this correlation was specific to the codon within the A site, with residual correlations at the P and E sites, which were potentially caused by some 5′ heterogeneity of ribosome footprints. The codon-specific excess ribosome densities were also anti-correlated with direct estimates of tRNA abundances obtained from our RNA-seq measurements (Figure S3, Table S1). In the meantime, others using flash-freezing without CHX pre-treatment have recently reported similar findings (Gardin et al., 2014).

**Figure 2:**
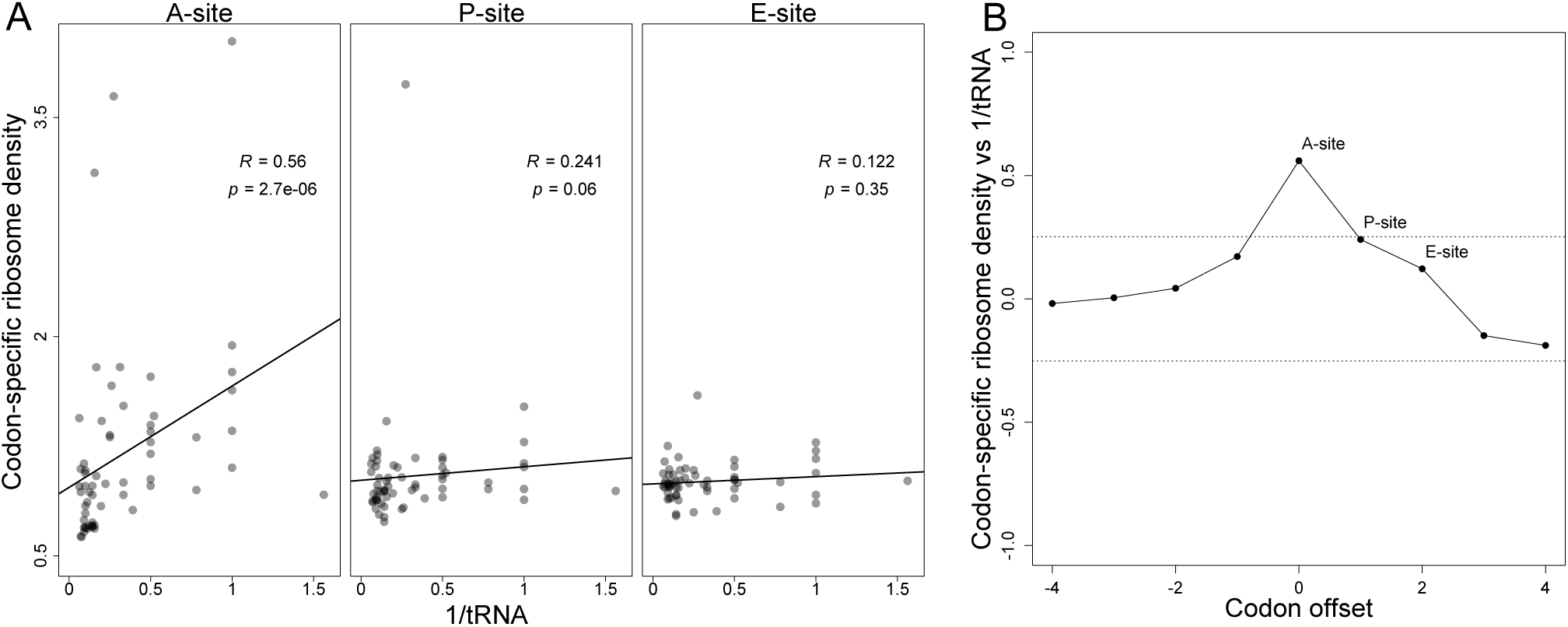
Codons corresponding to lower-abundance tRNAs are decoded more slowly. (**A**) Correlation between codon-specific excess ribosome densities and cognate tRNA abundances. Codons within RPFs were assigned to the A-, P-, and E-site positions based on the distance from the 5’ ends of fragments, and codon-specific excess ribosome densities were calculated (*v*_*k*_, Eqn. S19). Cognate tRNA abundances for each codon were estimated using the genomic copy numbers of iso-accepting tRNAs and wobble parameters (Table S4). Spearman R values are shown, with their significance (p values). (**B**) The correlations of codontRNA abundance at different positions relative to the A site. Analysis was as in (**A**) using varying offsets from the A-site position within RPFs (x axis) to calculate Spearman correlations (y axis). Dotted lines reflect the expected correlations at *p* = 0.05 significance.

When examining previously published ribosome-profiling datasets, we found that whenever CHX pre-treatment was employed, the relationship between ribosome occupancy and tRNA abundance was absent (Figure S3). We also report elsewhere a re-analysis of data from a study that used a wide range of CHX concentrations (Gerashchenko and Gladyshev, 2014) supporting our hypothesis that CHX treatment systematically disrupts the measured positions of ribosomes. Moreover, the concordance between these CHX pre-treatment datasets indicated a systematic bias (Figure S3), suggesting that an orthogonal set of mRNA sequence biases influence CHX binding. Taken together, these results strongly support the idea that differential cognate tRNA abundances drive differential elongation times among codons, as can be revealed using ribosome-profiling experiments that do not pre-treat with CHX.

At least three considerations help explain why CHX pre-treatment is expected to disrupt the correlation between tRNA abundances and measured ribosome densities at the A site. The first is that CHX, once bound to a ribosome, allows for an additional round of elongation before halting ribosomes (Schneider-Poetsch et al., 2010), which alone would remove a correlation at the A site and transfer it to the P site. Second, CHX binding is reversible, and at concentrations typically used in ribosome-profiling protocols, additional rounds of elongation might occur between CHX-binding events. Third, CHX prevents translocation of the ribosome by binding to the E site, with space for a deacylated tRNA (Schneider-Poetsch et al., 2010), and thus CHX binding affinity presumably varies with features of the E site and perhaps binding of the E-site tRNA if it influences CHX binding. Thus, in the presence of CHX pre-treatment, the ribosome density at a site is likely more a function of the on and off rates of CHX binding than a function of differential isoaccepting tRNA availability.

### Polybasic stretches induce widespread pausing of ribosomes

Aside from the abundance of cognate tRNA, the ribosome-footprint density at a particular codon in an ORF might also be influenced by interactions between the emerging nascent polypeptide and the ribosome. Due to the negatively charged nature of the ribosome exit tunnel, polybasic tracts tend to make extensive electrostatic interactions with the tunnel that are thought to stall elongation (Lu and Deutsch, 2008; Charneski and Hurst, 2013). Consistent with this hypothesis, we observed a peak of excess ribosome-footprint density (*z*_*ij*_, Eqn S7) roughly 9 amino acids after the start of highly positively charged regions (defined as six arginine or lysine residues within a 10 amino-acid window), which would position the basic residues within the exit tunnel (Figure 3A). For windows containing fewer basic residues, the pause amplitude steadily decreased as the number of basic residues decreased, but pausing was still apparent with as few as three basic residues within a 10 amino-acid window.

**Figure 3:**
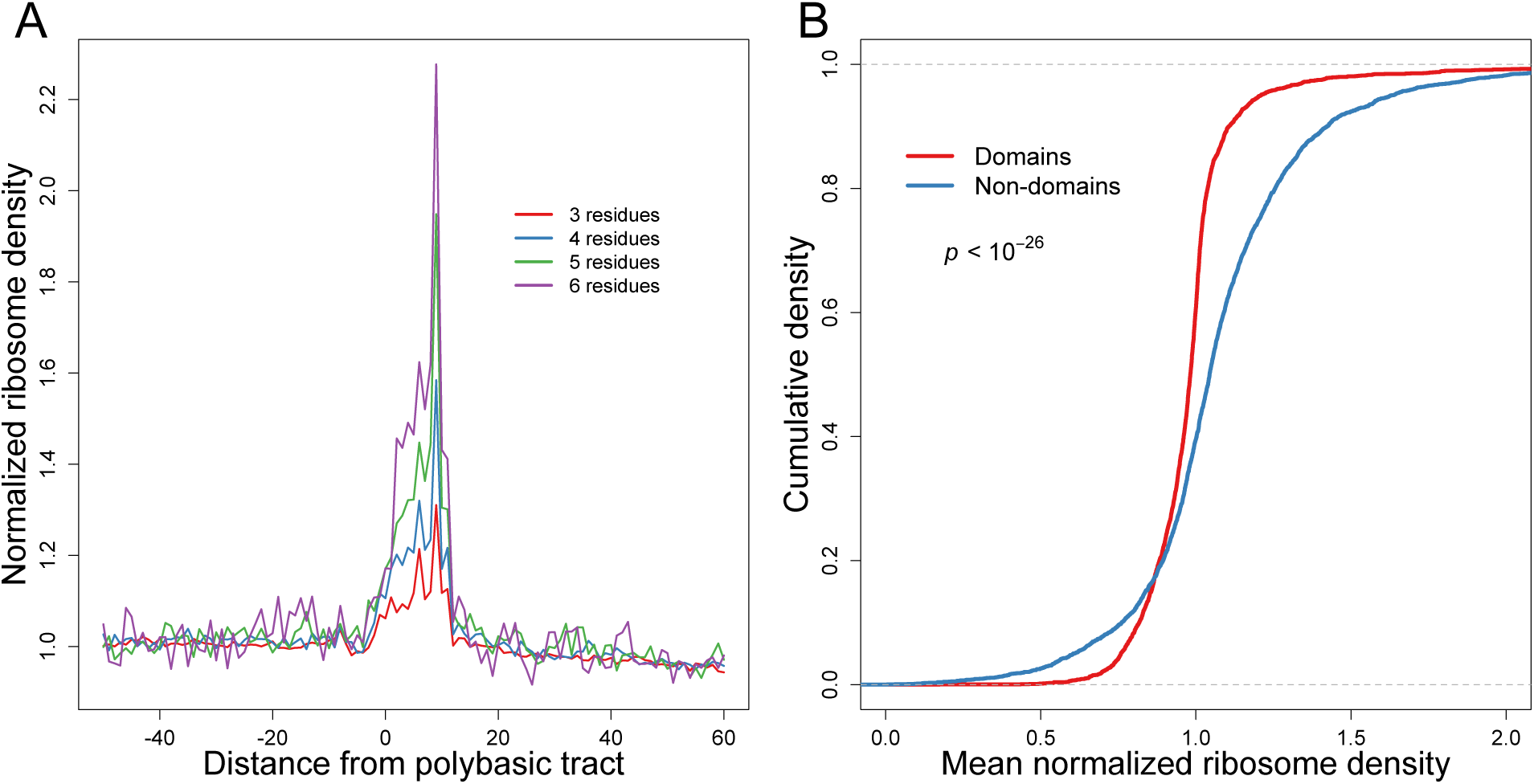
Elongation dynamics correlate with amino acid sequence and domain architecture. (**A**) Metagene analysis of normalized ribosome density surrounding polybasic tracts. Regions within ORFs that contained the indicated number of basic residues (arginine and lysine) within a stretch of 10 amino acids were aligned by the start of the region. Plotted are the normalized ribosome densities (*z*_*ij*_, Eqn. S7) observed at each codon position. (**B**) Cumulative distributions of normalized ribosome densities within and outside of protein-folding domains. Mean normalized RPF densities (*z*_*ij*_, Eqn. S7) for codons within the domain-encoding and non-domain-encoding regions were individually calculated for each ORF. Domain assignments were based on InterProScan classifications (JONES et al. 2014) obtained from the Superfamily database (WILSON et al. 2009). Statistical significance was evaluated using paired t-test (*p* < 10^-26^).

Published ribosome-profiling datasets that used CHX pre-treatment all fail to show this pattern (Figure S4), which again suggested that drug treatment obscures the locations of natural elongation pauses. Indeed, results related to ours were observed in an earlier ribosome-profiling study conducted without CHX pre-treatment, although in that study analyses were limited to only 103 highly charged regions (Brandman et al., 2012). In contrast, our analyses quantified ribosome-footprint densities surrounding polybasic stretches ranging from ∼1400 windows containing six basic residues to ∼48,000 windows containing three basic residues, which not only confirmed that elongation tends to stall as polybasic stretches reach the exit tunnel but also showed that this effect is far more widespread than anticipated. When simulating ribosome density surrounding polybasic stretches, we found a relative depletion of ribosomes in polybasic stretches (Figure S5), which indicated that the elongation stalls within polybasic stretches were not caused by biased codon usage and suggested that the observed excess ribosome-footprint density likely underestimated the direct effect of polybasic stretches on the elongation rate.

### Slower elongation at regions encoding inter-domain linkers

The modulation of ribosome-footprint densities by either tRNA abundances (Figure 2A) or polybasic stretches (Figure 3A) would be expected to influence the kinetics of co-translational folding. Indeed, slower elongation rates within interdomain linkers relative to the adjacent domains is reported to coordinate cotranslational folding of nascent polypeptides (Thanaraj and Argos, 1996; Kimchi-Sarfaty et al., 2007; Pechmann and Frydman, 2013). However, systematic experimental evidence for such differences in elongation rates has been lacking.

To examine whether our ribosome-profiling data reveals such differences, we first used InterProScan classifications (Jones et al., 2014) based on the Superfamily database (Wilson et al., 2009) to partition coding sequences into domain and linker regions. For each ORF we calculated the mean normalized ribosome-footprint densities (*z*_*ij*_, Eqn S7) for codons within the domain- and linker-encoding regions. Comparing between these subsets of sites, we found significantly lower densities in regions of genes that fell within domains compared to regions that fell outside of domains (Figure 3B, mean difference 0.094, paired t-test, *p* < 10^-26^). To eliminate any influence of the 5′ ramp, we repeated the analyses, excluding the first 200 codons. Although the size of the effect was smaller when excluding the first 200 codons (mean diff = 0.029), the difference in mean ribosome densities was still significant (*p* = 0.0002), indicating that the 5′ ramp was not solely responsible for differences in ribosome densities between domains and other regions of proteins (Figure S6A).

The trend towards relatively lower ribosome densities in domain regions holds even when restricted to each individual amino acid, with the exceptions of cysteine residues and the single-codon-encoded methionine and tryptophan residues (Figure S7). Thus, differences in amino-acid content between domains and linkers cannot account for the observed differences in bound ribosome densities. Moreover, for 54 out of 61 sense codons, we find significantly higher ribosome densities in domains compared to linkers (one-sides t-test, *p* < 0.05). We find significantly higher ribosome densities in domains even after excluding the first 200 codons, for 26 out of 61 codons (one-sides t-test, *p* < 0.05). This result implies that differences in synonymous codon usage between domain and linker region cannot alone account for the differences in ribosome densities. One possible mechanism for differential ribosome occupancy, independent of codon usage, is differential recruitment of chaperones and their associated effects on co-translational folding (Ingolia, 2014).

Similar results comparing ribosome densities in domain and linker regions were obtained when using InterProScan classifications based on the Pfam domain database (Bateman et al., 2002) instead of the Superfamily database (Figure S6B). Finally, consistent with earlier computational analyses (Pechmann and Frydman, 2013), analysis of our data indicated that differences in elongation rate exist at the level of protein secondary structures as well: regions corresponding to helices and sheets exhibited significantly lower ribosome-footprint densities than regions corresponding to loops (Figure S6C). Although requirements for cotranslational folding have long been hypothesized to influence elongation rates, these results provided the first systematic empirical support of this claim. Nonetheless, the magnitude of the signal was very small, suggesting that slower inter-domain elongation either has very little impact or impacts very few genes.

### Estimates of protein-synthesis rates

Taken together, our results indicate that the ribosome-footprint density at a given codon position within a gene is influenced by several factors, including the abundance of cognate tRNAs and whether the codon is immediately downstream of a polybasic stretch, falls within a protein domain, or lies in the 5’ region of the ORF. The non-uniform ribosome density along individual ORFs implies that the overall ribosome-footprint density on each gene (i.e., RPKM of RPFs) does not directly reflect the rate of protein synthesis (Li et al., 2014a). For example, the ribosome-footprint densities of genes enriched in more slowly elongated codons would tend to overestimate their protein synthesis rates; and the same would be true for shorter ORFs.

To more accurately quantify the protein synthesis rates of individual genes from ribosome-footprint densities, we used empirically derived correction factors to account for the position- and codon-specific effects we have observed (*f*_*j*_, Eqn S23). The ∼74.3 million RPFs that we sequenced enabled reliable estimates of protein-synthesis rates for 4839 genes (Eqn S28).

### Accurate measurement of yeast mRNA abundances

In addition to improving measurements of ribosome-footprint densities, we sought to improve measurements of mRNA abundances, which is also critical for accurately quantifying translational control. Prior experiments have typically measured yeast mRNA abundances by performing RNA-seq on poly(A)-selected RNA (Ingolia et al., 2009; Gerashchenko et al., 2012; Zinshteyn and Gilbert, 2013; Artieri and Fraser, 2014b; Guydosh and Green, 2014; McManus et al., 2014). However, poly(A) selection might bias mRNA-abundance measurements. For example, mRNAs that lack a poly(A) tail of sufficient length to stably hybridize to oligo(dT) might not be as efficiently recovered. Although *S. cerevisiae* is not known to contain translated mRNAs that altogether lack a poly(A) tail, the lengths of poly(A) tails found on *S. cerevisiae* mRNAs are relatively short (median length of 27 nt) (Subtelny et al., 2014). Another source of potential bias in poly(A)-selection is partial recovery of mRNAs endonucleolytically cleaved during RNA isolation or poly(A)-selection. The 5′ fragments resulting from mRNA cleavage are not recovered by poly(A) selection, which causes a 3′ bias in the resulting RNA-seq data (Nagalakshmi et al., 2008). Indeed, analyses of published RNA-seq datasets from ribosome-profiling studies revealed a severe 3′ bias in poly(A)-selected RNA-seq reads, ranging from 19– 130% excess reads (Eqn S15) (Figure S8). Because longer mRNAs have a higher probability of being cleaved, the abundances of longer mRNAs might be systematically underestimated by poly(A) selection (Figure S9).

One alternative to poly(A) selection is ribosomal RNA (rRNA) depletion, which enriches mRNAs by removing rRNA using subtractive hybridization. A potential concern with subtractive hybridization is the depletion of mRNAs that either cross-hybridize to the oligonucleotides used to remove rRNA sequences or adhere to the solid matrix to which the oligonucleotides are attached. To investigate the extent to which unintended mRNA depletion is a problem for the commercial reagents used for yeast RNA-seq library preparations, we subjected the same total RNA to each of three procedures: Dynabeads oligo(dT)_25_ (Life Technologies), RiboMinus Yeast Transcriptome Isolation Kit (Life Technologies), or Ribo-Zero Yeast Magnetic Gold Kit (Epicentre). As a reference, we also generated an RNA-seq library from the total RNA that was not enriched or depleted and therefore contained primarily rRNA (90.2% of ∼199.7 million genome-mapping reads). Although this reference sample was critical here for evaluating the biases of mRNA enrichment methods, performing RNA-seq on total RNA is not ideal as a general approach because of the large number of reads required to obtain sufficient coverage of the mRNA transcriptome. We also note that we used total RNA extracted from the lysate that was used for ribosome footprint profiling, as opposed to RNA extracted from whole cells as done in the original ribosome-profiling study (Ingolia et al., 2009). When comparing the 4540 mRNAs for which we obtained at least 64 reads in our total RNA library, only the Ribo-Zero-treated sample faithfully recapitulated the mRNA abundances observed in total RNA (*R*^2^=0.98, Figure 4A, Figure S10). The poly(A)-selected and RiboMinus-treated samples each had significantly lower correlations with total RNA (*R*^2^=0.85 and *R*^2^=0.87, respectively), indicating a skewed representation of the transcriptome. Compared to published RNA-seq data from ribosome-profiling studies, our Ribo-Zero-treated sample also exhibited the highest correlations with microarray-based estimates of mRNA abundances (Figure S11).

**Figure 4:**
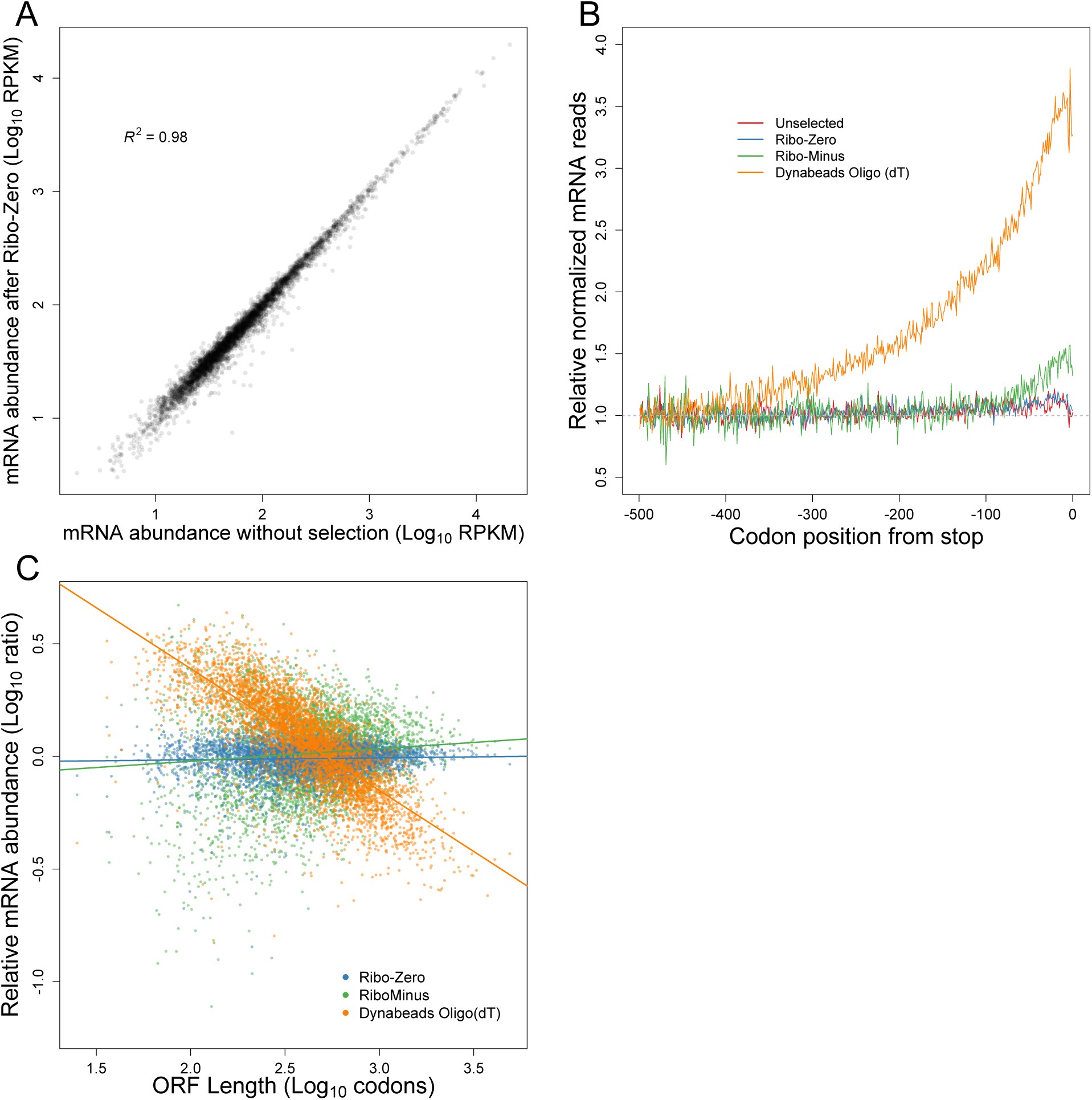
mRNA enrichment methods can bias mRNA abundance measurements. (**A**) mRNA abundances measured by RNA-seq of Ribo-Zero-treated RNA compared to those measured by RNA-seq of total unselected RNA. Pearson *R*^2^ is indicated. (**B**) Metagene analysis of RNA-seq read density in total unselected or mRNA-enriched RNA samples. Coding sequences were aligned by their stop codons, and RNA-seq reads were individually normalized by the mean reads within the ORF and then averaged with equal weight for each codon position across all ORFs (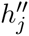 Eqn. S15). **C**)mRNA abundances for mRNA-enriched samples relative to total unselected RNA, as a function of ORF length.

The poly(A)-selected and RiboMinus-treated samples were more poorly correlated with each other (*R*^2^=0.755, Figure S10) than with the total RNA sample, indicating that the poly(A)-selected and RiboMinus-treated samples were affected by different biases. As anticipated, the poly(A)-selected sample contained a strong 3′ bias (Figure 4B), which caused a systematic underestimation of the abundances of longer genes (Figure 4C). After accounting for this strong bias in the poly(A)-selected sample, we did not detect a relationship between poly(A)-tail length and poly(A)-selection efficiency, suggesting that tail-length differences did not significantly contribute to the biases of poly(A)-selected RNA-seq data. In the case of the RiboMinus-treated sample, we suspect that the skewed mRNA abundances were likely due to cross-hybridization of the depletion probes to mRNAs. Such effects were largely absent from the Ribo-Zero-treated sample, perhaps owing to the more stringent hybridization conditions of the Ribo-Zero protocol. The RiboMinus-treated sample also suffered from severe rRNA contamination (44.5% of reads, originating primarily from the 5*S* rRNA), which correspondingly reduced coverage of the mRNA transcriptome.

Interestingly, even the total-RNA dataset contained a small 3′ bias, which was also observed in the Ribo-Zero-treated RNA (Figure 4B). This bias was consistent with the notion that the bodies of yeast mRNAs are primarily degraded in the 5′-to-3′ direction by the Xrn1 exoribonuclease (Hu et al., 2009). The decay intermediates of this vectorial degradation process would contribute more reads toward the 3′ ends of mRNAs, giving rise to the observed bias (especially when considering that our RNA samples were enriched for cytoplasmic RNA, which would diminish the countervailing vectorial mRNA synthesis process occurring in the nucleus). Together our results indicate that Ribo-Zero treatment enables deep coverage of the yeast transcriptome without substantially biasing mRNA abundances. For all subsequent analyses we use mRNA abundances estimated from Ribo-Zero-treated RNA.

### A narrow range of initiation efficiencies in log-phase yeast

We used our protein-synthesis rates and mRNA abundances to estimate the translation-initiation efficiencies of each gene. Because protein synthesis is typically limited by the rate of translation initiation (Bulmer, 1991; Shah et al., 2013), we defined the initiation efficiency (IE) of a gene as its protein-synthesis rate divided by its mRNA abundance (Eqn S27). Thus, the IE measure quantified the efficiency of protein production per mRNA molecule of a gene, in a typical cell. A wide range of IEs among genes would indicate that protein production is under strong translational control, whereas a narrow range of IEs would indicate that protein production is typically governed by mRNA abundances, and hence protein-synthesis rate is primarily controlled by mRNA transcription and decay. To facilitate comparisons with published datasets, we also calculated the translational efficiencies TEs, which have previously been used to quantify translational control (Ingolia et al., 2009). The TE value of a gene is the ribosome-footprint density normalized by the mRNA abundance. Because TE is calculated based on the ribosome-footprint density rather than the protein synthesis rate, TE does not account for differential rates of elongation associated with the 5′ ramp or codon identity. Nonetheless, IE and TE were highly correlated (*R* = 0.951, Figure S12).

The first ribosome-profiling study suggested a large amount of translational control in yeast, with the range of TEs reported to span roughly 100 fold (Ingolia et al., 2009). Indeed, we found that the 1–99 percentile range of TEs in those data spanned 73 fold (Figure S13). In contrast, the range of TEs observed in our data was narrower, with the 1–99 percentile spanning only a 15-fold range (Figure 5A). Although the range of IEs was marginally wider than that of TEs (1– 99 percentile spanning 21 fold, Figure S12), it was still substantially smaller than the range of TEs reported previously (Ingolia et al., 2009). The relatively narrow range of IEs in our data was also reflected by the high correlation between mRNA abundance and protein synthesis rate (*R*=0.948; Figure 5B), indicating that protein-synthesis rates are largely dictated by mRNA abundances with minimal contributions of differential initiation efficiencies (Csárdi et al. 2015). Consistent with this idea, when we examined mass-spectrometry-based measurements of steady-state protein abundance (de Godoy et al., 2008), we found indistinguishable correlations with mRNA abundances as with protein synthesis rates (Figure 5C). These analyses all suggest that mRNA abundance is a strong predictor of total protein production. Importantly, the range of ribosome-footprint densities closely mirrored the range of mRNA abundances except for a tail of lowly translated genes (Figure 5A, inset), indicating that translational control modestly expands the dynamic range of protein expression in log-phase yeast (Csárdi et al. 2015).

**Figure 5:**
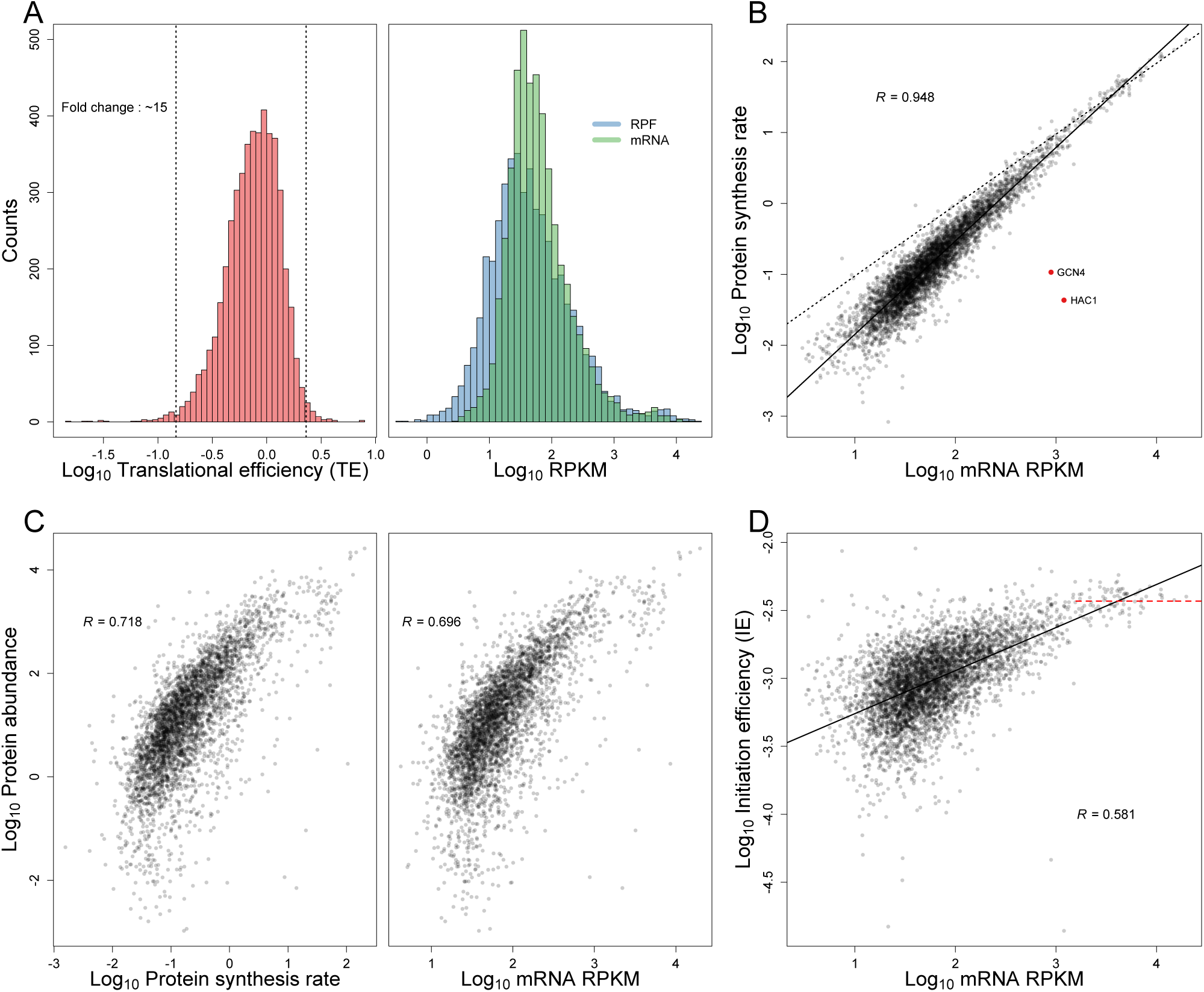
TEs and IEs span a narrow range in log-phase yeast cells. (**A**) Distributions of TE, RPF and mRNA measurements. The TE distribution is on the left, with vertical dashed lines marking the 1^*st*^ and 99^*th*^ percentiles, and the fold-change separating these percentiles indicated. Overlaid distributions of RPF densities (blue) and mRNA abundances (green) are on the right. All ORFs with at least 128 total reads between the ribosome-profiling and RNA-seq datasets were included (except YCR024C-B, which was excluded because it is likely the 3’ UTR of PMP1 rather than an independently transcribed gene). (**B**) Relationship between and protein-synthesis rate and mRNA abundance for genes shown in (**A**). GCN4 and HAC1 (red points) were the only abundant mRNAs with exceptionally low protein-synthesis rates. The best linear least-squares fit to the data is shown (solid line), with the Pearson R. For reference, a one-to-one relationship between protein synthesis rate and mRNA abundance is also shown, which fits well to the data for genes with RPKM > 1000 (dashed line). (**C**) Relationship between with experimentally measured protein abundance (de Godoy et al., 2008) and either protein-synthesis rate (left) or mRNA abundance (right). The 3,845 genes from (**A**) for which protein-abundance measurements were available were included in these analyses. Pearson correlations are shown (R). (**D**) Relationship between mRNA abundance and IE for genes shown in (**A**). The best linear least-squares fit to the data is shown (solid line), with the Pearson R. The plateauing of IE, observed for the genes with RPKM > 1000, is also indicated (dashed red line).

When we examined the range of TEs in other published datasets, we also found a more narrow range (as low as 22 fold from 1–99 percentiles) than that of Ingolia et al. (2009) (Figure S13). However, the TEs in published datasets— which were all generated using poly(A)-selected mRNA—are not particularly well correlated with each other (Table S2). These discrepancies in TEs are largely due to differences in measured mRNA abundances, whereas the ribosome-footprint abundances correlated almost perfectly. Collectively, these results indicate that the amount of translational control in log-phase yeast has been overestimated due to inaccuracies in TE measurements, largely caused by challenges in accurately measuring mRNA levels.

We also noticed that the shape of the TE distribution from our data, which was asymmetric, differed from that of the Ingolia data, which is highly symmetric. In particular, in our data there were relatively few genes in the right tail of the distribution (Figure 5A, note the location of the mode closer to the 99^th^ than the 1^st^ percentile). This observation implied that mRNAs from very few genes contain elements that impart an exceptionally high initiation efficiency and are thereby “translationally privileged”. Rather, most mRNAs initiate close to a maximum possible rate (likely set by the availability of free ribosomes or initiation factors) or contain features that modestly reduce the initiation rate.

To the extent that differences in IE were observed, the genes with lower IE, tended to be expressed at lower mRNA levels, with IE increasing roughly linearly with mRNA expression levels (Figure 5D). These results were consistent with the notion that abundant mRNAs have undergone evolutionary selection to be efficiently translated (Bennetzen and Hall, 1982; Gouy and Gautier, 1982; Sharp and Li, 1987; Andersson and Kurland, 1990; Plotkin and Kudla, 2011; Shah and Gilchrist, 2011). Interestingly, this effect plateaued for the highest expressed mRNAs (RPKMs >1000), for which the differences in protein syntheses essentially matched the differences in mRNA (Figure 5B and Figure 5D, dashed lines), which suggested that the efficiency for the highest expressed mRNAs has reached a level that is difficult to surpass.

We observed two notable outliers in the comparison of mRNA abundances and synthesis rates (Figure 5B, red dots). These two, which correspond to relatively abundant mRNAs with exceptionally low synthesis rates, were *HAC1* and *GCN4*. These are the two most well-known examples of translational control in log-phase yeast, and they are both involved in rapid stress responses. *HAC1* encodes a transcription factor that mediates the unfolded protein response (Cox and Walter, 1996; Kawahara et al., 1997). In the absence of protein-folding stress, *HAC1* is translationally repressed due to base pairing interactions between the 5′ UTR and intron (Ruegsegger et al., 2001). Upon stress, the non-canonical intron is spliced out, which relieves the translational repression and enables rapid expression of Hac1 protein. *GCN4* encodes a transcriptional activator that is the primary regulator of the transcriptional response to amino acid starvation (Hope and Struhl, 1985). The 5′ UTR of *GCN4* mRNA contains multiple uORFs that prevent translation of the Gcn4-coding ORF under nutrient replete conditions (Mueller and Hinnebusch, 1986). During amino acid starvation, reduced levels of the eIF2 ternary complex enable scanning ribosomes to bypass some of the uORFs and initiate translation at the *GCN4* start codon (Dever et al., 1992). The observation that *HAC1* and *GCN4* were the only abundant mRNAs that were strongly regulated at the translational level further emphasized that translational control only subtly modulates the protein production of most yeast genes.

### Potential contribution of translational control to proportional synthesis

Although more narrow than reported in earlier ribosome-profiling studies, the IE distribution that we observed was still large enough to enable the cell to tune synthesis rates via translational control. One scenario in which this might be important is in the proportional synthesis of the subunits for multiprotein complexes. A recent genome-wide study of protein synthesis rates in *E. coli* and *S. cerevisiae* concluded that components of multisubunit complexes are usually synthesized in precise proportion to their stoichiometry (Li et al., 2014a). In *E. coli*, the subunits of multiprotein complexes are usually encoded on the same polycistronic mRNA and thus can be synthesized in different proportions only if they have different translation-initiation rates. In eukaryotes, however, the subunits of protein complexes are encoded on separate mRNAs, which enables proportional synthesis to be achieved through control of mRNA abundance (via transcription rate and mRNA half-life). Nonetheless, translational control might still compensate for differences in mRNA abundance and thereby achieve more precise stoichiometry of synthesis rates.

To explore this possibility, we examined the synthesis rates, mRNA abundances, and IEs of the subunits of stably associated complexes previously shown to undergo proportional synthesis (Li et al., 2014a). mRNAs encoding subunits of heterodimeric complexes had roughly similar abundances (within 2 fold), indicating that most of their proportional synthesis is achieved through coordinated mRNA levels (Figure 6A). The same was true for mRNAs encoding multiprotein complexes, after accounting for subunit stoichiometry (within ∼2 fold, Figure 6B), as well for mRNAs encoding heterodimeric complexes containing alternative paralogous subunits (within 1.4 fold, Figure 6C). These observations were consistent with the narrow range of IEs in yeast; with limited translational control, proportional synthesis requires roughly proportional mRNA levels. However, in 12 out of 18 cases subunit stoichiometry was more accurately reflected by synthesis rates than by mRNA abundances (Figures 6Ans), as quantified by the coefficients of variation (Figure 6D). For example, the subunits of the heterodimeric FACT complex were translated at equal levels despite the Spt16-encoding mRNA being 59% more abundant than the Pob3-encoding mRNA. Similarly, in the mitochondrial alpha-ketoglutarate dehydrogenase complex, higher expression of the Kdg2 subunit relative to the Kdg1 subunit (which are present in a 2:1 stoichiometry in the complex) was achieved entirely at the level of translation. Although the fraction of 12 of 18 did not pass our threshold for statistical significance (*p* = 0.07, binomial test), these results hinted that translational control compensates for small differences in mRNA levels to achieve more proportional synthesis. We also found that mRNAs encoding proportionally synthesized subunits of heterodimeric complexes tended to have similar IEs (*R*^2^ = 0.72, Figure 6A), suggesting that such mRNAs might be coregulated at the level of translation.

**Figure 6:**
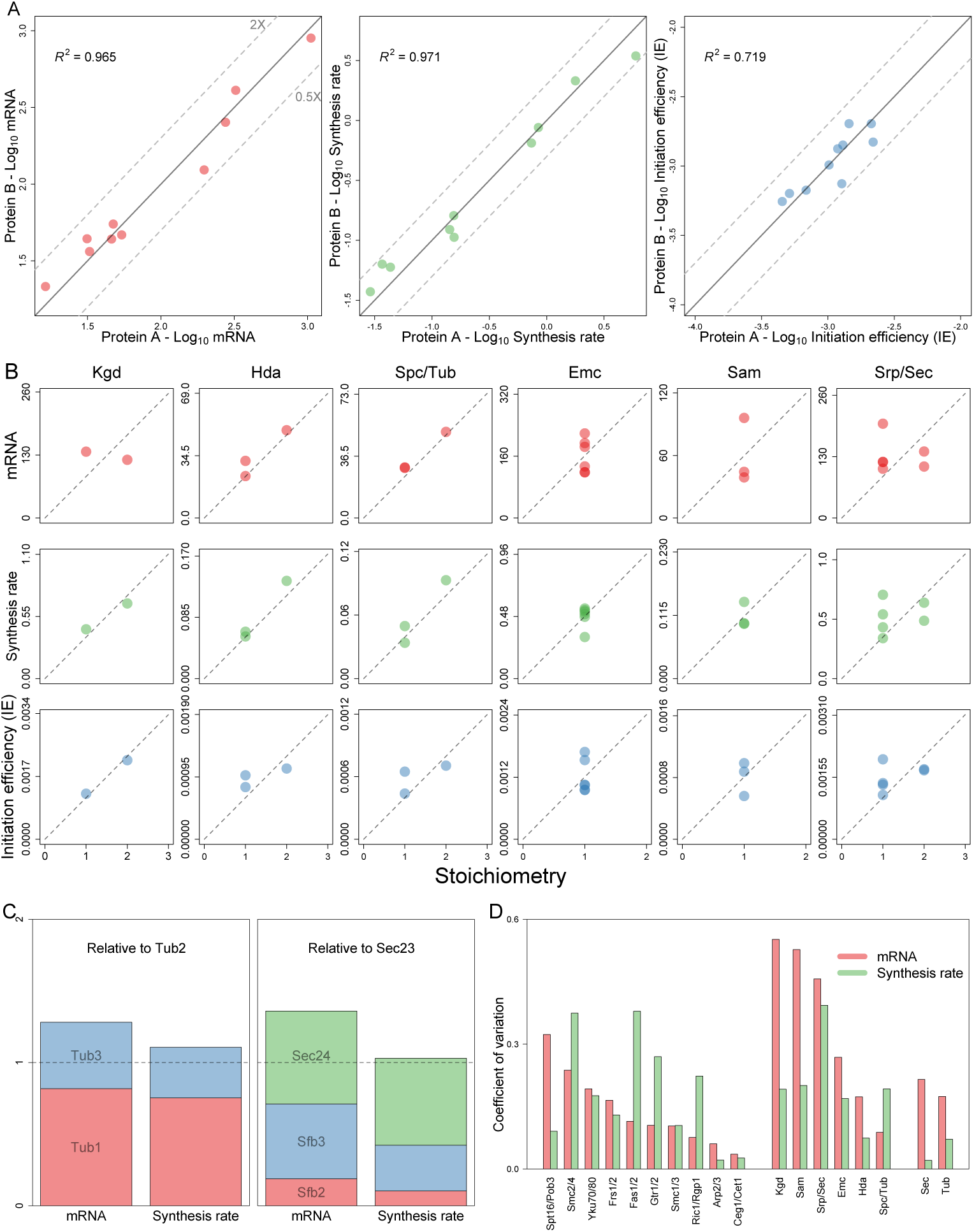
mRNA abundance and translational control contribute to proportional synthesis. (**A**) Analysis of complexes with two equimolar subunits. For each heterodimeric complex, the mRNA abundances (left), synthesis rates (middle), and IEs (right) of the individual subunits are plotted (with the subunit whose systematic name is first alphabetically on the x axis). All of the heterodimeric complexes characterized in Li et al. (2014) are shown, with the exception of the Smc2/4 complex, which substantially deviates from proportional synthesis in both our dataset and the published dataset. Dashed lines indicate 2-fold differences. (**B**) Analysis of multi-protein complexes. For each complex, the mRNA abundances, synthesis rates, and IEs of its subunits are plotted as a function of subunit stoichiometry. Dashed line passes through the origin and mean of the data-points. (**C**) Analyses of complexes containing paralogous subunits. For each complex, data for alternative subunits are plotted in the same column relative to the data for the constitutive subunit (Tub2 for the *αβ*-tubulin complex, Sec23 for the COPII vesicle coat). (**D**) Comparison of the differences in mRNA abundances with those of synthesis rates for each of the complexes in (**A**) and (**B**). Data were normalized to subunit stoichiometry, and coefficients of variation (CVs) were calculated using all subunits within each complex. CVs were calculated similarly for the Sec23- and Tub2-containing complexes, except that data for paralogous subunits (Tub1 and Tub3; Sec24, Sfb2, and Sfb3) were first summed.

### Determinants of initiation efficiencies in yeast

Next, we sought to identify sequence-based features that might explain the variation in measured IE values among genes. First we considered uORFS, which can inhibit translation by serving as decoys to prevent initiation at the start codons of bona fide ORFs, as observed in the extreme case of *GCN4*. Using recently described single-nucleotide-resolution 5′ UTR annotations (Arribere and Gilbert, 2013), we identified upstream ATGs (uATGs) in 303 out of the 2549 ORFs that had reproducibly uniform 5′ ends. Those genes containing uATGs had significantly lower IEs than genes without uATGs (Figure 7A, t-test *p* < 10^-16^), even after controlling for 5′ UTR lengths. These results confirmed that a general feature of uORFs is to decrease the translation of downstream ORFs, and that the presence of uATGs can explain some of the variance in IEs.

**Figure 7:**
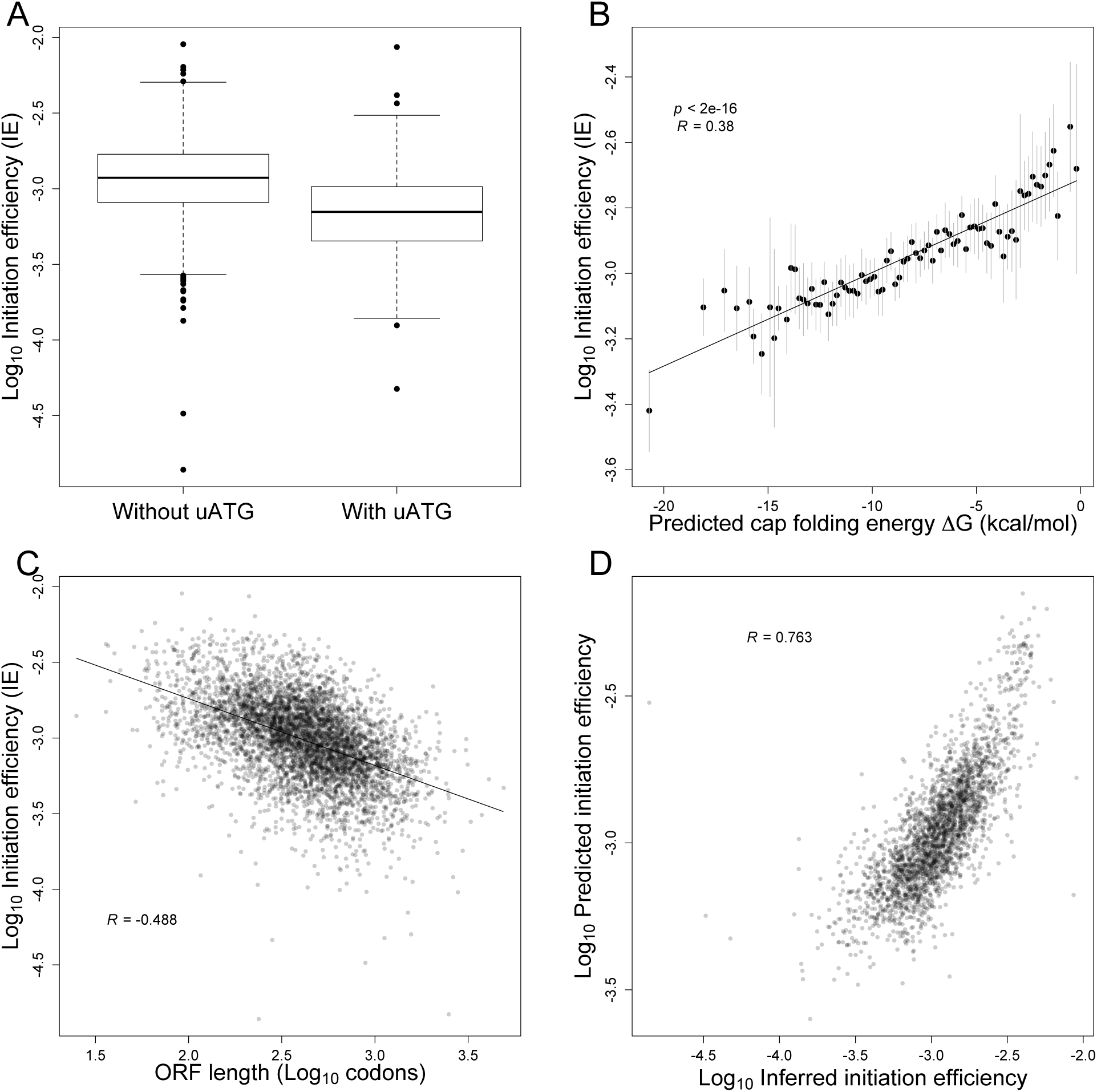
Sequence-based features of mRNAs largely explain yeast IEs. (**A**) Reduced IE values for genes with at least one upstream ATG (i.e., an ATG codon located within the annotated 5’ UTR). The plots indicated the median (line), quartile (box) and 1^*st*^ and 99^*th*^ percentiles (whiskers) of the distributions. (**B**) Inverse relationship between IE and the folding energy of predicted RNA secondary structure near the cap (Cap-folding energy). RNAfold was used to estimate folding energies for the first 70 nts of the mRNA. Gray bars indicate *±*2 SE of IE values for genes binned by predicted folding energy. The best linear least-squares fit to the data is shown (solid line), with the Pearson R. (**C**) Inverse relationship between IE and ORF length. The best linear least-squares fit to the data is shown (solid line), with the Pearson R. (**D**) Correspondence between predicted IEs and experimentally inferred IEs. Initiation efficiencies were predicted using a multiple-regression model, based on mRNA abundance and sequence-based features of the 2,549 genes with empirically determined 5’-UTRs. Shown is the Pearson *R*.

Another mRNA feature that has been linked to differences in synthesis rates is mRNA secondary structure. In bacteria, accessibility of the Shine–Delgarno sequence, which directly binds the 40*S* ribosome subunit during translation initiation, is likely the primary determinant of synthesis rates (Gold, 1988). In eukaryotes, cap-dependent translation initiation involves binding of the eIF4F complex to the cap, followed by scanning of the 40*S* ribosome to the start codon. Structure located near the 5′ cap might interfere with binding of the eIF4F cap-binding complex, while structure within the 5′ UTR could disrupt the scanning 40*S* ribosome. An open structure around the start codon might also be important for facilitating joining of the 60*S* subunit. Previous genome-wide structure analyses revealed a weak but significant inverse correlation between startcodon-proximal structure and TE (Kertesz et al., 2010; Ouyang et al., 2013), but the accessibility of the 5′ UTR more generally was not reported, and the TE values used in those studies were affected by RNA-seq biases. Incorporating improved 5′ UTR annotations (Arribere and Gilbert, 2013) and our IE measurements, we analyzed the effects of predicted secondary structure throughout the 5′ UTRs. For each mRNA with a single reproducible 5′ end, we predicted the accessibility of the 5′ cap by calculating the predicted folding energy of the sequence spanning increasing distances from the cap. For all distances examined, we observed a significant correlation between predicted cap accessibility and IE (t-test, *p* < 10^-6^ for each window; Figure 7B, Figure S14). This correlation rapidly increased with window length, approaching a maximum at 70–90 nucleotides (Pearson correlation, *R* ∼ 0.37 for windows 70-90 nts long) and then steadily declined for larger windows (Figure S14), consistent with local folding of the 5′ end determining cap accessibility. Notably, the correlations that we observed between mRNA structure and translation were the largest that have been reported between these features in eukaryotes, emphasizing the utility of our accurate IE measurements. Together, these results confirmed that mRNAs with less-structured 5′ UTRs tend to be initiated more efficiently (Godefroy-Colburn et al., 1985; Shah et al., 2013), which is consistent with eIF4F binding, 40*S* recruitment, or scanning as influential regulatory steps during eukaryotic initiation.

In addition to uORFs and mRNA structure, gene length has also been reported to correlate with translational efficiency. In early microarray-based studies of ribosome density, ORF length was strongly anti-correlated with ribosome density (Arava et al., 2003). However, our analysis of published ribosome-profiling data revealed essentially no correlation (or even a positive correlation in one case) between length and TE (Figure S15). In contrast, we observed a striking negative correlation in our IE (and correspondingly in our TE) data (Figure 7C, Figure S15). Our IE measure already corrected for the small “ramp” of elevated 5’ ribosome densities and thus the correlation between IE and ORF length was not caused by this ramp. Moreover, the negative correlation between ORF length and TE persisted even after removing the first 250 codons of each ORF, which further confirmed that the correlation was not caused by the 5’ ramp of elevated ribosome densities (Figure S15). The discrepancy between our data and earlier ribosome-profiling datasets was likely due to the RNA-seq 3′-bias caused by poly(A) selection (Figure 4B, Figure S8). Indeed, we could recover the anti-correlation between ORF length and TE in most other datasets when we controlled for the 3′ bias by estimating mRNA abundances based on mapped RNA-seq reads from only the 3′ ends of genes (Figure S16). Together, these results showed that the original report of shorter mRNAs having relatively higher initiation efficiencies (Arava et al., 2003) is correct, even after accounting for the CHX-enhanced 5′ ramp that confounded that analysis.

Based on these results, we used multiple linear regression to build a model that considered number of uATGs, predicted cap-proximal RNA-folding energy (and also GC content of the 5′ UTR as another metric for structure), and lengths of ORFs and 5′ UTRs to explain the variance in IE observed among genes. We also included an mRNA-abundance term in the model, because IE is greater for more abundant mRNAs (Figure 5D). To identify the most informative features, we used Akaike′s Information Criteria (AIC) for model selection and both step-up and step-down model-selection procedures (using the **stepAIC** function in the **MASS** package in R). The multiple regression model that best explained the variation in IE included all six variables, even after penalizing for model complexity (Figure 7D, Table S3). The dominant explanatory variable was mRNA abundance, which alone accounted for ∼40% of the variance in IE. Collectively, a model containing all six variables explained ∼58% of the variance in IE. A model that excluded mRNA abundance, and therefore depended on only sequence-based features, still explained ∼39% of the variance in IE. These results of our statistical modeling should help motivate mechanistic studies of how each of these mRNA features impacts translation.

## Discussion

### Widespread changes in elongation rate along mRNAs

We have shown that accurate measurements of both mRNA abundances and ribosome footprints can reveal new insights into the regulation and dynamics of eukaryotic translation. The native ribosome footprints that we isolated and sequenced are indicative of a dynamic and heterogeneous elongation process, with ribosomes transiting along mRNA molecules at variable rates depending on the distance from the start codon, codon identity, and polypeptide sequence.

The 5′ ramp of ribosomes that we observed was much smaller than ramps observed under CHX pre-treatment (Figure S1), which was consistent with the predicted effects of CHX. Codon usage accounted for about a third of the residual ramp we observed, but even the same codons were differentially occupied by ribosomes depending upon whether they occurred in the 5′ or 3′ ends of genes (Figure 1E), indicating that additional mechanisms must be involved. Potential mechanisms include ribosome drop-off during elongation or an overall (i.e., codon- and gene-independent) slower elongation rate during the early phase of translation. Although we cannot rule out ribosome drop-off as a contributing factor, the observation that ribosome-footprint density eventually becomes constant after 200 codons argues against a constant abortion rate during elongation. Instead, our results are most consistent with a global reduction in the early elongation rate irrespective of codon usage. One intriguing possibility is that the 80*S* ribosome remains engaged with one or more initiation factors during early elongation. In this scenario, the bound initiation factor would maintain the ribosome in a slower state until the factor stochastically dissociates from the ribosome within the first 200 codons. The eIF3 complex is a promising candidate for such a factor, as it binds the solvent-exposed face of the 40*S* ribosome (Siridechadilok et al., 2005) and can therefore bind to 80*S* ribosomes as well (Beznoskova et al., 2013). Maintaining eIF3 on early elongating ribosomes might also facilitate re-initiation after translation of short uORFs (Szamecz et al., 2008).

We also detected stalling of elongating ribosomes after translation of polybasic regions, which presumably interact electrostatically with the ribosome exit tunnel (Figure 3A). Although this stalling had been previously reported for highly charged regions (Brandman et al., 2012), our analyses indicate that this effect is detectable even for stretches with as few as three basic residues. As a result, polybasic-stretch-induced stalling is a surprisingly widespread phenomenon during translation elongation and might impose constraints on protein sequence evolution. In the case of highly charged regions, ribosome stalling has been shown to trigger nascent polypeptide degradation by the proteasome through the ribosome quality control (RQC) complex (Brandman et al., 2012). Whether the thousands of weaker stalling events similarly trigger RQC-mediated polypeptide degradation remains an open question.

### Impact of mRNA enrichment method

A practical finding of our studies is that the choice of mRNA enrichment method can have a significant impact on yeast mRNA-abundance measurements. rRNA depletion using the Ribo-Zero kit emerged as the only method that enriched for mRNAs without introducing substantial and systematic biases (Figure 4A, Figure S10). However, rRNA-depleted samples still contain large amounts of tRNAs, snRNAs, and other noncoding RNAs, which reduces coverage of the mRNA transcriptome. Another caveat of rRNA depletion is that nascent pre-mRNAs that lack a poly(A) tail are also recovered, which can inflate mRNA abundance measurements with respect to the pool of translatable mRNA molecules (although this concern is minimized when using cytoplasmically enriched lysate as the input material for RNA-seq). This effect may be more pronounced in metazoans that contain long introns and correspondingly long transcription times. The extent to which poly(A)-selection biases affect metazoan mRNA abundance data, which would thereby influence TE measurements, remains to be determined.

### Evaluating the range and accuracy of TE measurements

The initial report that TE spans a roughly 100-fold range across mRNAs in budding yeast spurred intensive investigation of the underlying TE determinants, yet there has been minimal success (Kertesz et al., 2010; Robbins-Pianka et al., 2010; Rojas-Duran and Gilbert, 2012; Ouyang et al., 2013; Rouskin et al., 2014). Our results showed that this apparently wide range of TEs is partly explained by inaccurate mRNA-abundance measurements, which comprise the denominators of the TE ratios. After identifying and minimizing this source of inaccuracy, we observed a more narrow range of TEs and IEs (Figure 5A, Figure S12), suggesting a more limited degree of translational control than previously reported (Csárdi et al. 2015).

The TE range that we observed in our experiments in yeast was more similar to the range observed in mouse embryonic stem cells (Ingolia et al., 2011), suggesting that limited translational control is a general principle of gene regulation in rapidly dividing eukaryotic cells. Notably, a study in mouse NIH3T3 cells reached the opposite conclusion: that protein abundance is predominantly controlled at the level of translation (Schwanhausser et al., 2011). However, reanalysis of the NIH3T3 data with more rigorous accounting for experimental error has raised doubts about the validity of that conclusion (Li et al., 2014b).

Although the range of TEs we identify is smaller than that reported earlier, it is nonetheless sufficient for a cell to tune protein synthesis levels. Indeed, IE differences might contribute to the remarkably proportional synthesis of subunits in multiprotein complexes in yeast (Figure 6). However, the potential contribution to proportional synthesis of subunits was lower than that observed in bacteria (Li et al., 2014a), presumably because the absence of operons in eukaryotes uncouples mRNA abundances across the transcriptome, reducing the dependence on translational control.

The weak agreement between our TEs and published TEs (Table S2) suggest that care should be taken in interpreting analyses of published TEs. For example, the previous inability to identify strong correlates of yeast TEs was primarily due to the TEs themselves. Importantly, however, analyses of how the TE of a gene changes across conditions (e.g., during a stress response or a developmental program) are less prone to the biases that we have identified, as gene-specific biases in RNA-seq measurements cancel out when taking the ratio of TEs for the same gene in two different conditions.

### Mechanisms by which structure and length affect yeast TEs

Using our IE measurements, we were able to generate a statistical model that explained a majority of the IE variance (Figure 7D, Table S3). One major finding is that secondary structure within the 5′ UTR appears to be an important determinant of IE. These results are in agreement with mechanistic studies demonstrating that cap accessibility correlates with initiation efficiency (Godefroy-Colburn et al., 1985) and that stable 5′-UTR secondary structures block the scanning ribosome. One caveat of our structure analyses is that we used *in silico* prediction of mRNA structure, which does not always accurately capture the *in vivo* structure of mRNA (Rouskin et al., 2014). Further indicating the inadequacy of *in silico* predictions was the benefit of also including 5′-UTR GC content as a feature in our model. Therefore, mRNA structure presumably explains even more variation in IE than our analyses suggest. Future work will be required to modify the *in vivo* genome-wide structure probing method (DMS-seq) to be able capture the 5′ UTRs of mRNAs, which are depleted in the current protocol (Rouskin and Weissman, personal communication).

We also find that longer ORFs tend to be more poorly translated in log-phase yeast, even after accounting for the small 5′ ramp (Figure 7C). Given that initiation occurs at the 5′ ends of mRNAs, how might initiation rates be sensitive to ORF lengths? One possibility is that shorter mRNAs, which include ribosomal proteins and other housekeeping genes (Hurowitz and Brown, 2003), might be under selection for faster initiation rates by virtue of their high expression. However, our stepwise regression showed that ORF length was informative even after accounting for mRNA abundance. Another intriguing possibility is that the 5′-UTR-bound initiation machinery can sense and be affected by ORF length via the closed-loop structure. In eukaryotes, translating mRNAs are thought to adopt a pseudo-circularized structure in which the 5′- and 3′-ends are in close proximity, enhancing translation and mRNA stability (Christensen et al., 1987). This closed-loop conformation is stabilized by the scaffold protein eIF4G, which can simultaneously interact with cap-bound eIF4E and poly(A)-tail-bound PABP (Tarun and Sachs, 1996; Wells et al., 1998). Previous biochemical analysis of the closed loop in yeast extracts revealed that only short mRNAs adopt a stable closed-loop structure *in vitro* (Amrani et al., 2008), presumably due to the relatively short distance between the mRNA termini. If the same principle applies *in vivo*, then inefficient closed-loop formation of long mRNAs could explain their relatively low IEs. Experimental evidence for this model comes from the observation that depletion of eIF4G, which would disrupt the closed loop, disproportionately reduces the TEs of shorter genes compared to longer genes (Clarkson et al., 2010; Park et al., 2011). However, direct evidence that closed-loop formation is affected by mRNA length and contributes to the IE–length correlation awaits the ability to assay closed loops genome-wide *in vivo*.

## Materials and Methods

### Yeast culture, harvesting, and lysate preparation

*S. cerevisiae* strain BY4741 (*MATa his3Δ1 leu2Δ0 met15Δ0 ura3Δ0*) was grown at 30° C in 500 ml YPD to OD_600_ 0.5. Cells were harvested by filtration onto a Supor 450 Membrane 0.45 μm Disc Filter (Pall #60206) that had been pre-wet with YPD using a Kontes Ultra-Ware Microfiltration Assembly (Kimble Chase Kontes #953825-0090). As the last liquid flowed through, the filtration apparatus was rapidly disassembled, cells were gently scraped off of the filter using a cell lifter (Corning #3008), and the scraper was immediately submerged in a 50-ml conical tube filled with liquid nitrogen. Once all of the liquid nitrogen had boiled off, the resulting pellet of yeast cells was stored in the conical tube at –80° C until lysis.

To lyse cells under cryogenic conditions, the cell pellet was transferred into a pre-chilled mortar that was surrounded and filled with liquid nitrogen. The pellet was ground to a fine powder with a pre-chilled pestle, transferred into a 50-ml conical tube filled with liquid nitrogen, and after boiling off the liquid stored at – 80° C.

Crude lysate was prepared by briefly thawing the cell powder on ice for 1 min and then resuspending in 4 ml Polysome Lysis Buffer (10 mM Tris-HCl pH 7.4, 5 mM MgCl_2_, 100 mM KCl, 1% Triton X-100). The lysate was spun at 1300*g* for 10 min, and the supernatant was flash frozen in single-use aliquots.

### RNA-seq

Total RNA was extracted from an aliquot of frozen yeast lysate using TRI reagent (according to the manufacturer’s protocol). The same total RNA was subjected to the different mRNA enrichment methods as follows:

*Total RNA*: 500 ng of RNA was immediately subjected to ethanol precipitation, as described below.

*Dynabeads oligo(dT)*_*25*_: Poly(A) selection was performed using 100 μl Dynabeads oligo(dT)_25_ (Life Technologies) according to the manufacturer’s instructions, except that 30 μg of total RNA was used as starting material.

*RiboMinus*: rRNA depletion was performed on 4 μg total RNA using the RiboMinus Yeast Transcriptome Isolation Kit (Life Technologies) according to the manufacturer’s instructions.

*Ribo-Zero Magnetic Gold*: rRNA depletion was performed on 10 μg total RNA using the Ribo-Zero Gold Yeast rRNA Removal Kit (Illumina) according to the manufacturer’s instructions, without the addition of RiboGuard RNase Inhibitor.

Following mRNA enrichment, RNA samples were diluted to 90 μl with nuclease-free water and precipitated with 10 μl 3 M NaCl, 30 μg GlycoBlue (Life Technologies), and 250 μl ethanol. RNA-seq was performed exactly as described in Subtelny et al. (2015) using 5 cycles of PCR.

### Ribosome profiling

Ribosome-protected fragments were isolated from an aliquot of frozen yeast lysate and sequenced on the Illumina HiSeq platform, as described in Subtelny et al. (2014). RNase I treatment was performed using 0.2 U/μL lysate. Subtractive hybridization to remove contaminating rRNA fragments was performed using a mixture of three biotinylated oligonucleotides: 5′- GATCGGTCGATTGTGCACCTC/3Bio/; 5′-CGCTTCATTGAATAAGTAAAG/3Bio/; 5′-GACGCCTTATTCGTATCCATC/3Bio/

### Estimating tRNA abundances from RNA-seq

To estimate tRNA abundances, RNA-seq reads from the Ribo-Zero-treated sample were mapped to annotated *S. cerevisiae* tRNA loci (downloaded from Ensembl) using Bowtie, allowing up to one mismatch and sampling alignments for multiply mapping reads (options -n 1 -l 25 -e 100 -M 1 --best --strata). The total number of reads corresponding to each tRNA anticodon were tallied, and wobble parameters were used to estimate cognate tRNA abundances for individual codons.

---

**Figure S1:**
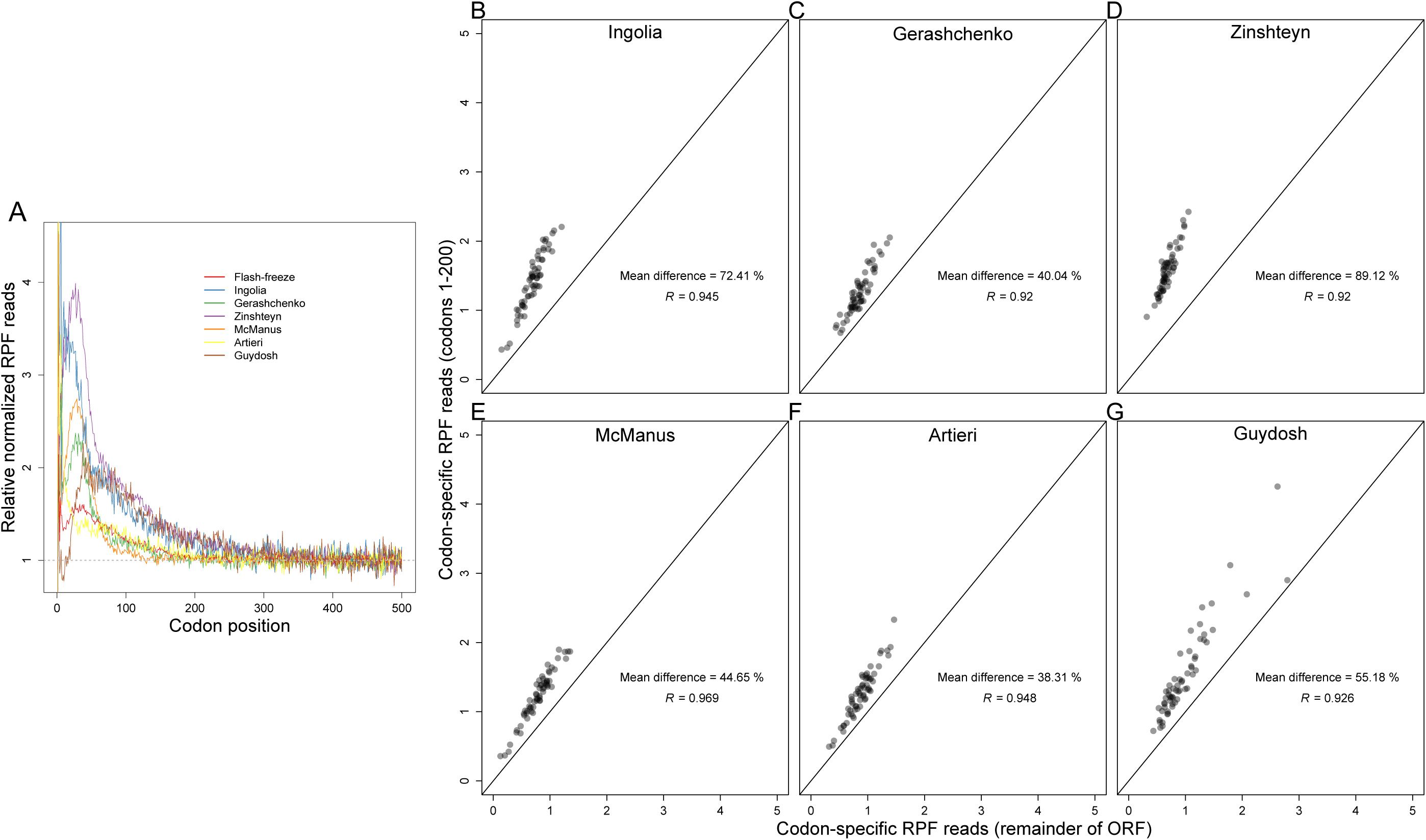
Analyses of the 5’ ramp of ribosomes. (**A**) Metagene analyses of RPF density, performed as in Figure 1D, comparing results of the current study (flash-freeze) to those of published studies. Published datasets are labeled by the first author’s name: Ingolia (INGOLIA et al. 2009), Gerashchenko (GERASHCHENKO et al. 2012), Zinshteyn (ZINSHTEYN and GILBERT 2013), McManus (MCMANUS et al. 2014), Artieri (ARTIERI and FRASER 2014) and Guydosh (GUYDOSH and GREEN 2014). (**B-G**) Comparison of codon-specific RPFs within and beyond the 5’ ramp in six ribosome-profiling experiments. Otherwise, as in Figure 1E.

**Figure S2:**
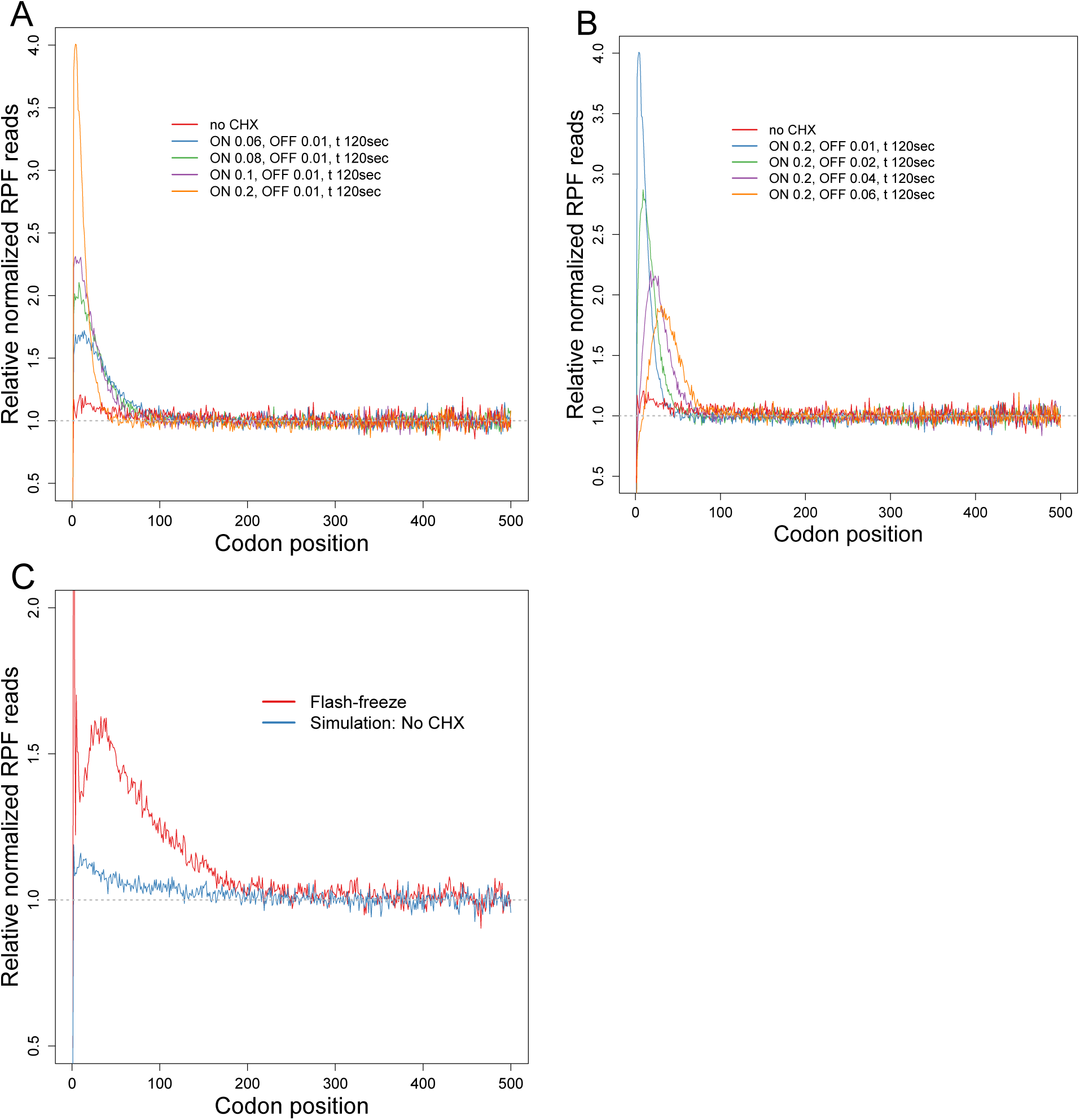
Whole-cell simulations of the 5’ ramp of ribosomes. (**A-B**) Metagene analyses of ribosome-footprint density based on the whole-cell simulation model (SHAH et al. 2013). Simulations with CHX pre-treatment were performed as described in Section 2 of the Supporting File. Simulated RPF reads in ORFs were individually normalized by the mean RPF reads within the ORF, and then averaged with equal weight for each codon position across all ORFs (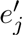 Eqn. S10). We simulated protein synthesis under no-CHX, and either (**A**) four different CHX arrest probabilities (*p*_*chx-on*_) and a fixed CHX dissociation rate (*r*_*chx-off*_), or (**B**) a fixed CHX arrest probability (*p*_*chx*-*on*_) and four different CHX dissociation rates (*r*_*chx*-*off*_). Increasing *p*_*chx*-*on*_ led to higher ramps, whereas increasing *r*_*chx*-*off*_ led to lower ramps as well as a shift of the peak ribosome density towards the 3’ end. (**B-G**) Metagene analysis of RPF density observed in the flash-freeze experiment compared with the results of the whole-cell simulation model. Otherwise, as in (**A**).

**Figure S3:**
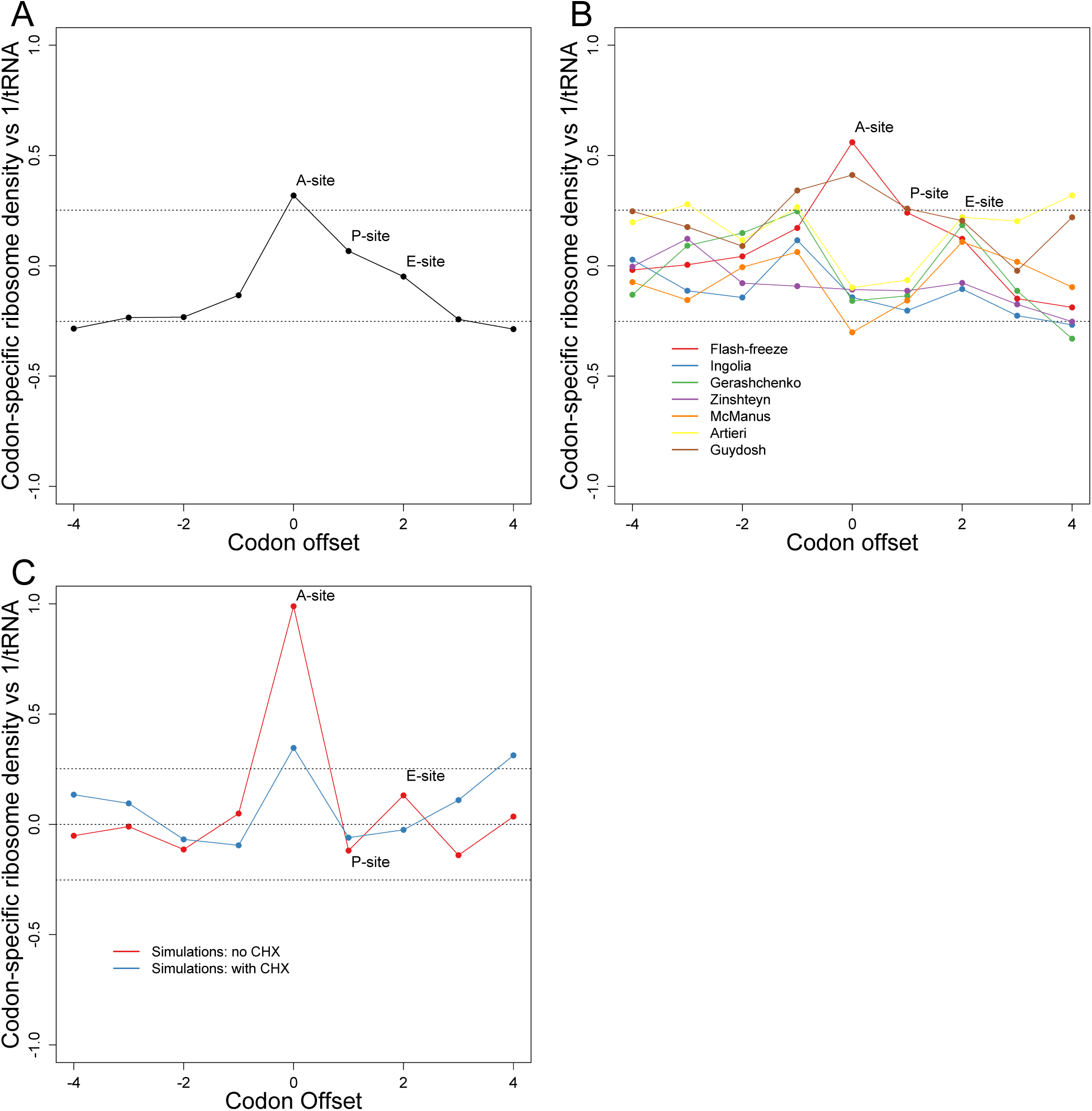
Relationship between cognate tRNA abundances and codon-specific ribosome densities. (**A**) Correlation between codon-specific excess ribosome densities and cognate tRNA abundances estimated using RNA-seq (Table S1). Otherwise, as in Figure 2B. (**B**) Correlation between codon-specific excess ribosome densities and cognate tRNA abundances at various codon offsets, comparing results from the current dataset (flash-freeze) with those of published datasets. Published datasets are labeled by the first author’s name: Ingolia (INGOLIA et al. 2009), Gerashchenko (GERASHCHENKO et al. 2012), Zinshteyn (ZINSHTEYN and GILBERT 2013), McManus (MC-MANUS et al. 2014), Artieri (ARTIERI and FRASER 2014) and Guydosh (GUYDOSH and GREEN 2014). Otherwise, as in Figure 2B. (**C**) Correlation between codon-specific excess ribosome densities and cognate tRNA abundances in simulations with (blue) and without (red) CHX pretreatment. Otherwise, as in Figure 2B.

**Figure S4:**
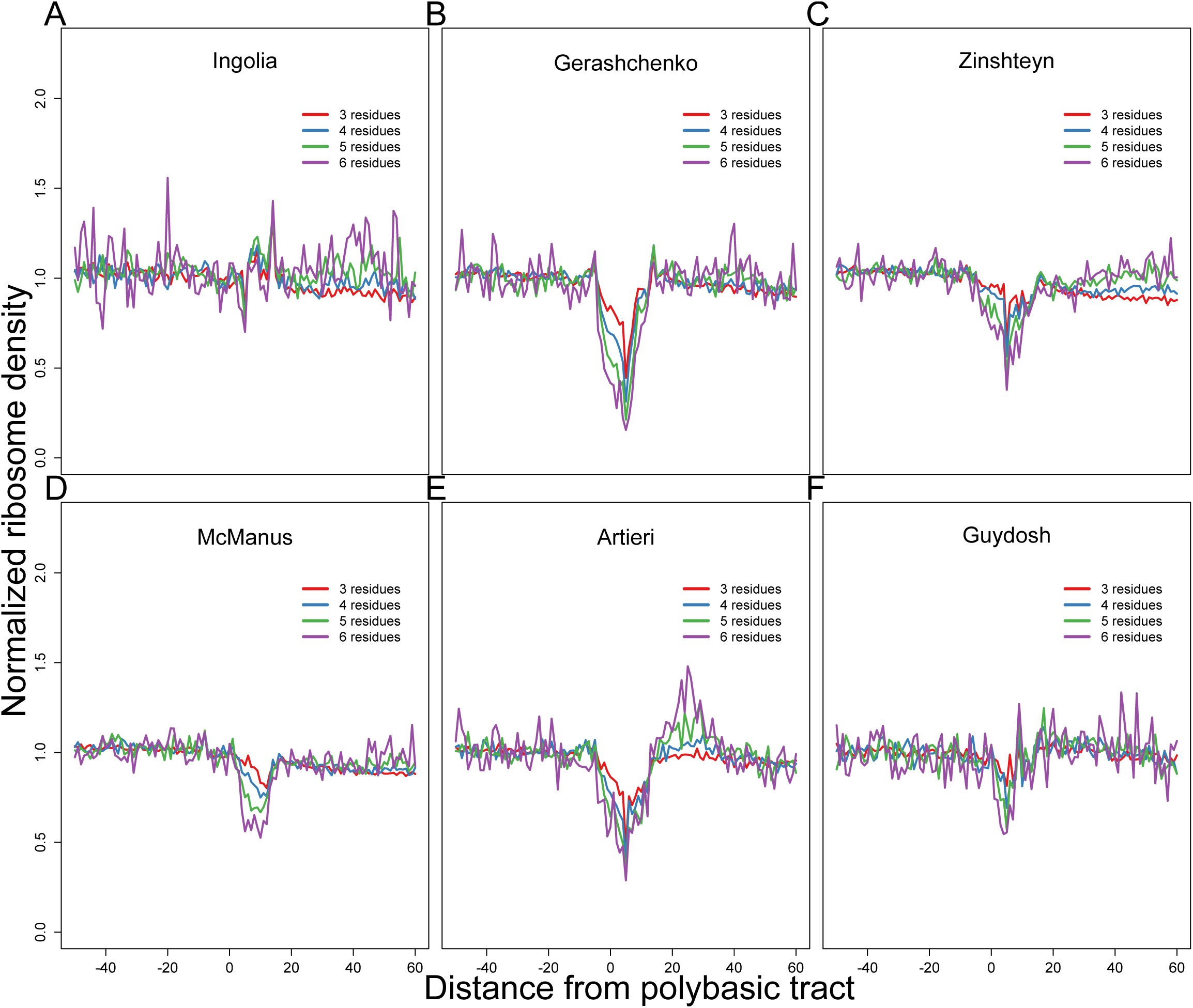
Analyses of ribosome stalling near polybasic tracts. (**A-F**) Metagene analysis of nor-malized ribosome density surrounding polybasic tracts in six published ribosome-profiling experiments. Published datasets are labeled by the first author’s name: Ingolia (INGOLIA et al. 2009), Gerashchenko (GERASHCHENKO et al. 2012), Zinshteyn (ZINSHTEYN and GILBERT 2013), McManus (MCMANUS et al. 2014), Artieri (ARTIERI and FRASER 2014) and Guydosh (GUYDOSH and GREEN 2014). Otherwise, as in Figure 3A.

**Figure S5:**
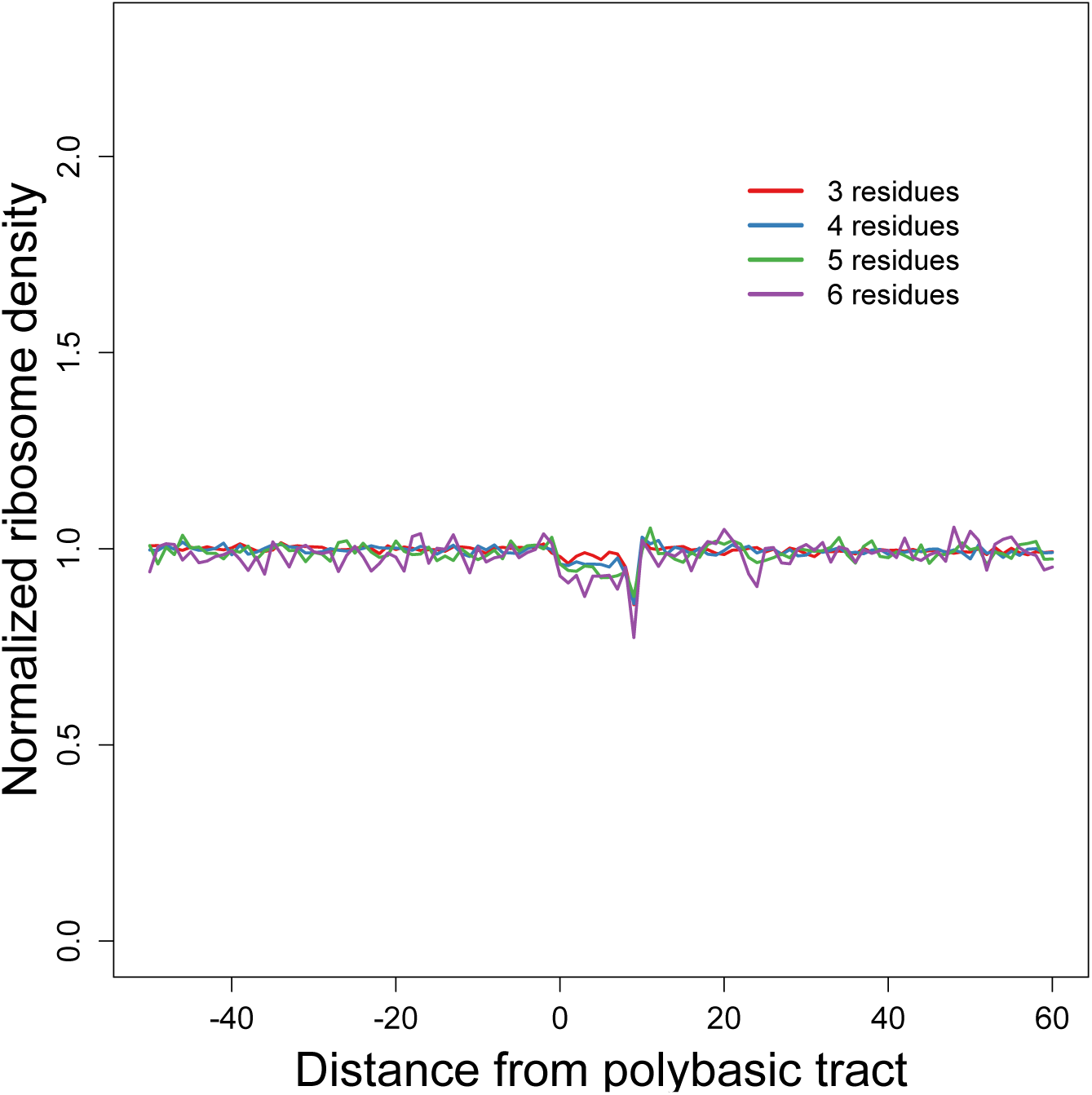
Whole-cell simulation of ribosome stalling near polybasic tracts. Metagene analysis of simulated normalized ribosome-footprint density surrounding polybasic tracts. Otherwise, as in Figure 3A.

**Figure S6:**
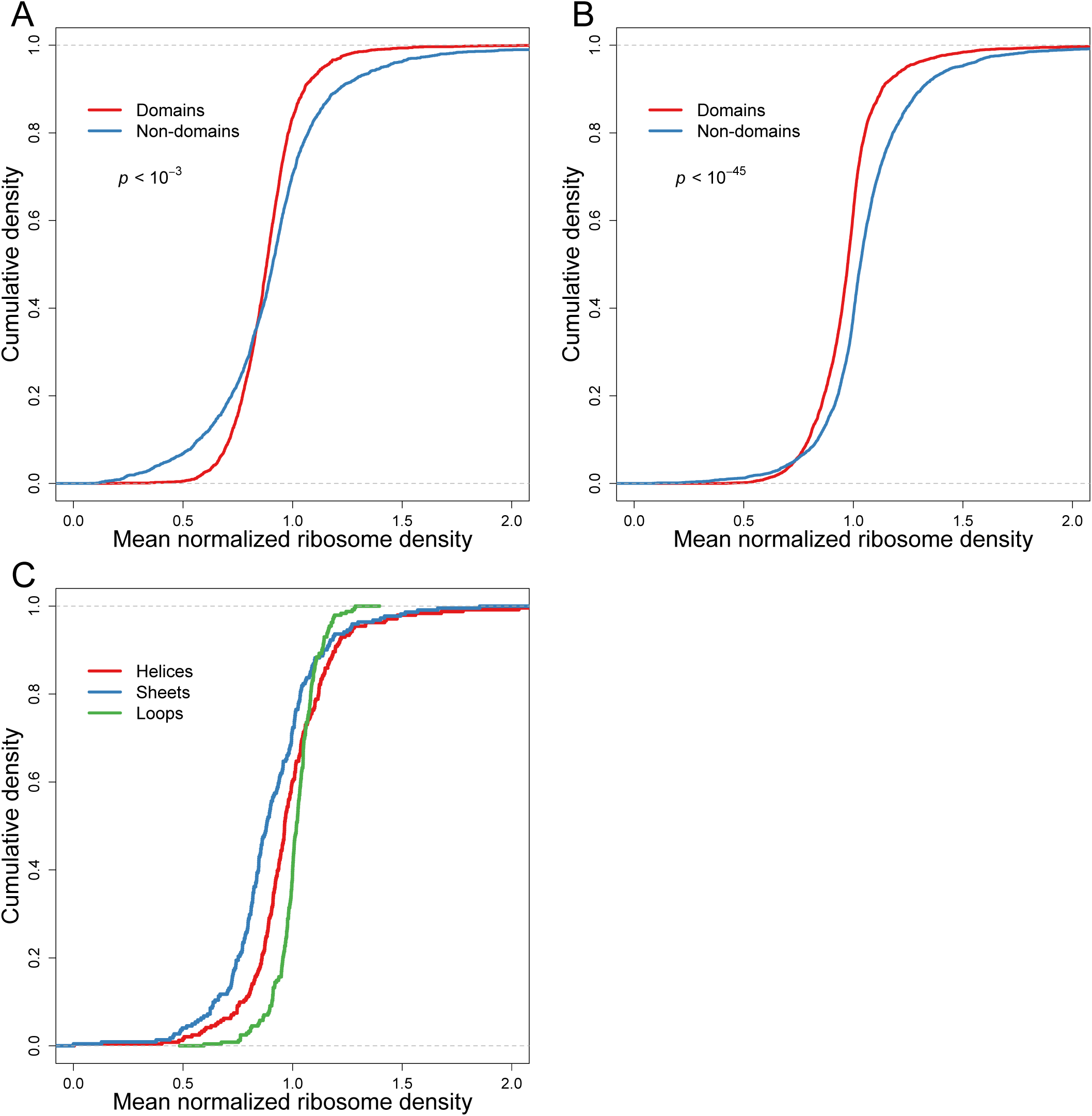
Relationship between elongation dynamics and either domain architecture or protein secondary structure. (**A**) Cumulative distributions of normalized ribosome densities within and outside of protein-folding domains, considering ORFs with at least 250 codons but ignoring the first 200 codons in each ORF. Otherwise, as in Figure 3B. (**B**) Cumulative distributions of normalized ribosome densities within and outside of protein-folding domains. Domain assignments were based on InterProScan classifications (JONES et al. 2014) using the Pfam database (BATEMAN et al. 2002). Otherwise, as in Figure 3B. (**C**) Cumulative distributions of normalized ribosome densities within the indicated classes of protein secondary structures (helices, sheets and loops). Secondary structure assignments for proteins were obtained from (PECHMANN and FRYDMAN 2013). Otherwise, as in Figure 3B.

**Figure S7:**
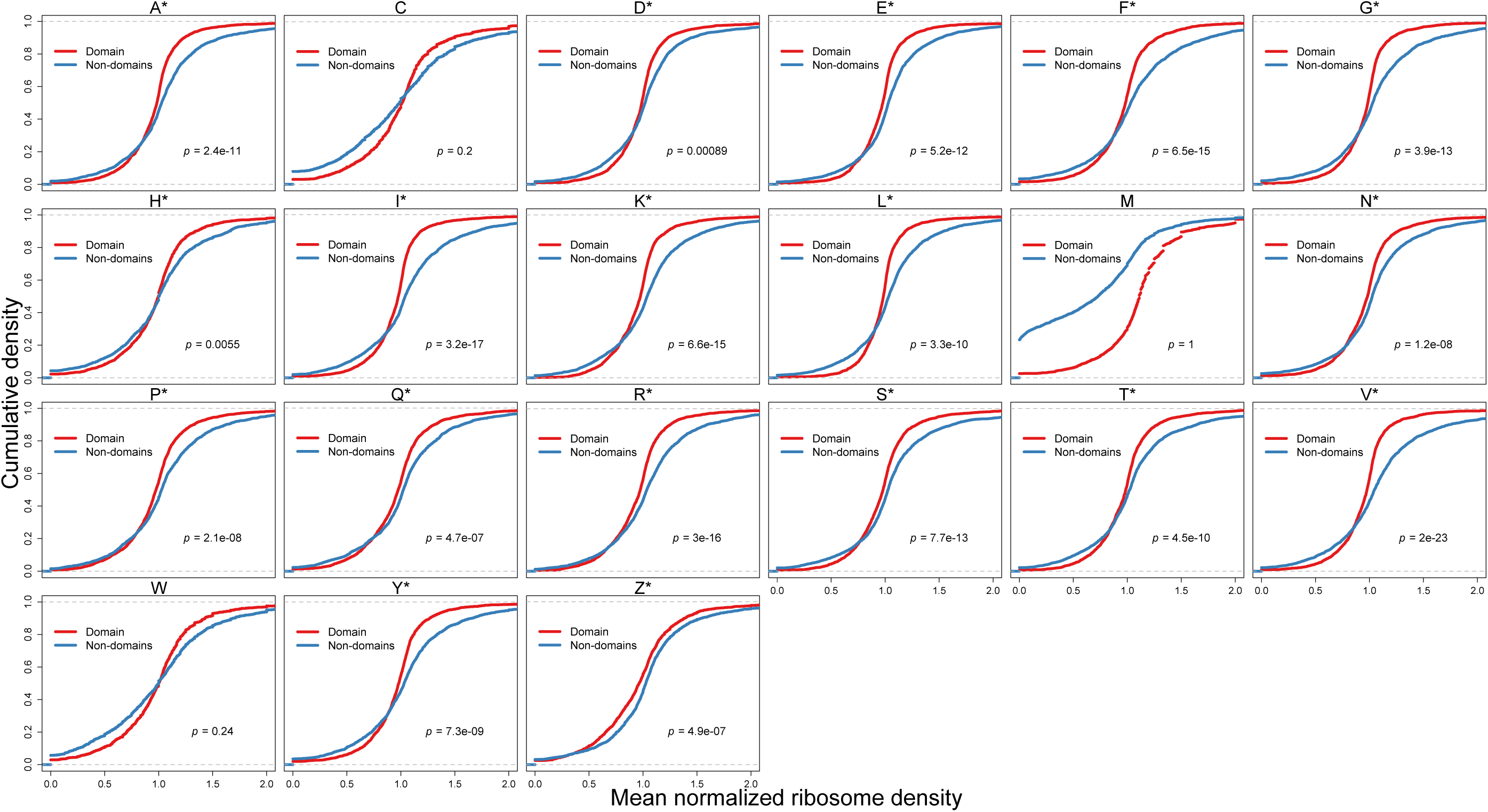
Relationship between elongation dynamics and domain architecture for individual amino acids. Cumulative distributions of normalized ribosome densities within and outside of protein-folding domains, analyzed for each amino acid. Mean normalized ribosome densities (*z*_*i,j*_, Eqn. S7) for codons within the domain-encoding and non-domain-encoding regions were individually calculated for each ORF. Panels are labeled with the single-letter code for each amino acid. Serine codons are partitioned into two sets with 4 and 2 codons (S and Z, respectively). Statistically significant (*p* < 0.05) slowing outside of protein-folding domains is indicated (asterisk). Otherwise, as in Figure 3B.

**Figure S8:**
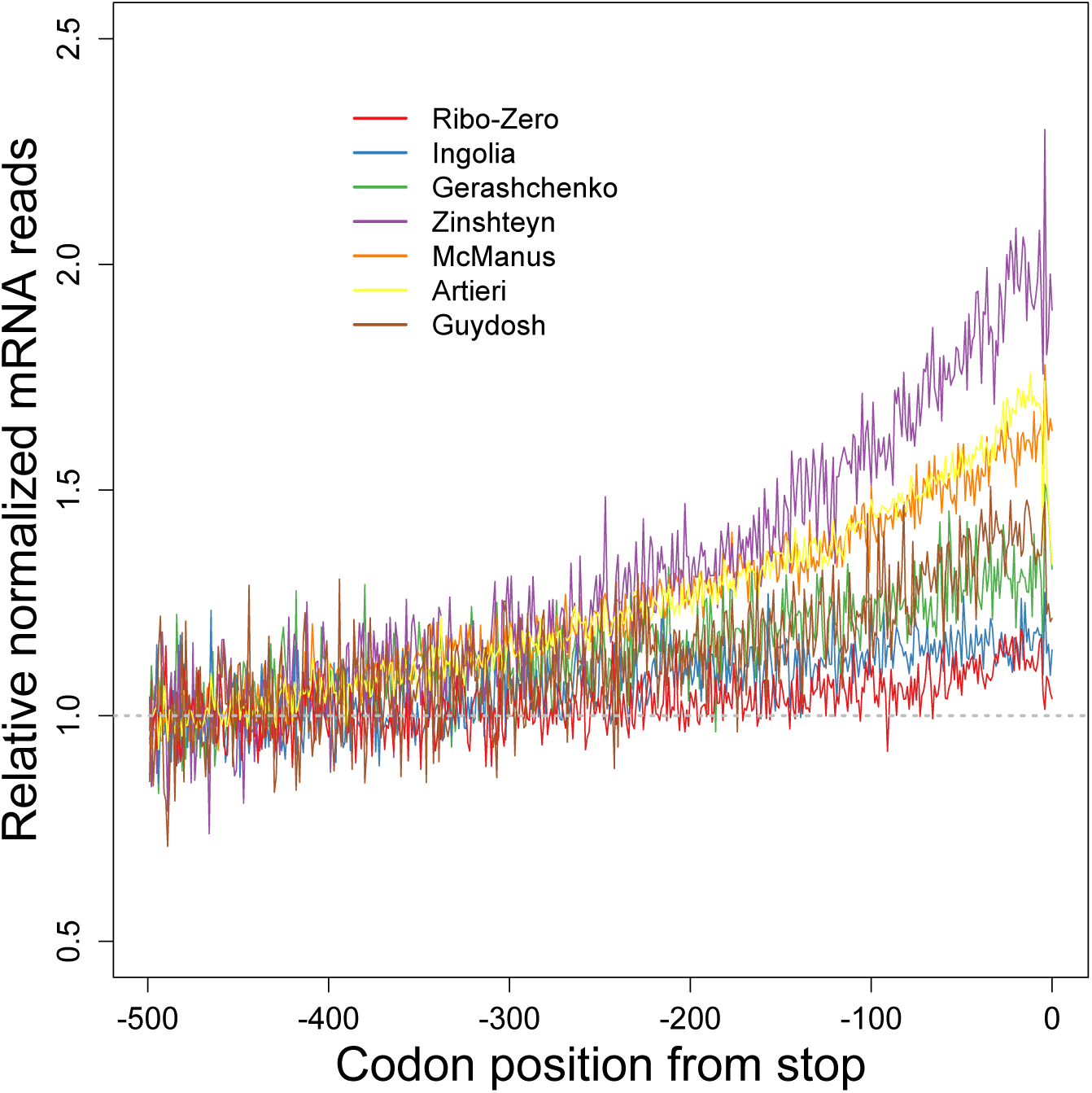
3’ bias observed in RNA-seq datasets. Metagene analysis of RNA-seq read density in six published ribosome-profiling experiments. Otherwise, as in Figure 4B.

**Figure S9:**
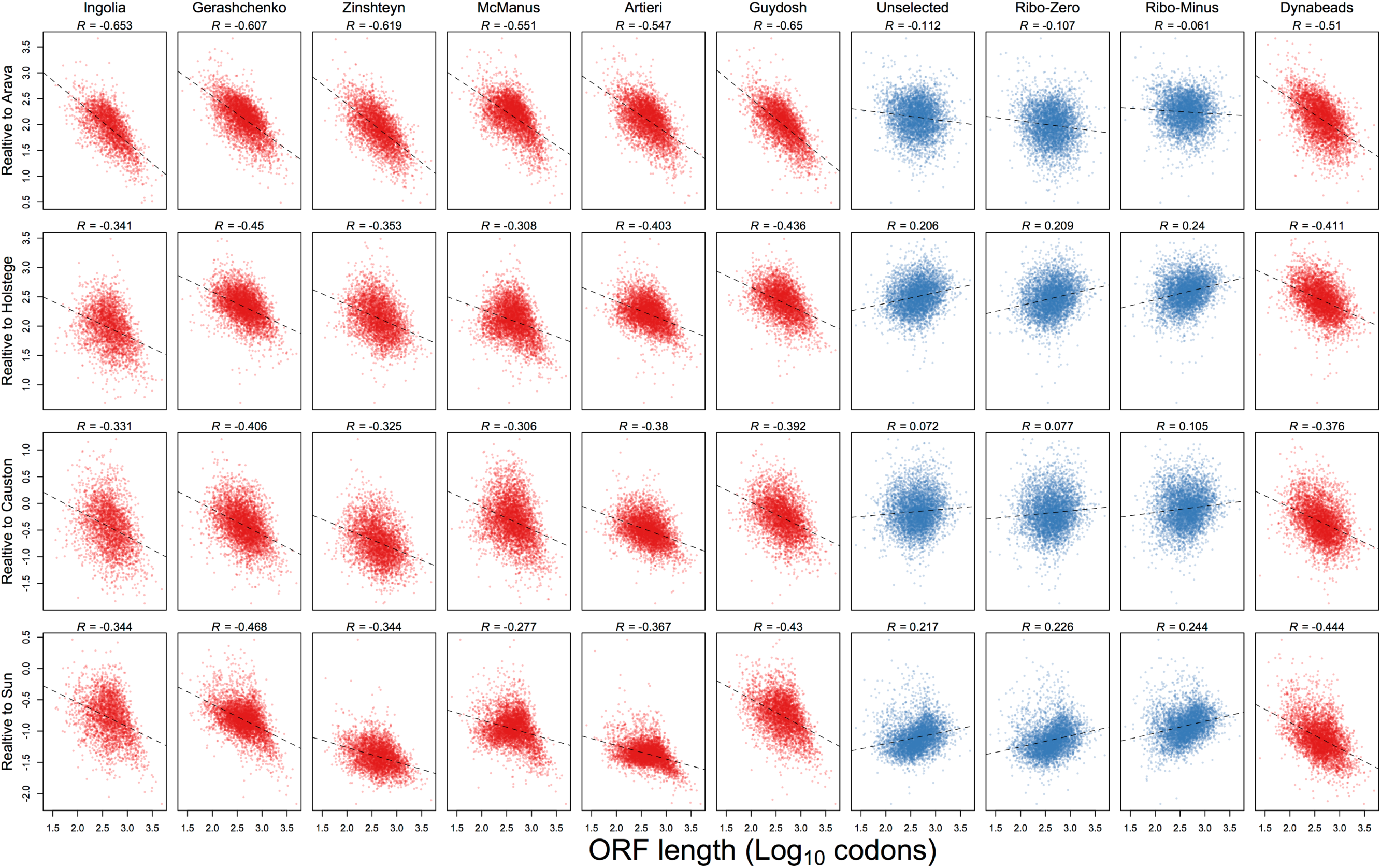
Analysis of the 3’ bias in RNA-seq datasets. Plotted for each analyzed gene is the Log_10_-transformed ratio of mRNA abundance (RPKM) from the RNA-seq of the indicated ribosome-profiling experiment relative to its abundance measured in the indicated microarray dataset, as a function of ORF length. Each column compares mRNA abundances from the same ribosome-profiling dataset, which is labeled by the first author’s name: Ingolia (INGOLIA et al. 2009), Gerashchenko (GERASHCHENKO et al. 2012), Zinshteyn (ZINSHTEYN and GILBERT 2013), McManus (MCMANUS et al. 2014), Artieri (ARTIERI and FRASER 2014) and Guydosh (GUYDOSH and GREEN 2014). Each row represents mRNA abundances from the same microarray dataset, which is labeled by the first author’s name: Arava (ARAVA 2003), Holstege (HOLSTEGE et al. 1998), Causton (CAUSTON et al. 2001) and Sun (SUN et al. 2012). The best linear least-squares fit to the data is shown (dashed line), with the Pearson R.

**Figure S10:**
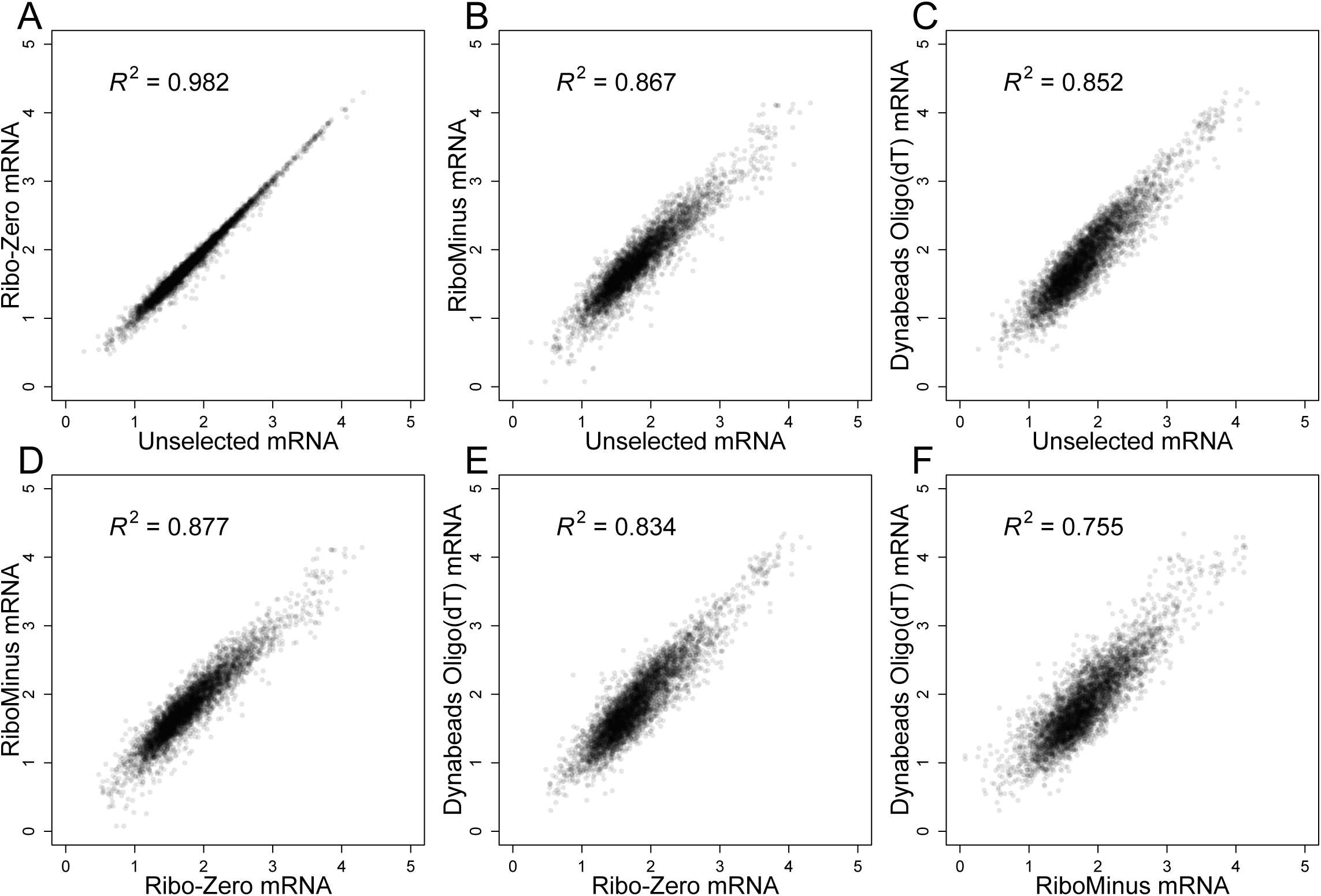
The influence of mRNA enrichment methods on mRNA abundance measurements. (**A-F**) Pairwise comparisons of mRNA abundances (Log_10_ RPKM) following mRNA enrichment by the indicated methods. Otherwise, as in Figure 4A. For comparison, Figure 4A is repeated as panel A.

**Figure S11:**
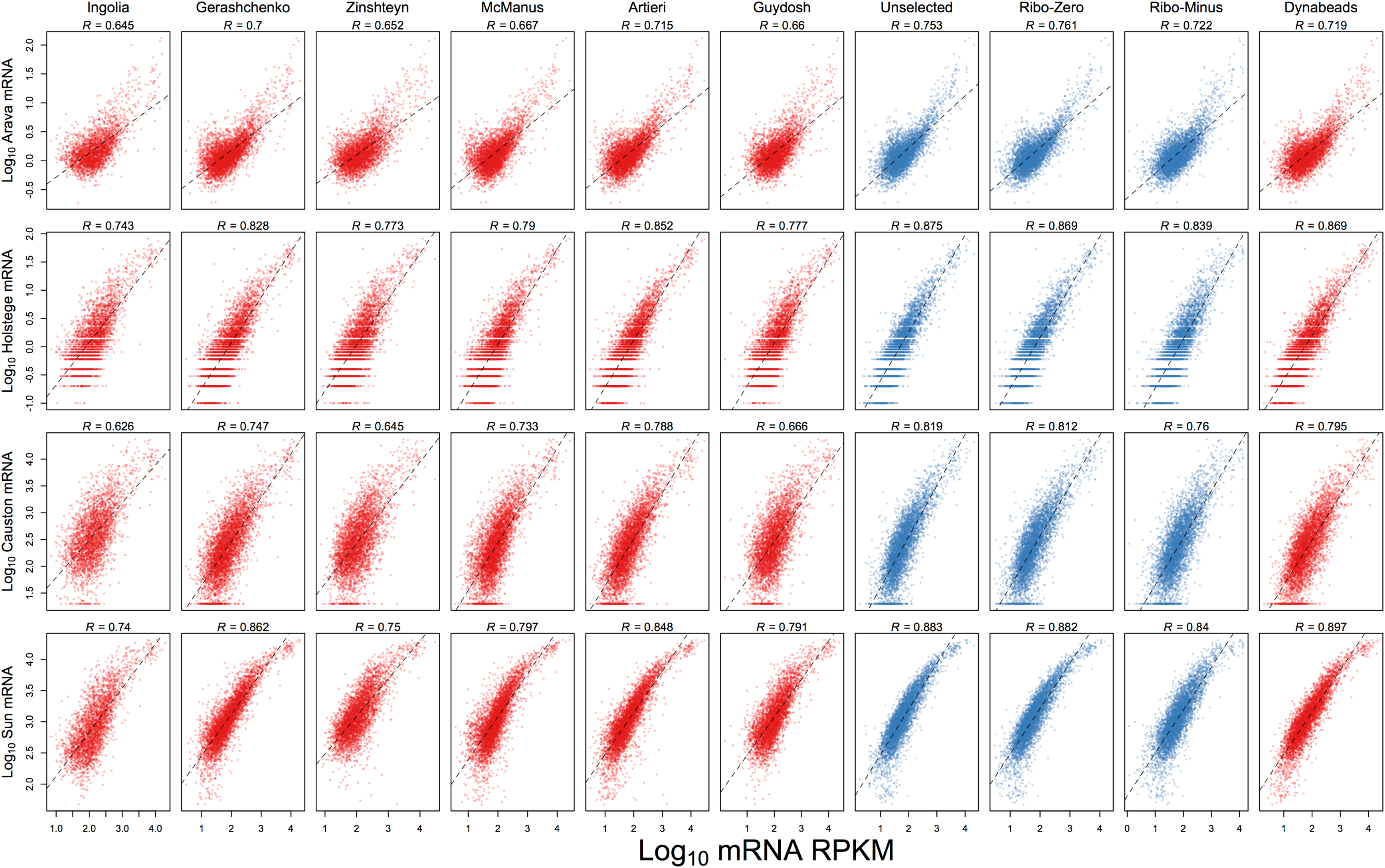
mRNA abundances from microarray studies compared to those measured in the RNA-seq of ribosome-profiling studies. mRNA abundances measured by the indicated microarray dataset are compared to those measured by the indicated ribosome-profiling dataset. Columns and rows are as in Figure S9. The best linear least-squares fit to the data is shown (dashed line), with the Pearson *R*.

**Figure S12:**
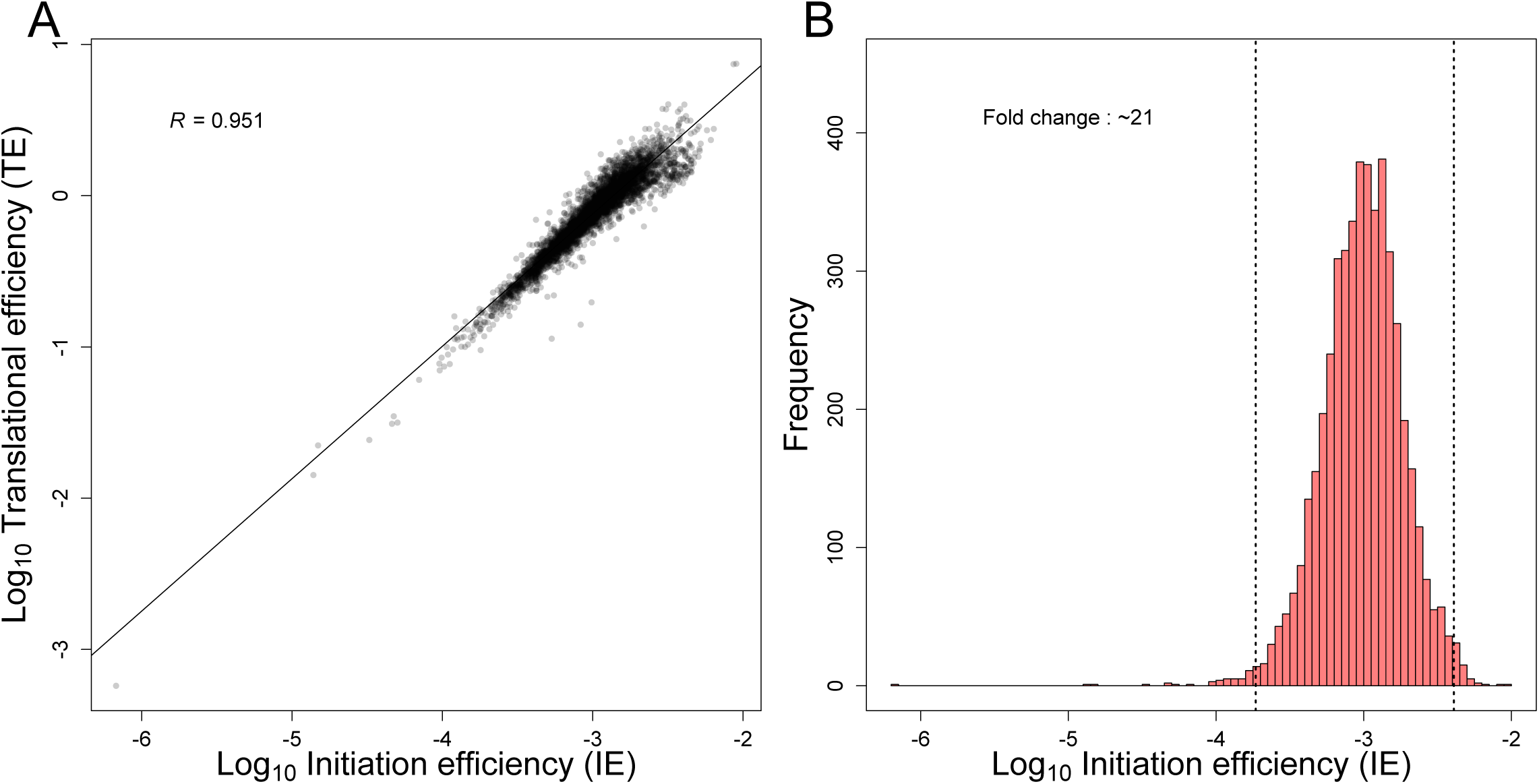
Analyses of IEs in log-phase yeast cells. (A) Relationship between TE values (*TE*, Eqn. S6) and IE values (*p*^*E*^, Eqn. S27). (B) Distribution of IE values. Otherwise, as in Figure 5A.

**Figure S13:**
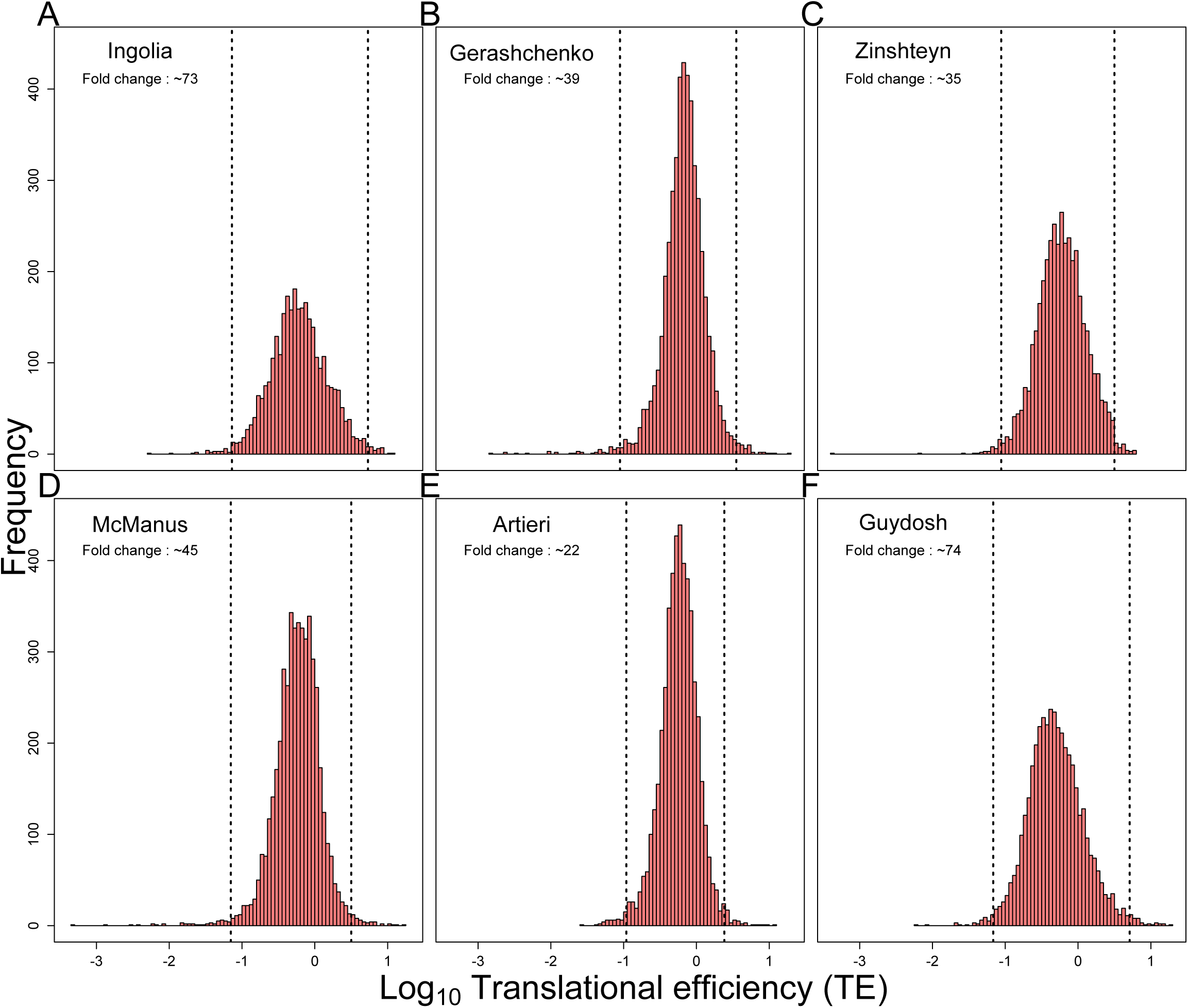
TE distributions from previous ribosome-profiling studies. (**A-F**) Distributions of TE measurements from six published ribosome-profiling experiments. Ribosome-profiling datasets are labeled by the first author’s name: Ingolia (INGOLIA et al. 2009), Gerashchenko (GERASHCHENKO et al. 2012), Zinshteyn (ZINSHTEYN and GILBERT 2013), McManus (MC-MANUS et al. 2014), Artieri (ARTIERI and FRASER 2014) and Guydosh (GUYDOSH and GREEN 2014). Otherwise, as in Figure 5A.

**Figure S14:**
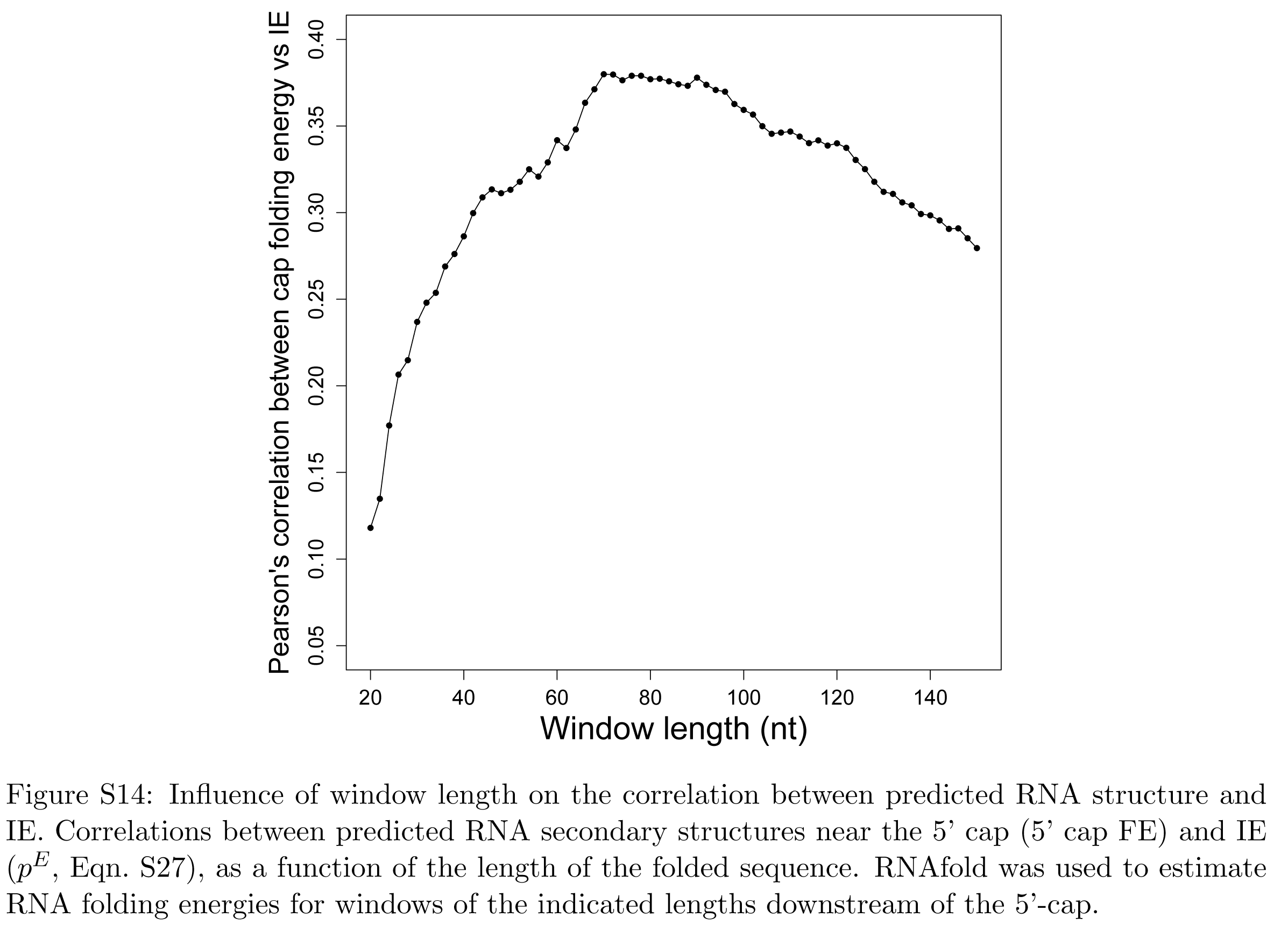
Influence of window length on the correlation between predicted RNA structure and IE. Correlations between predicted RNA secondary structures near the 5’ cap (5’ cap FE) and IE (*p*^*E*^, Eqn. S27), as a function of the length of the folded sequence. RNAfold was used to estimate RNA folding energies for windows of the indicated lengths downstream of the 5’-cap.

**Figure S15:**
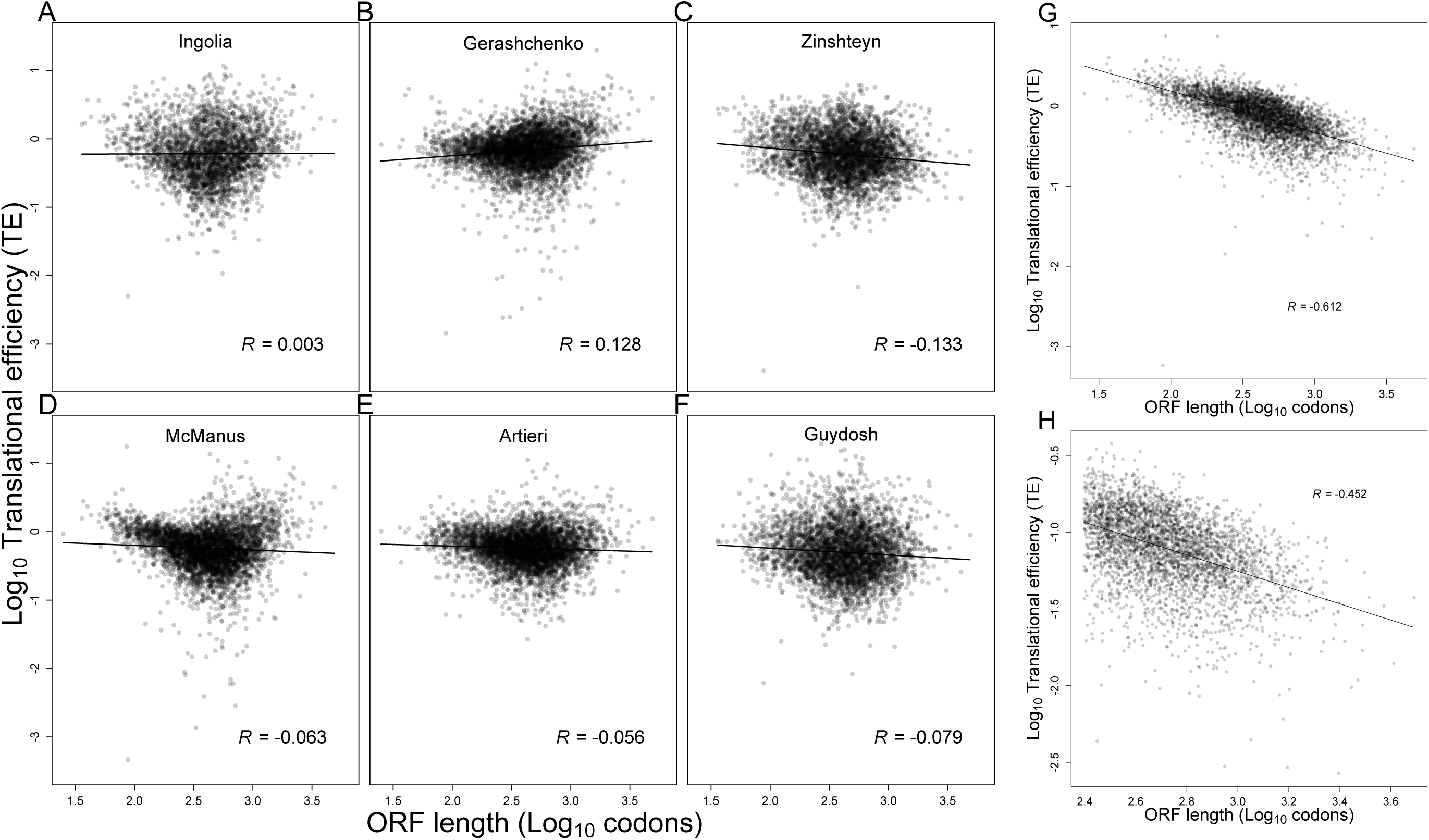
Analyses of the relationship between gene length and TE. (**A-F**) Relationship between TE and ORF length in published ribosome-profiling datasets. Ribosome-profiling datasets are labeled by the first author’s name: Ingolia (INGOLIA et al. 2009), Gerashchenko (GERASHCHENKO et al. 2012), Zinshteyn (ZINSHTEYN and GILBERT 2013), McManus (MCMANUS et al. 2014), Artieri (ARTIERI and FRASER 2014) and Guydosh (GUYDOSH and GREEN 2014). The best linear least-squares fit to the data is shown (line), with the Pearson *R*. (**G**) Relationship between TE and ORF length observed when analyzing the flash-freeze dataset. Otherwise, as in (**A**). (**H**) Relationship between TE and ORF length observed after excluding data from within the 5’ ramp. Only genes with 250 codons were considered, and RPF and RNA-seq reads mapping to the first 200 codons of each ORF were excluded. Otherwise, as in (**G**).

**Figure S16:**
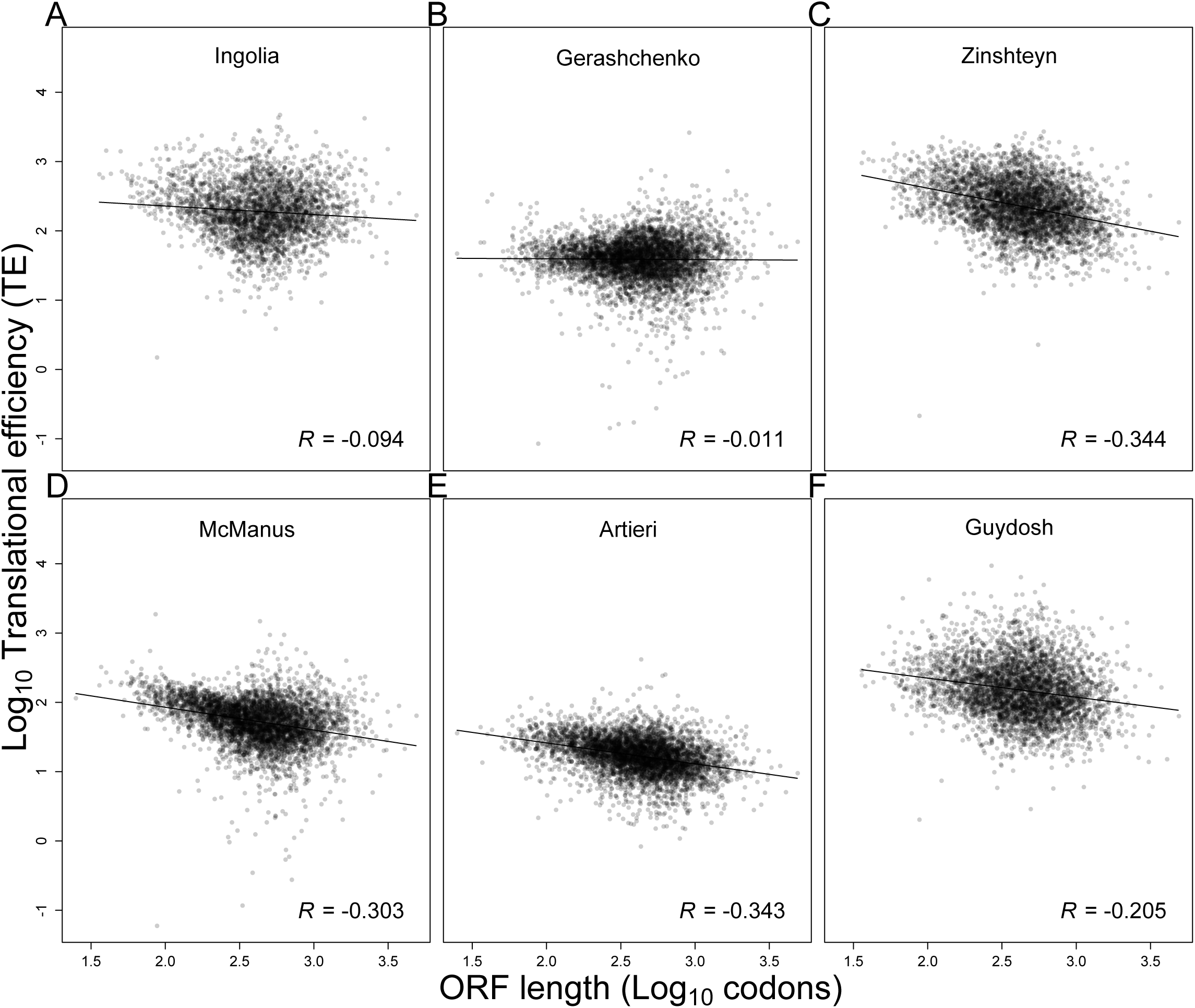
Relationship between TE and ORF length after correcting for the 3’ bias in RNA-seq reads. (**A-F**) Relationship between TE and ORF length after correcting for the 3’ bias observed in the RNA-seq reads of published ribosome-profiling experiments. For each ORF, the mRNA RPKM was calculated based only on reads mapping to the last 200 codons. Otherwise, as in Figure S15.

**Table S1:**

| **Pearson's correlation** | *(tRNA GCN \* Wobble)* | *tRNA abundance*<br>*(Microarray \* Wobble)* | *tAI* | *RiboDensity at*<br>*A-site* |
| --- | --- | --- | --- | --- |
| *tRNA abundance (RNA-seq \* Wobble)* | 0.748 | 0.608 | 0.713 | 0.268 |
| *(tRNA GCN \* Wobble)* |  | 0.778 | 0.947 | 0.512 |
| *tRNA abundance*<br>*(Microarray \* Wobble)* |  |  | 0.713 | 0.459 |
| *tAI* |  |  |  | 0.509 |

| **Spearman's correlation** | *(tRNA GCN \* Wobble)* | *tRNA abundance*<br>*(Microarray \* Wobble)* | *tAI* | *RiboDensity at*<br>*A-site* |
| --- | --- | --- | --- | --- |
| *tRNA abundance (RNA-seq \* Wobble)* | 0.777 | 0.605 | 0.752 | 0.319 |
| *(tRNA GCN \* Wobble)* |  | 0.785 | 0.917 | 0.560 |
| *tRNA abundance*<br>*(Microarray \* Wobble)* |  |  | 0.711 | 0.463 |
| *tAI* |  |  |  | 0.532 |

**Table S2:**

| **RPF** | **Ingolia** | **Gerashchenko** | **Zinshteyn** | **McManus** | **Artieri** | **Guydosch** |
| --- | --- | --- | --- | --- | --- | --- |
| **Flash-Freeze** | 0.982 | 0.972 | 0.971 | 0.973 | 0.972 | 0.98 |
| **Ingolia** |  | 0.965 | 0.98 | 0.961 | 0.962 | 0.979 |
| **Gerashchenko** |  |  | 0.945 | 0.966 | 0.952 | 0.961 |
| **Zinshteyn** |  |  |  | 0.966 | 0.96 | 0.973 |
| **McManus** |  |  |  |  | 0.967 | 0.958 |
| **Artieri** |  |  |  |  |  | 0.971 |

| **mRNA** | **Ingolia** | **Gerashchenko** | **Zinshteyn** | **McManus** | **Artieri** | **Guydosch** |
| --- | --- | --- | --- | --- | --- | --- |
| **Flash-Freeze** | 0.686 | 0.844 | 0.749 | 0.816 | 0.883 | 0.77 |
| **Ingolia** |  | 0.884 | 0.936 | 0.849 | 0.842 | 0.971 |
| **Gerashchenko** |  |  | 0.869 | 0.898 | 0.922 | 0.908 |
| **Zinshteyn** |  |  |  | 0.858 | 0.865 | 0.952 |
| **McManus** |  |  |  |  | 0.93 | 0.888 |
| **Artieri** |  |  |  |  |  | 0.886 |

| **TE** | **Ingolia** | **Gerashchenko** | **Zinshteyn** | **McManus** | **Artieri** | **Guydosch** |
| --- | --- | --- | --- | --- | --- | --- |
| **Flash-Freeze** | 0.479 | 0.498 | 0.55 | 0.553 | 0.561 | 0.542 |
| **Ingolia** |  | 0.729 | 0.907 | 0.777 | 0.797 | 0.924 |
| **Gerashchenko** |  |  | 0.633 | 0.774 | 0.757 | 0.674 |
| **Zinshteyn** |  |  |  | 0.673 | 0.731 | 0.873 |
| **McManus** |  |  |  |  | 0.784 | 0.702 |
| **Artieri** |  |  |  |  |  | 0.806 |

**Table S3:**
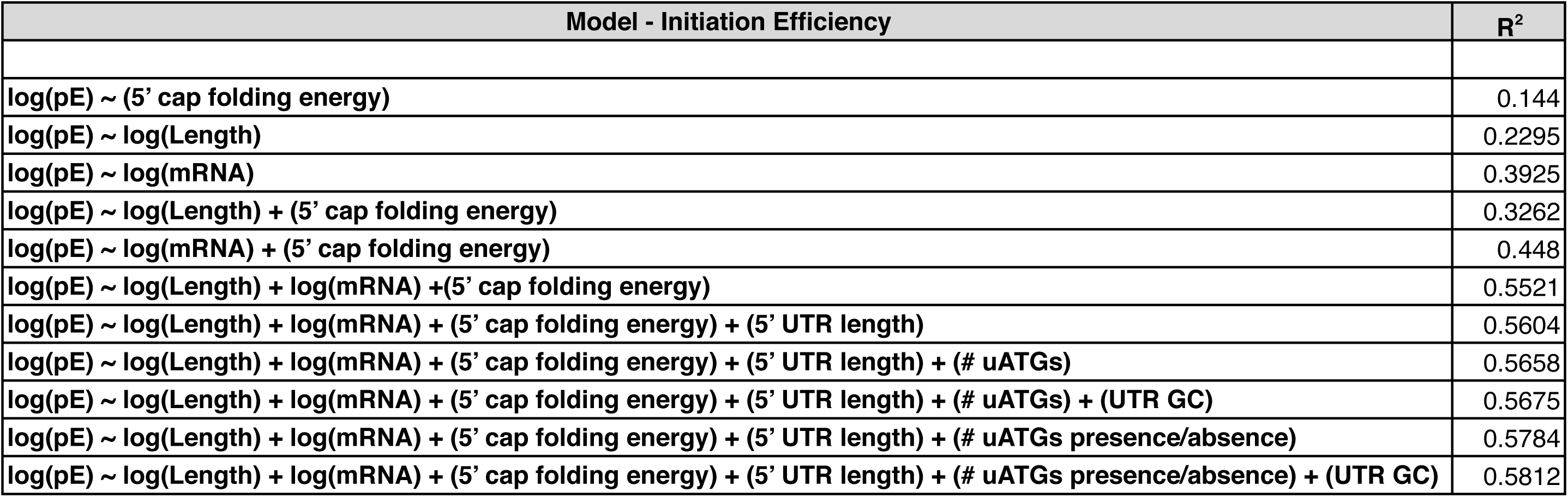

**Table S4:**
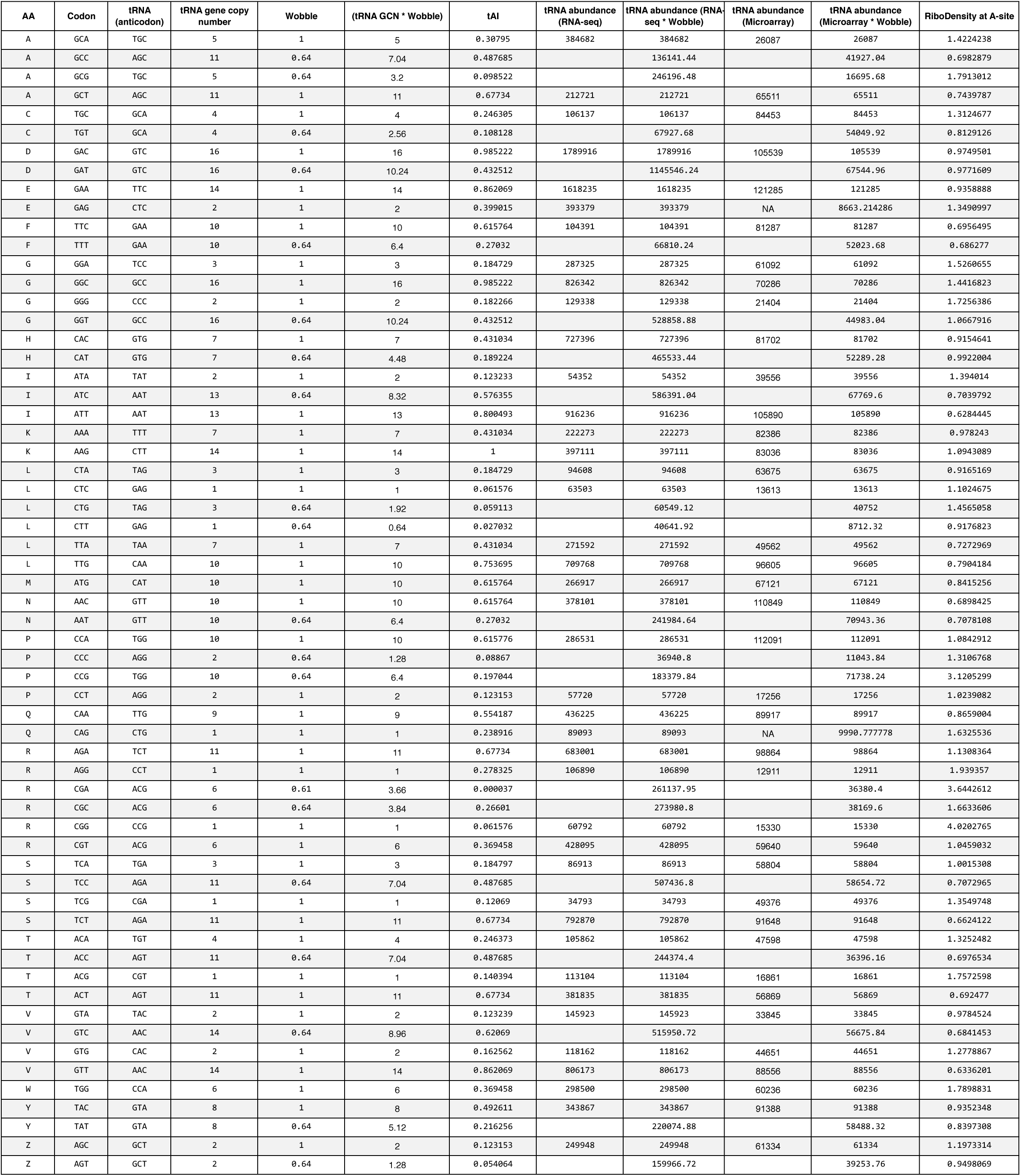

| AA | Codon | tRNA<br>(anticodon) | tRNA gene copy<br>number | Wobble | (tRNA GCN * Wobble) | tAI | tRNA abundance<br>(RNA-seq) | tRNA abundance (RNA-seq * Wobble) | tRNA abundance<br>(Microarray) | tRNA abundance<br>(Microarray * Wobble) | RiboDensity at A-site |
| --- | --- | --- | --- | --- | --- | --- | --- | --- | --- | --- | --- |
| A | GCA | TGC | 5 | 1 | 5 | 0.30795 | 384682 | 384682 | 26087 | 26087 | 1.4224238 |
| A | GCC | AGC | 11 | 0.64 | 7.04 | 0.487685 |  | 136141.44 |  | 41927.04 | 0.6982879 |
| A | GCG | TGC | 5 | 0.64 | 3.2 | 0.098522 |  | 246196.48 |  | 16695.68 | 1.7913012 |
| A | GCT | AGC | 11 | 1 | 11 | 0.67734 | 212721 | 212721 | 65511 | 65511 | 0.7439787 |
| C | TGC | GCA | 4 | 1 | 4 | 0.246305 | 106137 | 106137 | 84453 | 84453 | 1.3124677 |
| C | TGT | GCA | 4 | 0.64 | 2.56 | 0.108128 |  | 67927.68 |  | 54049.92 | 0.8129126 |
| D | GAC | GTC | 16 | 1 | 16 | 0.985222 | 1789916 | 1789916 | 105539 | 105539 | 0.9749501 |
| D | GAT | GTC | 16 | 0.64 | 10.24 | 0.432512 |  | 1145546.24 |  | 67544.96 | 0.9771609 |
| E | GAA | TTC | 14 | 1 | 14 | 0.862069 | 1618235 | 1618235 | 121285 | 121285 | 0.9358888 |
| E | GAG | CTC | 2 | 1 | 2 | 0.399015 | 393379 | 393379 | NA | 8663.214286 | 1.3490997 |
| F | TTC | GAA | 10 | 1 | 10 | 0.615764 | 104391 | 104391 | 81287 | 81287 | 0.6956495 |
| F | TTT | GAA | 10 | 0.64 | 6.4 | 0.27032 |  | 66810.24 |  | 52023.68 | 0.686277 |
| G | GGA | TCC | 3 | 1 | 3 | 0.184729 | 287325 | 287325 | 61092 | 61092 | 1.5260655 |
| G | GGC | GCC | 16 | 1 | 16 | 0.985222 | 826342 | 826342 | 70286 | 70286 | 1.4416823 |
| G | GGG | CCC | 2 | 1 | 2 | 0.182266 | 129338 | 129338 | 21404 | 21404 | 1.7256386 |
| G | GGT | GCC | 16 | 0.64 | 10.24 | 0.432512 |  | 528858.88 |  | 44983.04 | 1.0667916 |
| H | CAC | GTG | 7 | 1 | 7 | 0.431034 | 727396 | 727396 | 81702 | 81702 | 0.9154641 |
| H | CAT | GTG | 7 | 0.64 | 4.48 | 0.189224 |  | 465533.44 |  | 52289.28 | 0.9922004 |
| I | ATA | TAT | 2 | 1 | 2 | 0.123233 | 54352 | 54352 | 39556 | 39556 | 1.394014 |
| I | ATC | AAT | 13 | 0.64 | 8.32 | 0.576355 |  | 586391.04 |  | 67769.6 | 0.7039792 |
| I | ATT | AAT | 13 | 1 | 13 | 0.800493 | 916236 | 916236 | 105890 | 105890 | 0.6284445 |
| K | AAA | TTT | 7 | 1 | 7 | 0.431034 | 222273 | 222273 | 82386 | 82386 | 0.978243 |
| K | AAG | CTT | 14 | 1 | 14 | 1 | 397111 | 397111 | 83036 | 83036 | 1.0943089 |
| L | CTA | TAG | 3 | 1 | 3 | 0.184729 | 94608 | 94608 | 63675 | 63675 | 0.9165169 |
| L | CTC | GAG | 1 | 1 | 1 | 0.061576 | 63503 | 63503 | 13613 | 13613 | 1.1024675 |
| L | CTG | TAG | 3 | 0.64 | 1.92 | 0.059113 |  | 60549.12 |  | 40752 | 1.4565058 |
| L | CTT | GAG | 1 | 0.64 | 0.64 | 0.027032 |  | 40641.92 |  | 8712.32 | 0.9176823 |
| L | TTA | TAA | 7 | 1 | 7 | 0.431034 | 271592 | 271592 | 49562 | 49562 | 0.7272969 |
| L | TTG | CAA | 10 | 1 | 10 | 0.753695 | 709768 | 709768 | 96605 | 96605 | 0.7904184 |
| M | ATG | CAT | 10 | 1 | 10 | 0.615764 | 266917 | 266917 | 67121 | 67121 | 0.8415256 |
| N | AAC | GTT | 10 | 1 | 10 | 0.615764 | 378101 | 378101 | 110849 | 110849 | 0.6898425 |
| N | AAT | GTT | 10 | 0.64 | 6.4 | 0.27032 |  | 241984.64 |  | 70943.36 | 0.7078108 |
| P | CCA | TGG | 10 | 1 | 10 | 0.615776 | 286531 | 286531 | 112091 | 112091 | 1.0842912 |
| P | CCC | AGG | 2 | 0.64 | 1.28 | 0.08867 |  | 36940.8 |  | 11043.84 | 1.3106768 |
| P | CCG | TGG | 10 | 0.64 | 6.4 | 0.197044 |  | 183379.84 |  | 71738.24 | 3.1205299 |
| P | CCT | AGG | 2 | 1 | 2 | 0.123153 | 57720 | 57720 | 17256 | 17256 | 1.0239082 |
| Q | CAA | TTG | 9 | 1 | 9 | 0.554187 | 436225 | 436225 | 89917 | 89917 | 0.8659004 |
| Q | CAG | CTG | 1 | 1 | 1 | 0.238916 | 89093 | 89093 | NA | 9990.777778 | 1.6325536 |
| R | AGA | TCT | 11 | 1 | 11 | 0.67734 | 683001 | 683001 | 98864 | 98864 | 1.1308364 |
| R | AGG | CCT | 1 | 1 | 1 | 0.278325 | 106890 | 106890 | 12911 | 12911 | 1.939357 |
| R | CGA | ACG | 6 | 0.61 | 3.66 | 0.000037 |  | 261137.95 |  | 36380.4 | 3.6442612 |
| R | CGC | ACG | 6 | 0.64 | 3.84 | 0.26601 |  | 273980.8 |  | 38169.6 | 1.6633606 |
| R | CGG | CCG | 1 | 1 | 1 | 0.061576 | 60792 | 60792 | 15330 | 15330 | 4.0202765 |
| R | CGT | ACG | 6 | 1 | 6 | 0.369458 | 428095 | 428095 | 59640 | 59640 | 1.0459032 |
| S | TCA | TGA | 3 | 1 | 3 | 0.184797 | 86913 | 86913 | 58804 | 58804 | 1.0015308 |
| S | TCC | AGA | 11 | 0.64 | 7.04 | 0.487685 |  | 507436.8 |  | 58654.72 | 0.7072965 |
| S | TCG | CGA | 1 | 1 | 1 | 0.12069 | 34793 | 34793 | 49376 | 49376 | 1.3549748 |
| S | TCT | AGA | 11 | 1 | 11 | 0.67734 | 792870 | 792870 | 91648 | 91648 | 0.6624122 |
| T | ACA | TGT | 4 | 1 | 4 | 0.246373 | 105862 | 105862 | 47598 | 47598 | 1.3252482 |
| T | ACC | AGT | 11 | 0.64 | 7.04 | 0.487685 |  | 244374.4 |  | 36396.16 | 0.6976534 |
| T | ACG | CGT | 1 | 1 | 1 | 0.140394 | 113104 | 113104 | 16861 | 16861 | 1.7572598 |
| T | ACT | AGT | 11 | 1 | 11 | 0.67734 | 381835 | 381835 | 56869 | 56869 | 0.692477 |
| V | GTA | TAC | 2 | 1 | 2 | 0.123239 | 145923 | 145923 | 33845 | 33845 | 0.9784524 |
| V | GTC | AAC | 14 | 0.64 | 8.96 | 0.62069 |  | 515950.72 |  | 56675.84 | 0.6841453 |
| V | GTG | CAC | 2 | 1 | 2 | 0.162562 | 118162 | 118162 | 44651 | 44651 | 1.2778867 |
| V | GTT | AAC | 14 | 1 | 14 | 0.862069 | 806173 | 806173 | 88556 | 88556 | 0.6336201 |
| W | TGG | CCA | 6 | 1 | 6 | 0.369458 | 298500 | 298500 | 60236 | 60236 | 1.7898831 |
| Y | TAC | GTA | 8 | 1 | 8 | 0.492611 | 343867 | 343867 | 91388 | 91388 | 0.9352348 |
| Y | TAT | GTA | 8 | 0.64 | 5.12 | 0.216256 |  | 220074.88 |  | 58488.32 | 0.8397308 |
| Z | AGC | GCT | 2 | 1 | 2 | 0.123153 | 249948 | 249948 | 61334 | 61334 | 1.1973314 |
| Z | AGT | GCT | 2 | 0.64 | 1.28 | 0.054064 |  | 159966.72 |  | 39253.76 | 0.9498069 |

## Supplementary Methods

## 1 Simulation model

We use a whole-cell model of protein translation to simulate the dynamics of protein production (Shah et al., 2013). Briefly, the model assumes a fixed total number of ribosomes and tRNAs, and it describes how these entities initiate and elongate a fixed supply of mRNAs.

We define a genome with *n* = 4862 genes, each with a prescribed coding sequence, and fixed mRNA abundance *A*_*i*_. Gene *i* encodes an mRNA of length *L*_*i*_ codons and has a corresponding probability of translation initiation, denoted *p*_*i*_, which is described below.

Each codon of type *j* is decoded by one of 41 iso-accepting tRNA species, denoted *ϕ*(*j*), which has a fixed total abundance 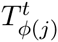 in the cell. Each molecule of tRNA species *ϕ*(*j*) is either free in the cell, or bound, along with a ribosome, to a codon of type *j* on an mRNA in the cell. Thus, the total number of tRNAs of type *ϕ*(*j*) can be decomposed into those that are currently bound and those that are currently free: 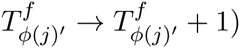. Similarly, the total number of ribosomes, *R*^*t*^, can be decomposed into bound and free: *R*^*t*^ = *R*^*b*^ + *R*^*f*^. Moreover, the number of bound ribosomes always equals the total number of bound tRNAs of all species: 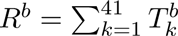.

Initiation and elongation events in the cell occur at rates that are determined by the current state of system (the number of free ribosomes, and the locations of all bound ribosomes) and by the underlying physical parameters of the cell. The underlying physical parameters are simply the volume of the cell, and the characteristic lengths and diffusion constants of ribosomes and tRNA molecules. The time between subsequent events are exponentially distributed, and Monte Carlo simulations proceed by incrementing time according to exponential deviates and re-computing rates of subsequent events (Gillespie, 1977).

### 1.1 Diffusion of ribosomes and tRNAs

In a cell of fixed volume, the average time required for any given molecule to move to one position, known as the characteristic time *τ*, is given by

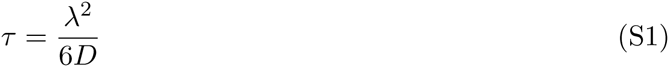

where *D* is the diffusion coefficient of the molecule and *λ* is its characteristic length. The characteristic times of tRNAs and ribosomes are *τ*_*t*_ = 4.45 × 10^-7^ s and *τ*_*r*_ = 5 × 10^-4^ s, respectively (Shah et al., 2013).

### 1.2 Translation initiation rates

Let *ρ*_*i*_ be the initiation rate at an mRNA of gene *i*. The rate *ρ*_*i*_ is set to zero if any of the first 10 codons of the mRNA is currently bound by a ribosome. Otherwise, the rate is

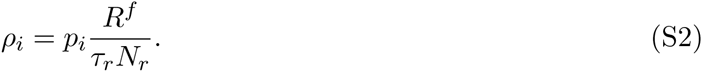

The term 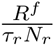 in this equation denotes the rate at which free ribosomes (*R*^*f*^) diffuse to a given mRNA molecule. And the term *p*_*i*_ denotes the probability with which a ribosome will actually initiate translation of an mRNA molecule, once it has diffused to its 5’ end. The parameters *p*_*i*_ allow us to account for sequence-specific variation in initiation rates among genes.

### 1.3 Translation elongation rates

Consider a ribosome bound at codon of type *j* at position *k* on an mRNA. Its elongation rate is set to zero if any of the following *k* + 10 codons of the mRNA are currently occupied by another ribosome. Otherwise, the elongation rate depends on the number of free cognate tRNAs for that codon 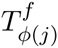 and the wobble parameter associated with the tRNA-codon pair *w*_*j*_. If there is a perfect match between the tRNA and the codon, then *w*_*j*_ = 1. Else *w*_*ry*/*yr*_ = 0.64 if the mismatch is due to a purine-pyrimidine wobble or *w*_*rr*/*yy*_ = 0.61 if the mismatch is due to purine-purine or pyrimidine-pyrimidine wobble (Curran & Yarus, 1989; Lim & Curran, 2001). The elongation rate is thus given by

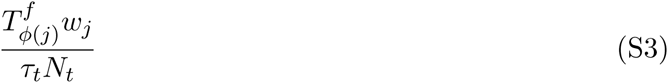

In addition, the time spent by a ribosome in selecting the cognate tRNA depends on the relative abundances of various competing tRNAs as well as organism specific kinetic rates associated with ribosomal proofreading. Using the method described in (Shah et al., 2013) we estimate the average time s spent by the ribosome in kinetic proofreading to select the correct tRNA. As a result, accounting for tRNA competition, the actual elongation rate of a codon is

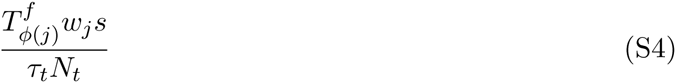

### 1.4 Translation termination

We assume that translation termination is an instantaneous event that occurs immediately after elongation of the last codon at position L. Upon termination the pool of free ribosomes and free tRNAs corresponding to the codon *j*′ at position L - 1 each increases by 1 (*R*^*f*^ → *R*^*f*^ + 1; 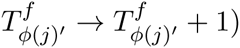.

### 1.5 Estimating initiation probabilities using ribosome-profiling data

We use analytical approximation of the whole-cell simulation model (Shah et al., 2013) described above to estimate the gene-specific probability of translation initiation once a free ribosome reaches the 5’ end of an mRNA. The gene-specific initiation probability *p*_*i*_ is given by Eqn. 27 in (Shah et al., 2013) as follows:

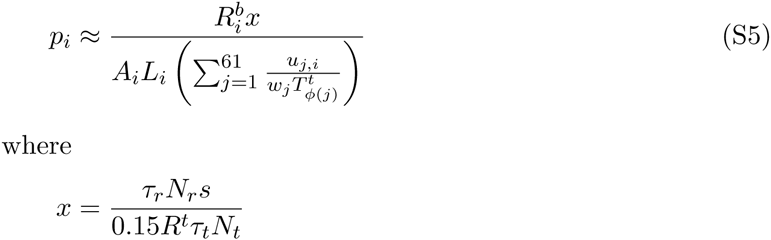

The term *x* depends on the bio-physical parameters of tRNAs, ribosomes and volume of an yeast cell, whose values are described above. The term 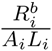 describes the density of ribosome on an mRNA of gene *i* and is equivalent to the translation efficiency (TE) described as:

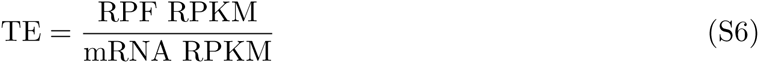

Thus we estimate the gene-specific initiation probability from ribosome profiling data by plugging in the experimentally determined TE ratios in Eqn. S5.

## 2 Elongation arrest by cycloheximide

In ribosome profiling experiments, ribosomes are often stabilized by addition of chemicals that arrest elongation to prevent run-offs during sample preparation. Cycloheximide (CHX) is usually the preferred elongation inhibitor (Ingolia et al., 2009; Zinshteyn & Gilbert, 2013; Brar et al., 2012; Gerashchenko et al., 2012; Artieri & Fraser, 2014; McManus et al., 2014). However, it is unclear whether addition of CHX biases estimates of ribosome densities on mRNAs and hence subsequently affects inferences of protein translation dynamics based on these estimates. To explore how addition of CHX affects ribosome densities, we simulate protein translation in a cell by modeling the action of CHX.

CHX arrests ribosomes on mRNAs by binding with a tRNA in the E-site of the ribosome (Schneider-Poetsch et al., 2010). However, the E-site of a ribosome is almost always empty except for a short period immediately following an elongation and translocation event (Chen et al., 2011). As a result, upon addition of CHX to the cell, a recently elongated ribosome becomes a potential target for CHX. We model the action of CHX by assuming that whenever a ribosome elongates a codon, there is a constant probability with which CHX binds and arrests it. Assuming that the binding of CHX is a reversible process, we model CHX dissociation with a constant rate per bound CHX.

We begin by first simulating protein translation in a normal cell yeast till it reaches equilibrium (1500 sec). After this, whenever a ribosome at codon positions *k* of an mRNA elongates to *k*+1, the ribosome is arrested at *k* + 1 by CHX with constant probability *p*_*chx*-*on*_. CHX dissociates from a bound ribosome at with a constant rate *r*_*chx*-*off*_. We vary the probability of CHX-binding *p*_*chx*-*on*_ and its dissociation rate *r*_*chx*-*off*_ to understand its effects on ribosome densities and dynamics of protein translation. We find that when CHX acts rapidly and has a low dissociation rate (*p*_*chx*-*on*_ = 0.2, *r*_*chx*-*off*_ = 0.01), we see peak excess relative ribosome-footprint density (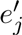 Eqn S10) following the first ten codons is *∼* 400%, which is to comparable to ramps observed in ribosome profiling experiments with CHX pre-treatment (Figure S2A,B) but significantly higher than the ramp observed without CHX in both simulations (*∼* 20%) and in the current study (*∼* 60%). This suggests that a large ramp of ribosome densities in the 5’ region is likely an artifact of the action of CHX (Gerashchenko & Gladyshev, 2014).

## 3 Mapping ribosome profiling reads

The *S. cerevisiae* reference genome sequence and transcript models were downloaded from Ensembl at ftp://ftp.ensembl.org/pub/release-74/fasta/saccharomyces_cerevisiae/dna/ and ftp://ftp.ensembl.org/pub/release-74/gtf/saccharomyces_cerevisiae/.

Data was processed using a framework written in Python. Reads were trimmed from the right of adapter sequences according to the specific library preparations used to generate each data set: reads from poly-adenlyated libraries were trimmed of all trailing As, and reads from libraries prepared with a pre-adynlyated linker (either ‘CTGTAGGCACCATCAAT’ or ‘TCGTATGCCGTCTTCTGCTTG’) were trimmed to the first position from the left at which the next 10 bases in the read were within a hamming distance of 1 from the first 10 bases of the linker sequence or to where a suffix of the read exactly matched a prefix of the linker sequence. For our data, 8 nt of randomized barcode sequence was trimmed from the left of each read and appended to the read’s name. Reads originating from ribosomal RNAs were pre-filtered by mapping to an index of yeast rRNA sequences with bowtie2 version 2.2.1. Filtered reads were then mapped to the yeast genome and spliced transcript models using tophat2 v2.0.9. Reads mapping to any tRNA or other noncoding RNA genes were discarded. For each annotated coding sequence, counts of the number of uniquely mapped reads on the sense strand whose 5’-most mapped base occupied every position from 50 nt upstream of the start codon to 50 nt downstream of the stop codon were calculated. To calculate codon occupancies, only trimmed reads of length 28, 29 and 30 (for which the identity of the codon occupying the A-site of the ribosome could be most reliably inferred) were used. Reads of length 28 and 29 were assigned to the codon at position +14, 15, or 16 from the start of the read, and reads of length 30 were assigned to the codon at position +15, 16, or 17. To calculate read densities, reads of all lengths were included.

Data sources: (GSE* indicates GEO accession number)

Flash-freeze: GSE53313

Ingolia (Science): GSE13750 (Ingolia et al., 2009)

Zinshteyn: GSE45366 (Zinshteyn & Gilbert, 2013)

Gerashchenko: personal communication with Maxim Gerashchenko (Gerashchenko et al., 2012)

Artieri: GSE50049 (Artieri & Fraser, 2014)

Mcmanus: GSE52119 (McManus et al., 2014)

Guydosh: GSE52968 (Guydosh & Green, 2014)

## 4 Metagene analyses of ribosome and mRNA densities

To understand how ribosome and mRNA densities vary along the length of a transcript, we estimate position-specific ribosome densities of individual genes into a composite metagene. Let *x*_*i,j*_ be the number of mapped RPF reads to position *j* of gene *i* based on its A-site. In order to avoid biases induced due genes with low coverage of reads, we restricted our analyses to genes with at least 128 mapped total mapped reads. To account for differences in initiation rates between different genes, we calculate the normalized ribosome density *z*_*i,j*_ at codon position *j* by normalizing the mapped reads at that codon by the mean number of mapped reads in that gene.

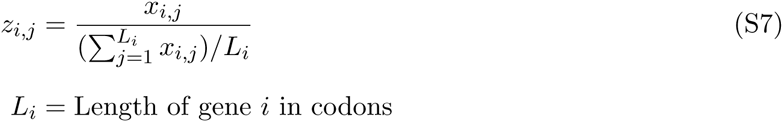

We calculate the excess ribosome densities *e*_*j*_ at a particular position *j* by averaging the normalized ribosome density *z*_*i,j*_ across all genes whose length is at least *j* (*L*_*i*_ <= *j*).

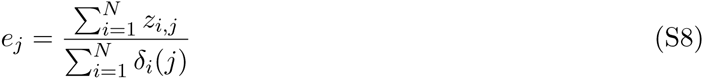

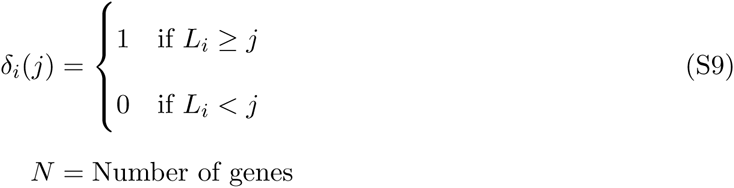

The amount of excess ribosome densities *e*_*j*_ at the 5’ ends of genes vary with each dataset. As a result, the excess ribosome densities asymptote at different values depending on the dataset, making it harder to compare the peaks of ribosome densities across datasets. To account for these differences, we estimated relative excess densities 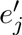 by normalizing excess ribosome-densities *e*_*j*_ with excess ribosome densities in the region spanning 450 and 500 codons. This region was chosen based on the observation that excess ribosome densities in all ribosome profiling dataset reach an asymptote around 450-500 codons. The relative excess ribosome densities 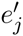 were calculated as follows:

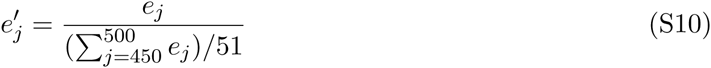

We report peak excess ribosome densities as the maximum of 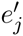 in a region spanning 10 to 500 codons. We ignore the first few codons as they are highly variable and result in sharp peaks due to continued initiation events.

We calculate excess mRNA reads at codon position *j* similar to excess RPF reads described above. Let *y*_*i,j*_ be the number of mapped mRNA reads to position *j* of gene *i*. The normalized mRNA density at *g*_*i,j*_ at codon position *j* by normalizing the mapped reads at that codon by the mean number of mapped reads in that gene.

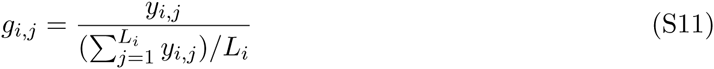

We calculate the excess mRNA densities *h*_*j*_ at a particular position *j* by averaging the normalized mRNA density *g*_*i,j*_ across all genes whose length is at least *j* (*L*_*i*_ <= *j*).

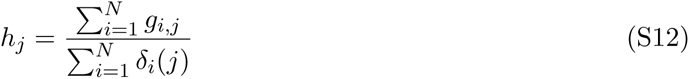

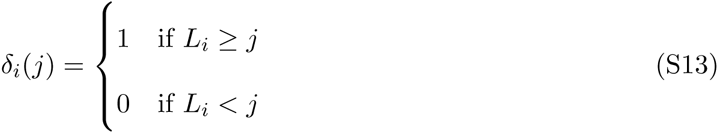

The relative excess mRNA densities 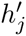 based on 450-500 codons were calculated as follows:

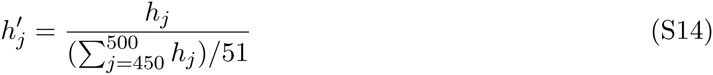

To estimate the degree of bias in mRNA measurements in the 3’ ends of genes, we scale excess mRNA densities at a codon *j* by 450-500 codons from the stop codon as follows:

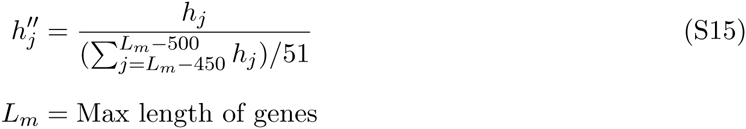

## 5 Comparing 5’ and 3’ codon-specific ribosome densities

To estimate codon-specific ribosome densities in the 5’ and 3’ ends of genes, we begin by first calculating normalized ribosome densities within a gene *z*_*i,j*_ (Eqn. S7) for all genes with at least 250 codons. Normalizing ribosome-densities within a gene removes the effect of differences in initiation rates among genes when comparing normalized reads across many genes. The average ribosome density of all ribosome reads at codon type *k* in the 5’ (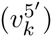) and 3’ (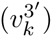) ends across the genome is then given by

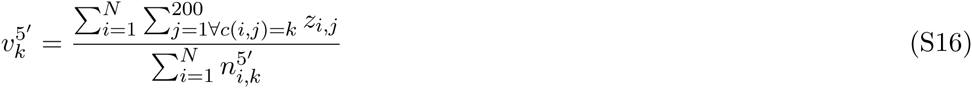

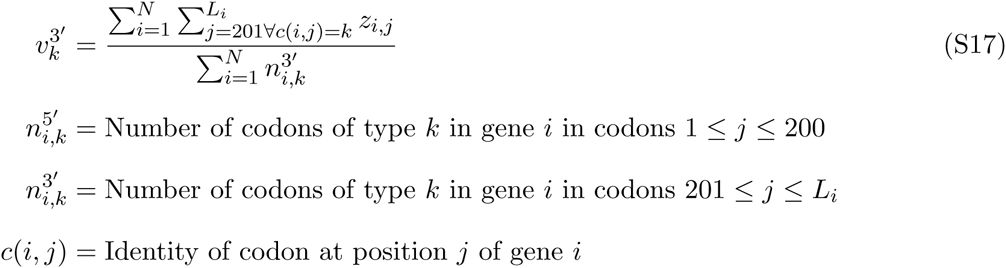

## 6 Estimating codon-specific elongation times

Most theoretical studies of protein translation assume a negative relationship between the elongation time of a codon and its tRNA abundance. To test this, we estimate codon-specific elongation times from ribosome densities as follows:

In order to avoid the confounding effects of 5’ ribosomal ramp on our estimates of codon-specific ribosome densities, we restrict our analyses to genes with at least 250 codons and only consider RPF reads mapped from codon position 200 onwards. Let *x*_*i,j*_ be the number of mapped RPF reads to position *j* of gene *i*. The normalized ribosome density 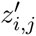 at codon position *j* >= 200 is given by normalizing the mapped reads at that codon by the mean number of mapped reads from codon position 200 to *L*_*i*_.

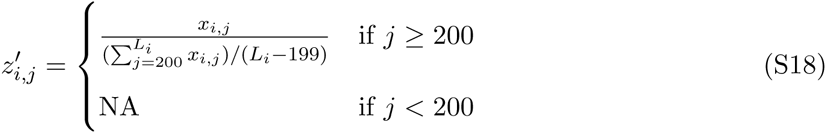

Normalizing ribosome-densities within a gene removes the effect of differences in initiation rates among genes when comparing normalized reads across many genes. The average ribosome density of all ribosome reads at codon type *k* (*v*_*k*_) across the genome is then given by

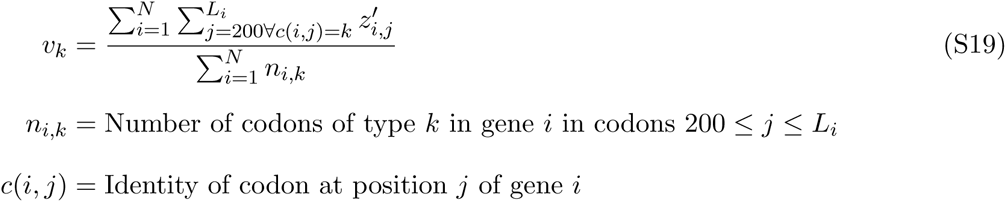

The expected elongation time of a codon is then directly proportional to average codon-specific ribosome density, *v*_*k*_ because codons with longer elongation times have higher average ribosome densities.

## 7 Estimating protein synthesis rates

We estimate protein synthesis rates of individual genes using the densities of ribosomes on their mRNAs. However, the ribosome densities per mRNA as estimated by taking ratios of RPF RPKM and mRNA RPKM are likely biased. The main source of this bias is the presence of a 5’ ramp of RPF reads that varies with position along a transcript.

### 7.1 Ramp correction factor

In order to obtain unbiased estimates of gene-specific ribosome-densities, we apply a position-dependent correction factor to RPF reads that accounts for the ramp. However, the observed ramp of ribosome densities is partly a result of codon ordering within genes in addition to experimental artifacts. Our position-dependent correction factor accounts for the ribosomal ramp due to experimental artifacts by explicitly taking into account the contribution of codon usage dependent ramp.

The total excess ribosome density at a position *j* across all gene is given by *e*_*j*_ (see above, Eqn. S8). To calculate the expected excess ribosome density at a position *j* due to codon ordering within a gene, we first calculate average ribosome density for codon *k*, *v*_*k*_ as described above (Eqn. S19). The expected excess ribosome density *d*_*j*_ at position *j* due to patterns of codon usage is given as follows:

The relative codon-usage expected ribosome density *q*_*i,j*_ at position *j* of gene *i* is given by normalizing the expected ribosome density *v*_*c*(*i,j*)_ at that codon by the mean expected ribosome density for that gene.

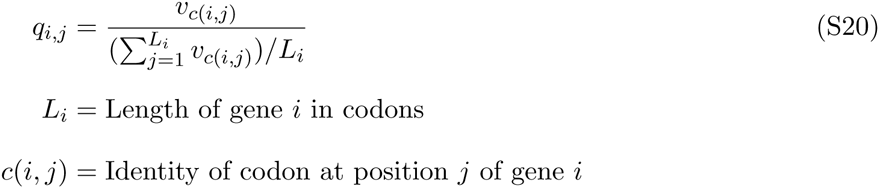

The average codon-usage expected ribosome density at position *j* (*d*_*j*_) across the genome is then given by

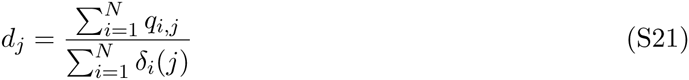

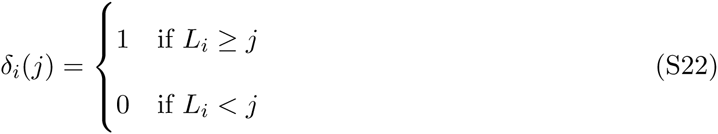

The ramp correction factor *f*_*j*_ at position *j* along any gene is then defined as the ratio of observed ramp *e*_*j*_ over the expected ramp *d*_*j*_.

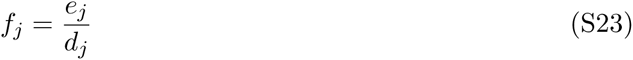

### 7.2 Unbiased estimate of ribosome density per mRNA

Let *x*_*i,j*_ be the number of mapped RPF reads and *y*_*i,j*_ be the number of mapped mRNA reads to position *j* of gene *i*. The unbiased estimate of ribosome density *r*_*i*_ per mRNA for gene *i* is defined as

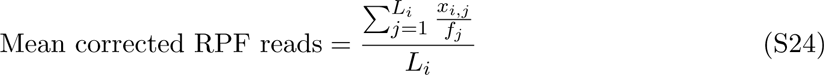

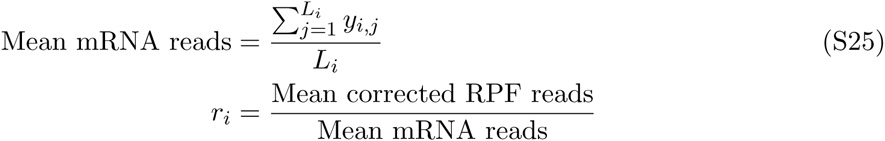

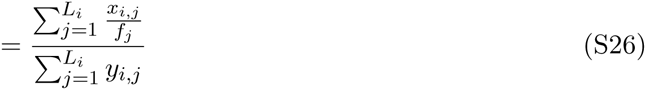

where *f*_*j*_ is the ramp correction factor (see above, Eqn. S23).

### 7.3 Initiation efficiency

We estimate initiation efficiency of a genes using the analytic approximations for the initiation probability *p*_*i*_ based on steady-state behavior of a whole cell simulation described above (Eqn. S5).

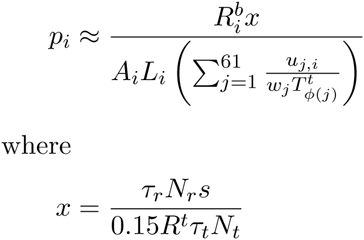

see Methods (1. Simulation model section) above for details on parameter notations and values. 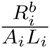 describes the ribosome density per mRNA on gene *i*. Here we substitute 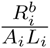 with unbiased estimates of ribosome density per mRNA *r*_*i*_ (Eqn. S26) in Eqn. S5. Moreover, estimates of codon-specific ribosome densities *v*_*k*_ (Eqn. S19) reflect average elongation times of codons – codons with longer elongation times have higher ribosome densities. Therefore, we substitute expected elongation time of a codon given by 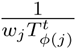 in Eqn. S5 with estimates of codon-specific ribosome densities *v*_*k*_. As a result, our initiation efficiency 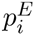 for gene *i* is estimated solely from ribosome profiling data and is defined as

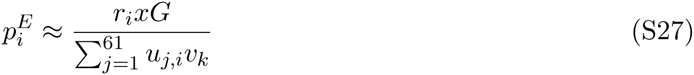

where *G* is the global scaling parameter, which scales *r*_*i*_ such that the total number of ribosomes within a cell are 200, 000, total number of mRNAs are 60, 000 and total number of tRNAs are 3, 300, 000 based on empirical estimates (Shah et al., 2013).

### 7.4 Protein synthesis rates

Protein synthesis rate *S*_*i*_ of a gene *i* within a cell depend on total number of mRNAs for that gene (*A*_*i*_) and initiation rate per mRNA (*ρ*_*i*_ Eqn. S2) and the number of free ribosomes. Here we modify Eqn. S2) by substituting initiation probabilities (*p*_*i*_) with estimates of initiation efficiencies (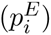) based on profiling data (Eqn. S27).

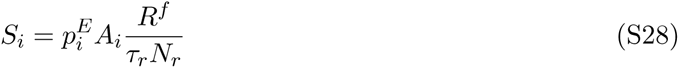

In estimating *S*_*i*_, we assume that 15% of the total 200,000 ribosomes are free and the rest are bound to mRNAs, such that *R*^*f*^ = 3 × 10^4^. Furthermore, we scale mRNA abundances as measured by mRNA RPKM such that the total number of mRNAs are 60,000 (∑_*i*_ *A*_*i*_ = 6 × 10^4^).

## 8 Co-translational folding and inter-domain linkers

During protein translation, a growing polypeptide chain begins to fold as soon as it emerges from the ribosome - a process known as co-translational folding. Several studies have suggested that pausing of ribosome at specific instances is necessary for the nascent polypeptide to take native-like folds (Kimchi-Sarfaty et al., 2007; Pechmann & Frydman, 2013). If ribosomal pausing significantly affects co-translation folding, then we expect a higher density of ribosomes in regions between protein domains. To test this, we downloaded domain assignments for individual genes in the *S. cerevisiae* genome from SGD – http://downloads.yeastgenome.org/curation/calculated_protein_info/domains/domains.tab based on InterProScan http://www.ebi.ac.uk/interpro on Jan 14, 2014. Domain assignments were based on InterProScan classifications (Jones et al., 2014) obtained from the Superfamily database (Wilson et al., 2009) and Pfam database.

## 9 Regression model

Initiation efficiency (*p*^*E*^) of a gene depends on several features of a coding sequence. In order to identify a set of features that explain the most variation in *p*^*E*^, we use the multiple regression framework in R (R Core Team, 2012). We regressed the *p*^*E*^ of a gene against its length, mRNA abundance (RPKM), 5’ UTR length and its GC content, 5’ cap folding energy, and number of upstream ATGs in 5’ UTR (uATG). We restricted the analyses to 2549 genes with experimentally verified 5’ UTR lengths and number of upstream ATGs (Arribere & Gilbert, 2013). To estimate 5’ cap folding energy we used sequences of length 70 nts from the 5’ end of the mRNA transcript as sequences of these lengths showed the highest correlation with TE ratios (Figure S14). We calculated the folding energies using RNAfold algorithm from Vienna RNA package (Hofacker et al., 1994) at 37°C. The values of *p*^*E*^, mRNA RPKM and protein length were log-transformed in the regression model.

To identify which features explain the highest amount of variation in initiation efficiencies, we used Akaike’s Information Criteria (AIC) for model selection. We performed both step-up and step-down model selection using the stepAIC function in MASS package in R. We find that the multiple regression model that best explains the variation in *p*^*E*^ even after penalizing for model complexity includes all the 6 variables considered (Table S3). This model explains *∼* 58% of variation in *p*^*E*^ across all the genes considered. We find that initiation efficiencies scale positively with predicted folding energies and mRNA abundances. This indicates that genes with weaker 5’ cap structure and RNA structure around the start site have higher rates of initiation. In contrast, genes with a higher number of uATGs and longer 5’ UTRs and coding sequence length have lower *p*^*E*^. Moreover, genes that have higher mRNA abundances also tend to have higher *p*^*E*^ and hence higher ribosome densities (TE) on them. This suggests that genes under selection for higher protein abundances are selected both at the level of transcription – leading to higher mRNA abundances and at the translation level – leading to higher ribosome densities.

## References

Amrani, N., Ghosh, S., Mangus, D.A., and Jacobson, A. (2008). Translation factors promote the formation of two states of the closed-loop mRNP. Nature 453, 1276–1280.

Andersson, S.G., and Kurland, C.G. (1990). Codon preferences in free-living microorganisms. Microbiological reviews 54, 198–210.

Arava, Y., Wang, Y., Storey, J.D., Liu, C.L., Brown, P.O., and Herschlag, D. (2003). Genome-wide analysis of mRNA translation profiles in Saccharomyces cerevisiae. Proceedings of the National Academy of Sciences of the United States of America 100, 3889–3894.

Arribere, J.A., and Gilbert, W.V. (2013). Roles for transcript leaders in translation and mRNA decay revealed by transcript leader sequencing. Genome research 23, 977–987.

Artieri, C.G., and Fraser, H.B. (2014a). Accounting for biases in riboprofiling data indicates a major role for proline in stalling translation. Genome research 24, 2011–2021.

Artieri, C.G., and Fraser, H.B. (2014b). Evolution at two levels of gene expression in yeast. Genome research 24, 411–421.

Ashe, M.P., De Long, S.K., and Sachs, A.B. (2000). Glucose depletion rapidly inhibits translation initiation in yeast. Molecular biology of the cell 11, 833–848.

Bateman, A., Birney, E., Cerruti, L., Durbin, R., Etwiller, L., Eddy, S.R., Griffiths-Jones, S., Howe, K.L., Marshall, M., and Sonnhammer, E.L. (2002). The Pfam protein families database. Nucleic acids research 30, 276–280.

Bennetzen, J.L., and Hall, B.D. (1982). Codon selection in yeast. The Journal of biological chemistry 257, 3026–3031.

Beznoskova, P., Cuchalova, L., Wagner, S., Shoemaker, C.J., Gunisova, S., von der Haar, T., and Valasek, L.S. (2013). Translation initiation factors eIF3 and HCR1 control translation termination and stop codon read-through in yeast cells. PLoS genetics 9, e1003962.

Brandman, O., Stewart-Ornstein, J., Wong, D., Larson, A., Williams, C.C., Li, G.W., Zhou, S., King, D., Shen, P.S., Weibezahn, J., et al. (2012). A ribosome-bound quality control complex triggers degradation of nascent peptides and signals translation stress. Cell 151, 1042–1054.

Brar, G.A., Yassour, M., Friedman, N., Regev, A., Ingolia, N.T., and Weissman, J.S. (2012). High-resolution view of the yeast meiotic program revealed by ribosome profiling. Science 335, 552–557.

Bulmer, M. (1991). The selection-mutation-drift theory of synonymous codon usage. Genetics 129, 897–907.

Charneski, C.A., and Hurst, L.D. (2013). Positively charged residues are the major determinants of ribosomal velocity. PLoS biology 11, e1001508.

Christensen, A.K., Kahn, L.E., and Bourne, C.M. (1987). Circular polysomes predominate on the rough endoplasmic reticulum of somatotropes and mammotropes in the rat anterior pituitary. The American journal of anatomy 178, 1–10.

Clarkson, B.K., Gilbert, W.V., and Doudna, J.A. (2010). Functional overlap between eIF4G isoforms in Saccharomyces cerevisiae. PLoS one 5, e9114.

Cox, J.S., and Walter, P. (1996). A novel mechanism for regulating activity of a transcription factor that controls the unfolded protein response. Cell 87, 391–404.

Crombie, T., Swaffield, J.C., and Brown, A.J. (1992). Protein folding within the cell is influenced by controlled rates of polypeptide elongation. Journal of molecular biology 228, 7–12.

Csárdi G, Franks A, Choi DS, Airoldi EM, Drummond DA. (2015). Accounting for experimental noise reveals that mRNA levels, amplified by post-transcriptional processes, largely determine steady-state protein levels in yeast. PLoS Genet 11, e1005206.

de Godoy, L.M., Olsen, J.V., Cox, J., Nielsen, M.L., Hubner, N.C., Frohlich, F., Walther, T.C., and Mann, M. (2008). Comprehensive mass-spectrometry-based proteome quantification of haploid versus diploid yeast. Nature 455, 1251–1254.

Dever, T.E., Feng, L., Wek, R.C., Cigan, A.M., Donahue, T.F., and Hinnebusch, A.G. (1992). Phosphorylation of initiation factor 2 alpha by protein kinase GCN2 mediates gene-specific translational control of GCN4 in yeast. Cell 68, 585–596.

Dunn, J.G., Foo, C.K., Belletier, N.G., Gavis, E.R., and Weissman, J.S. (2013). Ribosome profiling reveals pervasive and regulated stop codon readthrough in Drosophila melanogaster. eLife 2, e01179.

Eichhorn, S.W., Guo, H., McGeary, S.E., Rodriguez-Mias, R.A., Shin, C., Baek, D., Hsu, S.H., Ghoshal, K., Villen, J., and Bartel, D.P. (2014). mRNA Destabilization Is the Dominant Effect of Mammalian MicroRNAs by the Time Substantial Repression Ensues. Molecular cell 56, 104–115.

Gardin, J., Yeasmin, R., Yurovsky, A., Cai, Y., Skiena, S., and Futcher, B. (2014). Measurement of average decoding rates of the 61 sense codons in vivo. eLife 3.

Gerashchenko, M.V., and Gladyshev, V.N. (2014). Translation inhibitors cause abnormalities in ribosome profiling experiments. Nucleic acids research.

Gerashchenko, M.V., Lobanov, A.V., and Gladyshev, V.N. (2012). Genome-wide ribosome profiling reveals complex translational regulation in response to oxidative stress. Proceedings of the National Academy of Sciences of the United States of America 109, 17394–17399.

Godefroy-Colburn, T., Ravelonandro, M., and Pinck, L. (1985). Cap accessibility correlates with the initiation efficiency of alfalfa mosaic virus RNAs. European journal of biochemistry / FEBS 147, 549–552.

Gold, L. (1988). Posttranscriptional regulatory mechanisms in Escherichia coli. Annual review of biochemistry 57, 199–233.

Gouy, M., and Gautier, C. (1982). Codon usage in bacteria: correlation with gene expressivity. Nucleic acids research 10, 7055–7074.

Guo, H., Ingolia, N.T., Weissman, J.S., and Bartel, D.P. (2010). Mammalian microRNAs predominantly act to decrease target mRNA levels. Nature 466, 835–840.

Guydosh, N.R., and Green, R. (2014). Dom34 rescues ribosomes in 3’ untranslated regions. Cell 156, 950–962.

Hope, I.A., and Struhl, K. (1985). GCN4 protein, synthesized in vitro, binds HIS3 regulatory sequences: implications for general control of amino acid biosynthetic genes in yeast. Cell 43, 177–188.

Hosoda, N., Kobayashi, T., Uchida, N., Funakoshi, Y., Kikuchi, Y., Hoshino, S., and Katada, T. (2003). Translation termination factor eRF3 mediates mRNA decay through the regulation of deadenylation. The Journal of biological chemistry 278, 38287–38291.

Hsieh, A.C., Liu, Y., Edlind, M.P., Ingolia, N.T., Janes, M.R., Sher, A., Shi, E.Y., Stumpf, C.R., Christensen, C., Bonham, M.J., et al. (2012). The translational landscape of mTOR signalling steers cancer initiation and metastasis. Nature 485, 55–61.

Hu, W., Sweet, T.J., Chamnongpol, S., Baker, K.E., and Coller, J. (2009). Cotranslational mRNA decay in Saccharomyces cerevisiae. Nature 461, 225–229.

Hunt, R.T., Hunter, A.R., and Munro, A.J. (1968). Control of haemoglobin synthesis: a difference in the size of the polysomes making alpha and beta chains. Nature 220, 481–483.

Ikemura, T. (1981). Correlation between the abundance of Escherichia coli transfer RNAs and the occurrence of the respective codons in its protein genes: a proposal for a synonymous codon choice that is optimal for the E. coli translational system. Journal of molecular biology 151, 389–409.

Ikemura, T. (1985). Codon usage and tRNA content in unicellular and multicellular organisms. Molecular biology and evolution 2, 13–34.

Ingolia, N.T. (2014). Ribosome profiling: new views of translation, from single codons to genome scale. Nature reviews Genetics 15, 205–213.

Ingolia, N.T., Brar, G.A., Rouskin, S., McGeachy, A.M., and Weissman, J.S. (2012). The ribosome profiling strategy for monitoring translation in vivo by deep sequencing of ribosome-protected mRNA fragments. Nature protocols 7, 1534–1550.

Ingolia, N.T., Ghaemmaghami, S., Newman, J.R., and Weissman, J.S. (2009). Genomewide analysis in vivo of translation with nucleotide resolution using ribosome profiling. Science 324, 218–223.

Ingolia, N.T., Lareau, L.F., and Weissman, J.S. (2011). Ribosome profiling of mouse embryonic stem cells reveals the complexity and dynamics of mammalian proteomes. Cell 147, 789–802.

Jayaprakash, A.D., Jabado, O., Brown, B.D., and Sachidanandam, R. (2011). Identification and remediation of biases in the activity of RNA ligases in small-RNA deep sequencing. Nucleic acids research 39, e141.

Jones, P., Binns, D., Chang, H.Y., Fraser, M., Li, W., McAnulla, C., McWilliam, H., Maslen, J., Mitchell, A., Nuka, G., et al. (2014). InterProScan 5: genome-scale protein function classification. Bioinformatics 30, 1236–1240.

Kawahara, T., Yanagi, H., Yura, T., and Mori, K. (1997). Endoplasmic reticulum stressinduced mRNA splicing permits synthesis of transcription factor Hac1p/Ern4p that activates the unfolded protein response. Molecular biology of the cell 8, 1845–1862.

Kertesz, M., Wan, Y., Mazor, E., Rinn, J.L., Nutter, R.C., Chang, H.Y., and Segal, E. (2010). Genome-wide measurement of RNA secondary structure in yeast. Nature 467, 103–107.

Kimchi-Sarfaty, C., Oh, J.M., Kim, I.W., Sauna, Z.E., Calcagno, A.M., Ambudkar, S.V., and Gottesman, M.M. (2007). A “silent” polymorphism in the MDR1 gene changes substrate specificity. Science 315, 525–528.

Kozak, M. (1984). Selection of initiation sites by eucaryotic ribosomes: effect of inserting AUG triplets upstream from the coding sequence for preproinsulin. Nucleic acids research 12, 3873–3893.

Kozak, M. (1986a). Influences of mRNA secondary structure on initiation by eukaryotic ribosomes. Proceedings of the National Academy of Sciences of the United States of America 83, 2850–2854.

Kozak, M. (1986b). Point mutations define a sequence flanking the AUG initiator codon that modulates translation by eukaryotic ribosomes. Cell 44, 283–292.

Kudla, G., Murray, A.W., Tollervey, D., and Plotkin, J.B. (2009). Coding-sequence determinants of gene expression in Escherichia coli. Science 324, 255–258.

Lareau, L.F., Hite, D.H., Hogan, G.J., and Brown, P.O. (2014). Distinct stages of the translation elongation cycle revealed by sequencing ribosome-protected mRNA fragments. eLife 3, e01257.

Letzring, D.P., Dean, K.M., and Grayhack, E.J. (2010). Control of translation efficiency in yeast by codon-anticodon interactions. Rna 16, 2516–2528.

Li, G.W., Burkhardt, D., Gross, C., and Weissman, J.S. (2014a). Quantifying absolute protein synthesis rates reveals principles underlying allocation of cellular resources. Cell 157, 624–635.

Li, G.W., Oh, E., and Weissman, J.S. (2012). The anti-Shine-Dalgarno sequence drives translational pausing and codon choice in bacteria. Nature 484, 538–541.

Li, J.J., Bickel, P.J., and Biggin, M.D. (2014b). System wide analyses have underestimated protein abundances and the importance of transcription in mammals. PeerJ 2, e270.

Lu, J., and Deutsch, C. (2008). Electrostatics in the ribosomal tunnel modulate chain elongation rates. Journal of molecular biology 384, 73–86.

McManus, C.J., May, G.E., Spealman, P., and Shteyman, A. (2014). Ribosome profiling reveals post-transcriptional buffering of divergent gene expression in yeast. Genome research 24, 422–430.

Mueller, P.P., and Hinnebusch, A.G. (1986). Multiple upstream AUG codons mediate translational control of GCN4. Cell 45, 201–207.

Nagalakshmi, U., Wang, Z., Waern, K., Shou, C., Raha, D., Gerstein, M., and Snyder, M. (2008). The transcriptional landscape of the yeast genome defined by RNA sequencing. Science 320, 1344–1349.

Ouyang, Z., Snyder, M.P., and Chang, H.Y. (2013). SeqFold: genome-scale reconstruction of RNA secondary structure integrating high-throughput sequencing data. Genome research 23, 377–387.

Park, E.H., Zhang, F., Warringer, J., Sunnerhagen, P., and Hinnebusch, A.G. (2011). Depletion of eIF4G from yeast cells narrows the range of translational efficiencies genome-wide. BMC genomics 12, 68.

Pechmann, S., and Frydman, J. (2013). Evolutionary conservation of codon optimality reveals hidden signatures of cotranslational folding. Nature structural & molecular biology 20, 237–243.

Plotkin, J.B., and Kudla, G. (2011). Synonymous but not the same: the causes and consequences of codon bias. Nature reviews Genetics 12, 32–42.

Qian, W., Yang, J.R., Pearson, N.M., Maclean, C., and Zhang, J. (2012). Balanced codon usage optimizes eukaryotic translational efficiency. PLoS genetics 8, e1002603.

Robbins-Pianka, A., Rice, M.D., and Weir, M.P. (2010). The mRNA landscape at yeast translation initiation sites. Bioinformatics 26, 2651–2655.

Rojas-Duran, M.F., and Gilbert, W.V. (2012). Alternative transcription start site selection leads to large differences in translation activity in yeast. Rna 18, 2299–2305.

Rouskin, S., Zubradt, M., Washietl, S., Kellis, M., and Weissman, J.S. (2014). Genomewide probing of RNA structure reveals active unfolding of mRNA structures in vivo. Nature 505, 701–705.

Ruegsegger, U., Leber, J.H., and Walter, P. (2001). Block of HAC1 mRNA translation by long-range base pairing is released by cytoplasmic splicing upon induction of the unfolded protein response. Cell 107, 103–114.

Schneider-Poetsch, T., Ju, J., Eyler, D.E., Dang, Y., Bhat, S., Merrick, W.C., Green, R., Shen, B., and Liu, J.O. (2010). Inhibition of eukaryotic translation elongation by cycloheximide and lactimidomycin. Nature chemical biology 6, 209–217.

Schwanhausser, B., Busse, D., Li, N., Dittmar, G., Schuchhardt, J., Wolf, J., Chen, W., and Selbach, M. (2011). Global quantification of mammalian gene expression control. Nature 473, 337–342.

Shah, P., Ding, Y., Niemczyk, M., Kudla, G., and Plotkin, J.B. (2013). Rate-limiting steps in yeast protein translation. Cell 153, 1589–1601.

Shah, P., and Gilchrist, M.A. (2011). Explaining complex codon usage patterns with selection for translational efficiency, mutation bias, and genetic drift. Proceedings of the National Academy of Sciences of the United States of America 108, 10231–10236.

Sharp, P.M., and Li, W.H. (1987). The codon Adaptation Index--a measure of directional synonymous codon usage bias, and its potential applications. Nucleic acids research 15, 1281–1295.

Siridechadilok, B., Fraser, C.S., Hall, R.J., Doudna, J.A., and Nogales, E. (2005). Structural roles for human translation factor eIF3 in initiation of protein synthesis. Science 310, 1513–1515.

Sorefan, K., Pais, H., Hall, A.E., Kozomara, A., Griffiths-Jones, S., Moulton, V., and Dalmay, T. (2012). Reducing ligation bias of small RNAs in libraries for next generation sequencing. Silence 3, 4.

Sorensen, M.A., and Pedersen, S. (1991). Absolute in vivo translation rates of individual codons in Escherichia coli. The two glutamic acid codons GAA and GAG are translated with a threefold difference in rate. Journal of molecular biology 222, 265–280.

Subtelny, A.O., Eichhorn, S.W., Chen, G.R., Sive, H., and Bartel, D.P. (2014). Poly(A)tail profiling reveals an embryonic switch in translational control. Nature 508, 66–71.

Szamecz, B., Rutkai, E., Cuchalova, L., Munzarova, V., Herrmannova, A., Nielsen, K.H., Burela, L., Hinnebusch, A.G., and Valasek, L. (2008). eIF3a cooperates with sequences 5’ of uORF1 to promote resumption of scanning by post-termination ribosomes for reinitiation on GCN4 mRNA. Genes & development 22, 2414–2425.

Tarun, S.Z., Jr., and Sachs, A.B. (1996). Association of the yeast poly(A) tail binding protein with translation initiation factor eIF-4G. The EMBO journal 15, 7168–7177.

Thanaraj, T.A., and Argos, P. (1996). Ribosome-mediated translational pause and protein domain organization. Protein science : a publication of the Protein Society 5, 1594–1612.

Thoreen, C.C., Chantranupong, L., Keys, H.R., Wang, T., Gray, N.S., and Sabatini, D.M. (2012). A unifying model for mTORC1-mediated regulation of mRNA translation. Nature 485, 109–113.

Tuller, T., Carmi, A., Vestsigian, K., Navon, S., Dorfan, Y., Zaborske, J., Pan, T., Dahan, O., Furman, I., and Pilpel, Y. (2010). An evolutionarily conserved mechanism for controlling the efficiency of protein translation. Cell 141, 344–354.

Varenne, S., Buc, J., Lloubes, R., and Lazdunski, C. (1984). Translation is a non-uniform process. Effect of tRNA availability on the rate of elongation of nascent polypeptide chains. Journal of molecular biology 180, 549–576.

Wells, S.E., Hillner, P.E., Vale, R.D., and Sachs, A.B. (1998). Circularization of mRNA by eukaryotic translation initiation factors. Molecular cell 2, 135–140.

Wilson, D., Pethica, R., Zhou, Y., Talbot, C., Vogel, C., Madera, M., Chothia, C., and Gough, J. (2009). SUPERFAMILY--sophisticated comparative genomics, data mining, visualization and phylogeny. Nucleic acids research 37, D380–386.

Zhang, D., and Shan, S.O. (2012). Translation elongation regulates substrate selection by the signal recognition particle. The Journal of biological chemistry 287, 7652–7660.

Zhou, J., Liu, W.J., Peng, S.W., Sun, X.Y., and Frazer, I. (1999). Papillomavirus capsid protein expression level depends on the match between codon usage and tRNA availability. Journal of virology 73, 4972–4982.

Zinshteyn, B., and Gilbert, W.V. (2013). Loss of a conserved tRNA anticodon modification perturbs cellular signaling. PLoS genetics 9, e1003675.

## References

Arribere, J. A. & Gilbert, W. V. (2013). Roles for transcript leaders in translation and mRNA decay revealed by transcript leader sequencing. Genome Res, 23(6), 977–987.

Artieri, C. G. & Fraser, H. B. (2014). Evolution at two levels of gene expression in yeast. Genome Res, 24(3), 411–421.

Brar, G. A., Yassour, M., Friedman, N., Regev, A., Ingolia, N. T., & Weissman, J. S. (2012). Highresolution view of the yeast meiotic program revealed by ribosome profiling. Science, 335(6068), 552–557.

Chen, C., Stevens, B., Kaur, J., Smilansky, Z., Cooperman, B. S., & Goldman, Y. E. (2011). Allosteric vs. spontaneous exit-site (E-site) tRNA dissociation early in protein synthesis. Proc Natl Acad Sci USA, 108(41), 16980–16985.

Curran, J. F. & Yarus, M. (1989). Rates of aminoacyl-tRNA selection at 29 sense codons in vivo. J Mol Biol, 209(1), 65–77.

Gerashchenko, M. V. & Gladyshev, V. N. (2014). Translation inhibitors cause abnormalities in ribosome profiling experiments. Nucl Acids Res, 42(17), e134–e134.

Gerashchenko, M. V., Lobanov, A. V., & Gladyshev, V. N. (2012). Genome-wide ribosome profiling reveals complex translational regulation in response to oxidative stress. Proc Natl Acad Sci USA, 109(43), 17394–17399.

Gillespie, D. (1977). Exact stochastic simulation of coupled chemical reactions. The journal of physical chemistry, 81(25), 2340–2361.

Guydosh, N. R. & Green, R. (2014). Dom34 rescues ribosomes in 3’ untranslated regions. Cell, 156(5), 950–962.

Hofacker, I. L., Fontana, W., & Stadler, P. (1994). Fast folding and comparison of RNA secondary structures. Monatshefte für Chemie, 125, 167–188.

Ingolia, N. T., Ghaemmaghami, S., Newman, J. R. S., & Weissman, J. S. (2009). Genome-Wide Analysis in Vivo of Translation with Nucleotide Resolution Using Ribosome Profiling. Science, 324(5924), 218–223.

Jones, P., Binns, D., Chang, H. Y., Fraser, M., Li, W., McAnulla, C., McWilliam, H., Maslen, J., Mitchell, A., Nuka, G., Pesseat, S., Quinn, A. F., Sangrador-Vegas, A., Scheremetjew, M., Yong, S. Y., Lopez, R., & Hunter, S. (2014). InterProScan 5: genome-scale protein function classification. Bioinformatics, 30(9), 1236–1240.

Kimchi-Sarfaty, C., Oh, J. M., Kim, I.-W., Sauna, Z. E., Calcagno, A. M., Ambudkar, S. V., & Gottesman, M. M. (2007). A “silent” polymorphism in the MDR1 gene changes substrate specificity. Science, 315(5811), 525–528.

Lim, V. I. & Curran, J. F. (2001). Analysis of codon:anticodon interactions within the ribosome provides new insights into codon reading and the genetic code structure. RNA, 7(7), 942–957.

McManus, C. J., May, G. E., Spealman, P., & Shteyman, A. (2014). Ribosome profiling reveals post-transcriptional bu?ering of divergent gene expression in yeast. Genome Res, 24(3), 422–430.

Pechmann, S. & Frydman, J. (2013). Evolutionary conservation of codon optimality reveals hidden signatures of cotranslational folding. Nat Struct Mol Biol, 20(2), 237–243.

R Core Team (2012). R: A language and environment for statistical computing. R foundation for Statistical Computing.

Schneider-Poetsch, T., Ju, J., Eyler, D. E., Dang, Y., Bhat, S., Merrick, W. C., Green, R., Shen, B., & Liu, J. O. (2010). Inhibition of eukaryotic translation elongation by cycloheximide and lactimidomycin. Nat Chem Biol, 6(3), 209–217.

Shah, P., Ding, Y., Niemczyk, M., Kudla, G., & Plotkin, J. B. (2013). Rate-limiting steps in yeast protein translation. Cell, 153(7), 1589–1601.

Wilson, D., Pethica, R., Zhou, Y., Talbot, C., Vogel, C., Madera, M., Chothia, C., & Gough, J. (2009). SUPERFAMILY–sophisticated comparative genomics, data mining, visualization and phylogeny. Nucl Acids Res, 37(Database issue), D380–6.

Zinshteyn, B. & Gilbert, W. V. (2013). Loss of a conserved tRNA anticodon modification perturbs cellular signaling. PLoS Genet, 9(8), e1003675.

